# Rodent Automated Bold Improvement of EPI Sequences (RABIES): A standardized image processing and data quality platform for rodent fMRI

**DOI:** 10.1101/2022.08.20.504597

**Authors:** Gabriel Desrosiers-Gregoire, Gabriel A. Devenyi, Joanes Grandjean, M. Mallar Chakravarty

## Abstract

Functional magnetic resonance imaging (fMRI) in rodents holds great potential for advancing our understanding of brain networks. Unlike the human fMRI community, there remains no standardized resource in rodents for image processing, analysis and quality control, posing significant reproducibility limitations. Our software platform, Rodent Automated Bold Improvement of EPI Sequences (RABIES), is a novel pipeline designed to address these limitations for preprocessing, quality control, and confound correction, along with best practices for reproducibility and transparency. We demonstrate the robustness of the preprocessing workflow by validating performance across multiple acquisition sites and both mouse and rat data. Building upon a thorough investigation into data quality metrics across acquisition sites, we introduce guidelines for the quality control of network analysis and offer recommendations for addressing issues. Taken together, the RABIES software will allow the emerging community to adopt reproducible practices and foster progress in translational neuroscience.

## INTRODUCTION

Functional magnetic resonance imaging (fMRI) experiments in rodents have enabled important investigations of brain activity organization (Chuang & Nasrallah, 2017). Several novel avenues are available for probing the biological underpinnings of brain networks by coupling fMRI experiments conducted in model organisms with transcriptomics (Mills et al., 2018), calcium imaging (using wide-field (Lake et al., 2020) or fibre-photometry (Schlegel et al., 2018)), optogenetics (Grandjean et al., 2019; Zerbi et al., 2019), and more. Such applications hold great potential for bridging across biological and functional scales, and in turn, provide a powerful translational tool to address open questions pertaining to behavior, brain development or neuropsychiatric disease.

The study of resting-state networks in rodents is a rapidly expanding neuroscientific discipline. Whereas workflow standardization emerged for human fMRI applications (Ciric et al., 2018; Esteban et al., 2019), the rodent imaging community has yet to deliver agreed-upon methodological standards across various aspects of experimental designs. Recent multi-site studies have demonstrated significant variability in the ability to detect canonical brain networks in mice (Grandjean et al., 2020) and rats (Grandjean et al., 2023). Accordingly, reproducibility concerns are becoming increasingly salient within the rodent community, drawing parallels with the ongoing conversation within the human literature (Ioanas et al., 2021; Mandino et al., 2020). Despite recent successes towards standardization of rodent acquisition protocols (Grandjean et al., 2023), there are two outstanding problems: 1) laboratories primarily rely on custom-made solutions for image processing since no readily-available software has been validated across multiple acquisition sites, and 2) there are no governing standards for data quality assessment for rodent images. The choices across image processing steps, including image alignment to a commonspace, head motion realignment, and appropriate confound correction, can have significant implications for downstream analysis outcomes; the impact of these experimental design choices further emphasizes the need for a standardized state-of-the-art processing workflow in this domain (Esteban et al., 2019). Developing robust image processing workflows for rodent acquisitions presents new challenges due to site-specific field strengths (4.7T, 7T, 9.4T, 11.7T), coil types (room-temperature vs cryogenic) and geometry (i.e. surface vs saddle vs quadrature coil), and acquisition parameters (Grandjean et al., 2020; Ioanas et al., 2021). Although rodent-adapted software pipelines were recently introduced for either mice and rats (Celestine et al., 2020; Diao et al., 2021; Ioanas et al., 2021; Lee et al., 2021), no existing software has been thoroughly validated across acquisition sites and species in terms of preprocessing quality. Additionally, thorough data quality assessment is also essential for comparability between studies. In the human literature, after noting that poor accounting for confounds may lead to false positive results (Power et al., 2012), there has been significant debate regarding how to appropriately account for fMRI confounds during connectivity analyses. In rodents, investigations into confound characterization are emerging (Diao et al., 2021; Zerbi et al., 2015), but in spite of the aforementioned acquisition protocol guidelines (Grandjean et al., 2023), sources of divergence in data quality between datasets remain, and systematic guidelines for determining the influence of confounds are still needed for the community.

In this work, we aim to address the urgent need for standardized workflow and quality control measures in network analyses. We address these shortcomings in two sections: 1) implementing an adaptive image registration workflow offering consistent high-quality outputs across sites and species, and 2) conducting thorough inspection of data quality markers across acquisition protocols to introduce reliable guidelines for quality control in network analysis. Inspired from previous initiatives fostering reproducibility and standardization in humans (Craddock et al., 2013; Esteban et al., 2019), these innovations are integrated through the distribution of an open-source software platform, Rodent Automated Bold Improvement of EPI Sequences (RABIES; publicly available to the community https://github.com/CoBrALab/RABIES and thoroughly documented online https://rabies.readthedocs.io/en/stable/) that provides an integrative solution for preprocessing, confound correction, analysis, together with data diagnostic tools encouraging best practices for quality control.

## RESULTS

### PART 1: A ROBUST IMAGE PROCESSING WORKFLOW FOR REPRODUCIBLE RODENT FMRI

#### Software design in line with best practices for reproducible science and transparency

RABIES is an open-source software largely inspired by fMRIPrep’s design for reproducibility (Esteban et al. 2019). We made three important design choices that follow best practices for reproducibility in neuroimaging research: 1) the input data format requires that users follow the BIDS standards (K. J. Gorgolewski et al., 2016); 2) quality control can be conducted using automatically-generated reports; and 3) the software distribution is handled via containerized versions (https://hub.docker.com/r/gabdesgreg/rabies; executable through Docker or Singularity (Kurtzer et al., 2017)), ensuring reproducible execution across computing environments. Like fMRIprep, RABIES is built using Nipype (K. Gorgolewski et al., 2011), which allows optimal resource allocation for parallelization and for a flexible workflow re-organization according to provided input parameters and dataset features (e.g. number of subjects, the availability of structural images, and the presence of multiple fMRI sessions). Log and quality control reports are automatically generated in portable, standardized formats, allowing for rapid and reliable consultation while encouraging sharing along presentations and publications.

The major pipeline steps include a registration workflow conducting essential preprocessing steps prior to group-level analysis (**figure 1A; sup. table 1**), the quality control of registration operations using a visual report (**figure 1B**), confound correction (**figure 1C**), and connectivity measurement in individual scans (**figure 1D**), altogether providing outputs prepared for subsequent statistical analyses and hypothesis testing. Outputs from each pipeline stage are generated in standardized NifTi or CSV formats, readily available for exportation and subsequent uses external to RABIES.

**Figure 1:**
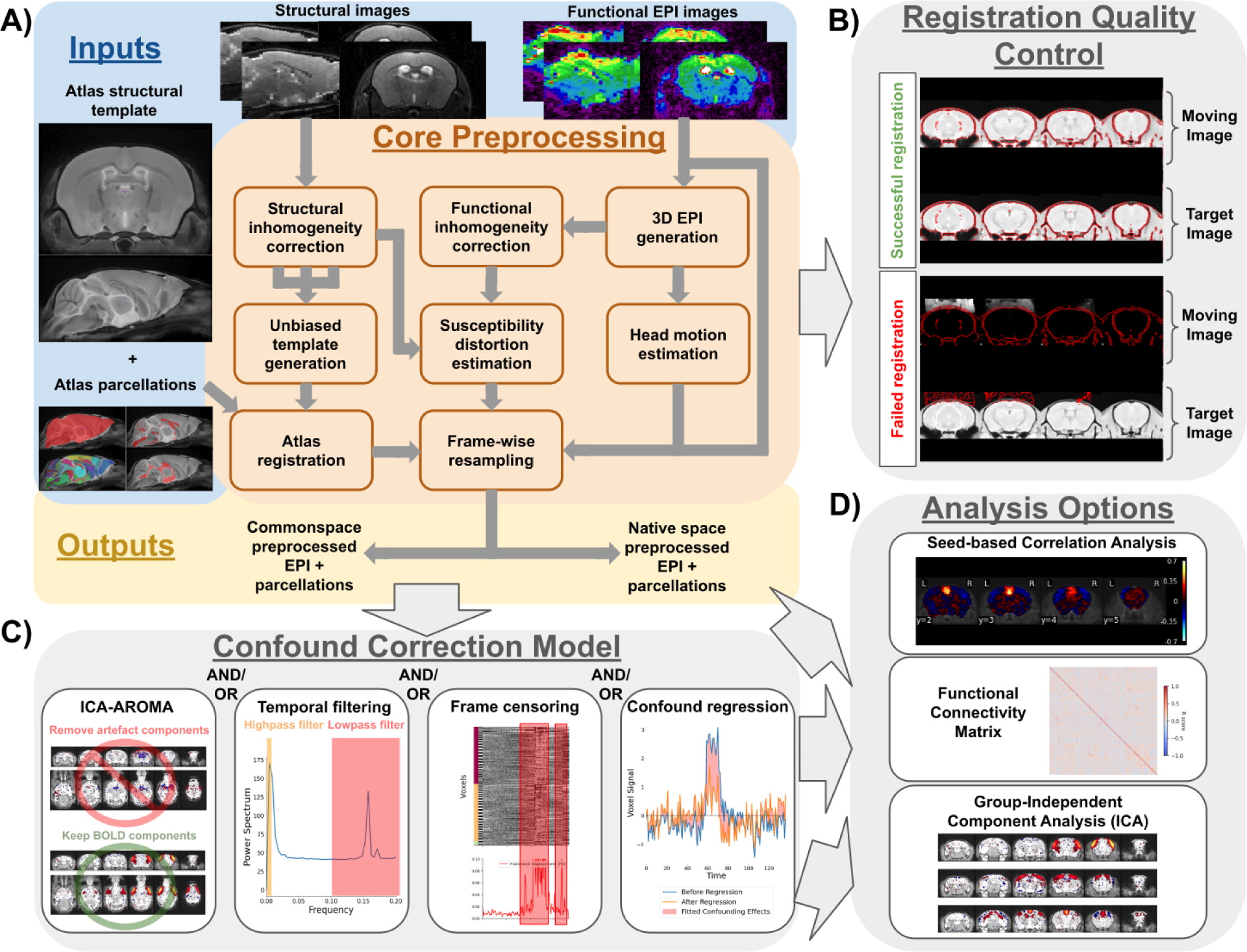
RABIES allows the preprocessing, quality control, confound correction and analysis of rodent fMRI datasets. **A)** Schematic organization of the preprocessing pipeline. RABIES takes as inputs a dataset of EPI functional scans along with their structural scan (optional) in BIDS format, together with an external reference structural atlas containing anatomical masks and labels. The core preprocessing architecture conducts commonspace alignment, head motion realignment and susceptibility distortion correction (detailed in **methods section 1; sup. table 1**). **B)** Along the execution of the preprocessing pipeline, RABIES generates automatically PNG images allowing the visual quality control of each failure-prone preprocessing step (**methods section 4**). **C)** Following preprocessing, an array of confound correction strategies are made optional and can be customized according to user needs (**methods section 5**). **D)** After applying confound correction, a final workflow is made available to conduct basic resting state analysis (**methods section 8**).

#### A generalizable registration workflow across rodent species and acquisition sites

To evaluate the robustness of our registration workflow, we used multi-site fMRI data spanning field strengths (4.7T, 7T, 9.4T, and 11.7T), coil types (room temperature and cryogenically cooled coils), anesthesia protocols (awake, isoflurane [iso], medetomidine [med], med and iso, or halothane), ventilation protocols (free breathing and ventilated), and rodent species (mice and rats) (23 publicly available datasets total, 441 mouse and 232 rat echo-planar imaging (EPI) scans, 29 EPI scans per dataset on average; **sup. table 2**). The visual quality control report (**figure 1B**) was examined across datasets to benchmark every critical registration operation throughout the pipeline, consisting of brain masking for inhomogeneity correction, alignment to commonspace and EPI distortion correction (**methods section 4; sup. figure 1**). Several innovations were required to ensure high-quality image alignment in all 23 datasets (detailed in **sup. table 3**), given the extensive variability in species, inhomogeneity artefacts, brain coverage and anatomical contrast of the EPI images (**sup. figure 2**). Despite a common workflow, mild differences in preprocessing parameters per dataset were necessary to achieve ideal performance across all datasets (listed in **sup. table 4)**.

In the improved workflow, failures were marginal (**sup. table 2; sup. figure 3**). For the 17 mouse datasets lacking structural images, we achieved a 100% (255/255) success rate for the inhomogeneity correction of EPIs, 99.6% (254/255) success during cross-subject alignment to the unbiased template. For the 3 mouse datasets including supporting structural scans, we obtained 100% success rate for both structural (110/110) and EPI (185/185) inhomogeneity correction. We achieved a success rate of 99.1% (109/110) for the alignment to the unbiased template and 100% (185/185) for the cross-modal alignment of the EPI to the structural scan. Finally, in the 3 rat datasets (1 dataset only included EPI scans), 98.9% (88/89) and 100% (230/230) of scans passed inhomogeneity correction for anatomical and EPI scans respectively, and we obtained 99.6% (230/231) success rate for cross-subject alignment and 98.9% (87/88) success rate during cross-modal alignment. Detailed reports for each dataset can be found in **sup. table 2**, and the visual quality control report from each preprocessing run is provided in the supplementary files. We consider that this high success rate can address the significant preprocessing needs within the rodent fMRI community, despite the current cross-site variability in acquisition equipment and parameters.

#### Standardized tools for confound correction and connectivity analysis

Following image preprocessing, RABIES integrates a selection of tools for conducting confound correction (**figure 1C; methods section 5)** and for deriving connectivity measures (**figure 1D; methods section 7**). The available options for confound correction (Satterthwaite et al., 2013) most standard strategies in the literature, including censoring (or scrubbing) (Power et al., 2012), confound regression (Friston et al., 1996), or frequency filtering (Satterthwaite et al., 2013). Operations are combined and orchestrated to maximize effectiveness while preventing the re-introduction of artefacts (e.g. ringing artefact when filtering prior to censoring (Carp, 2013)), as recommended in (Power et al., 2014) and (Lindquist et al., 2019). Following confound correction, connectivity can be measured in each individual scan, and results then exported for statistical testing. Analysis options include seed-based correlation (B. Biswal et al., 1995), whole-brain functional connectivity matrix generation (B. B. Biswal et al., 2010), and dual regression (Beckmann et al., 2009; Nickerson et al., 2017). Integrating standardized operations for confound correction and connectivity facilitates reproducibility of these steps as a means of improving comparisons across studies (Botvinik-Nezer et al., 2020).

### PART 2: DATA QUALITY ASSESSMENT AND GUIDELINES FOR QUALITY CONTROL OF NETWORK ANALYSIS

Defining interpretable and reliable quality control measures is challenging. Most common practices in humans include estimating confounding effects from non-neural sources (e.g.: motion or respiration) on connectivity measures (Ciric et al., 2018). Additionally in rodents, canonical network detectability varies significantly between acquisition sites both in mice (Grandjean et al., 2020) and rats (Grandjean et al., 2023) irrespective of non-neural confounds. This could be explained by the relatively lower signal to noise ratio (SNR) in small animals and/or anesthesia effects (known to decrease network activity) (Chuang & Nasrallah, 2017; Hutchison et al., 2014). Canonical networks detectability is essential for experimental reproducibility, and should be accounted for during quality control. Here, we thus define guidelines for network analysis quality control across two separate axes: spurious effects from confounds and network detectability. We identify metrics that capture key features of spurious connectivity and network detectability at the scan level (i.e. scan diagnosis), from which we derive guidelines to conduct quality control prior to group statistics. Finally, we discuss how these tools can support the improvement of data quality through optimization of confound correction, as well as the revision of analysis designs and acquisition protocols. The tools described in this section follow upon previous stages of the RABIES pipeline, leading to the generation of connectivity outputs (**figure 1**), allowing evaluation of confound correction and analysis design as well as informing subsequent statistical testing.

#### Data quality markers and pitfalls identified through scan-level diagnosis

To define an appropriate set of metrics benchmarking data quality, a large set of features were assessed from the fMRI scans across multi-site mouse datasets covering a comprehensive range of quality (N=19 datasets, 17 sites, 367/369 scans passing preprocessing quality control; see **methods section 9**). This was conducted using a scan-level diagnostic report (i.e. *spatiotemporal diagnosis*; **sup. material section 1**)) which compiles metrics featuring temporal and spatial properties of BOLD. From these observations, we determined that scan quality can be categorized into four main groups (**figure 2**): specific connectivity (i.e. the network is well detected) without confound signatures (**specific**), confounds leading to spurious connectivity (**spurious**), no network connectivity nor confound signatures (**absent**), or scans which did not neatly fit within one category, with a mixture of network and confound signatures (**mixed**). A subset of features from the spatiotemporal diagnosis are key in characterizing these quality divergences, and are detailed below:

A. **BOLD variability:** The temporal standard deviation at each voxel reflects the spatial distribution of signal variability. The resulting BOLD variability map presents an homogeneous contrast in uncorrupted scans, and can otherwise reveal the anatomical signature of confounds, thus allowing to identify the type of confound (**sup. figure 6 & 7**).
B. **Global signal covariance:** The global signal covariance map (the temporal covariance of each voxel with the global signal timecourse), is sensitive to coordinated fluctuations across voxels (i.e. does not include random noise as with the BOLD variability). As such it is sensitive to both non-neural confounds (e.g. **figure 2D**) and network signatures (e.g. **figure 2A**). The global signal covariance thus reflects whether network or confound sources dominate coordinated fluctuations, and can delineate the most likely contributors to downstream connectivity measures.
C. **Network map:** When conducting dual regression or seed-based connectivity analysis, the network connectivity map is displayed for each individual network to determine if regions known to belong to specific networks are effectively captured by the connectivity analysis (i.e. network specificity). As reported in the same multi-site datasets (Grandjean et al., 2020), in certain scans the network was absent or the network shape is distorted and shows spurious connectivity signatures.
D. **Network and confound timecourses:** Finally, the respective timecourses for networks and confounds can be compared to reveal direct relationships between network amplitude and confounds in the temporal domain. Although this metric does not describe the type of confound, it is the most direct indicator of spurious connectivity. It is an important complement to the inspection of network shape, since spurious effects may only affect amplitude with minimal impact on shape (**sup. figure 12**). To model confound timecourses, dual regression analysis is conducted with a complete set of components from Independent Component Analysis representing a mixture of networks and confounds from various origins (**sup. figure 5**), and the timecourses from confound components are compiled to summarize a broad set of potential confounds.

**Figure 2:**
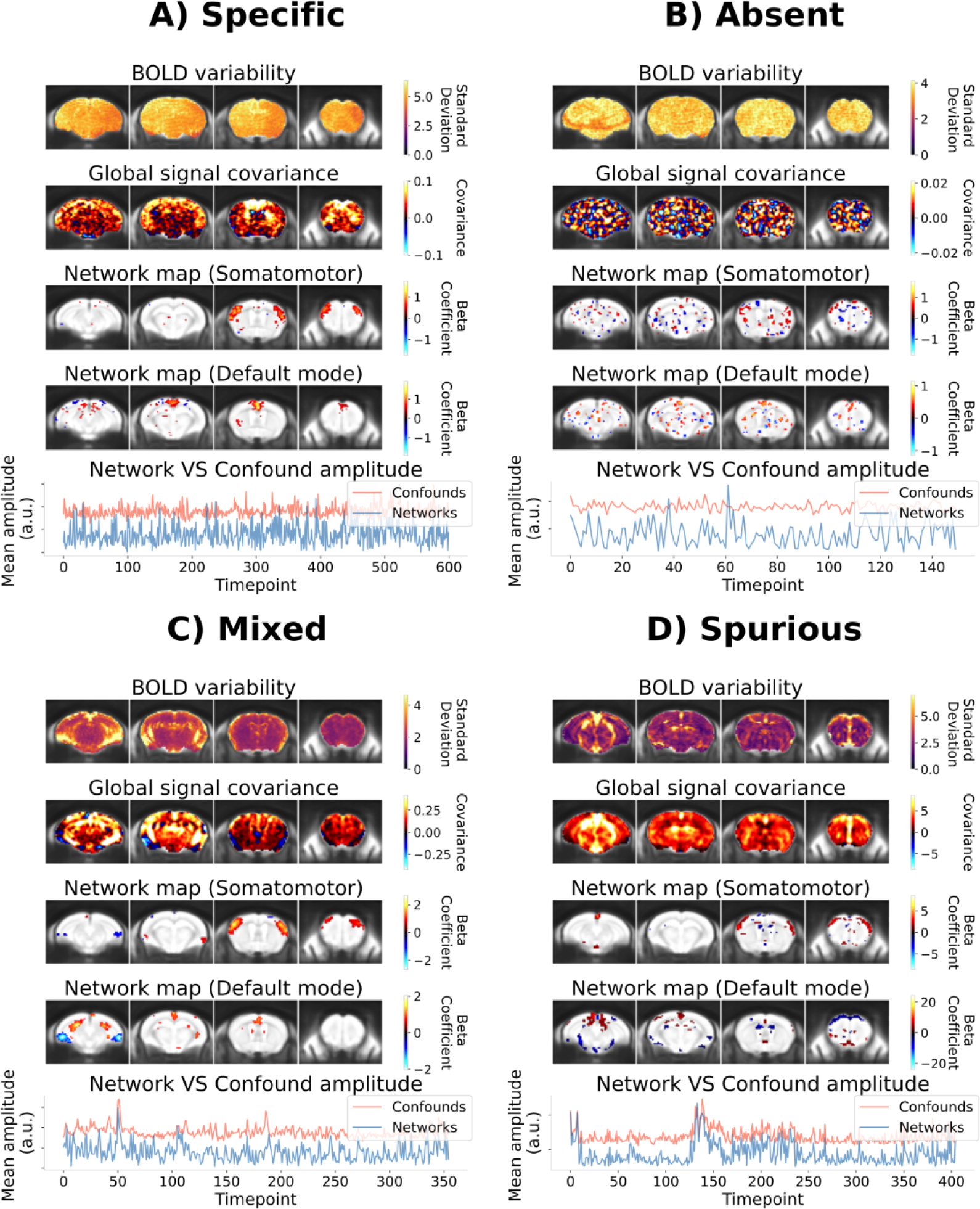
The subset of quality features delineating categories of scan quality. The BOLD variability is estimated by calculating the temporal standard deviation at each voxel, and the global signal covariance corresponds to the covariance between each voxel and the global signal (i.e. the mean signal within the brain across time). In the spatiotemporal diagnosis, a network map is generated for each network analysis conducted (in this case, the somatomotor and default mode networks were derived with dual regression). For each network analysis, a thresholded version of the network map is shown, keeping the top 4% of voxels with highest absolute value. The thresholding allows to delineate the core area of the network, and whether it reproduces the expected anatomical shape of the network. For each of these spatial features, the measures are displayed here across four coronal slices overlapped onto the commonspace reference template. Lastly, to visualize spurious relationships from the network timecourses, the amplitude of networks across time, averaged across networks (here the somatomotor and default mode networks derived from dual regression), is displayed next to the average amplitude of confound components estimated with dual regression. To derive representative averages for each category, each timecourse (from a network or confound) is initially variance normalized with an L2-norm, and an average is derived by taking the mean across the absolute-valued (i.e. positively-weighted) timecourses within a given category. For each scan category, a representative example is shown. **A)** In the specific category, the BOLD variability is mostly homogeneous, the contrast of the global signal covariance is predominantly in the gray matter without spurious features of confounds, the network maps reproduce features of the canonical network, and the networks do not display obvious correlation with confounds across time. **B)** In the absent category, the contrast from the global signal covariance and network maps is largely random. **C)** In this example scan showing mixed features, spurious signatures are observable along anatomical edges in the BOLD variability map, and overlap with features of the global signal covariance. Core network features are derived from the network maps, although there are also some spurious features. Some correlations can be noticed between the network and confound timecourses. **D)** In the spurious category, signatures of confounds predominate in all features. In this example, a confound signature is predominant in the ventricles and major brain vessels, as observed from the BOLD variability and global signal covariance. The network maps resemble spurious features, and the network timecourse is predominantly correlated with confounds.

These 4 features robustly capture the essential characteristics of network detectability and spurious connectivity at the single scan level. The remaining features from the spatiotemporal diagnosis provide additional details regarding timeseries properties, the motion parameters, or confound regression, and can further support confound characterization (**sup. table 5; sup. figure 17**). Altogether, this in-depth investigation at the scan-level allows for detailed reporting of quality features in a given dataset, and supports defining key dimensions of data quality to control for during analysis.

#### A network-level quality control framework prior to statistical analysis

While the scan-level observations described above allow for characterizing specific data quality pitfalls, it does not establish how these may impact group-level statistical inference. Statistics in network analyses are commonly controlled for using group-level quality indices relating connectivity to confound metrics. For instance, the group-level correlation between mean framewise displacement and connectivity can be estimated across subjects (Ciric et al., 2018). Here, we provide recommendations, in the form of systematic guidelines, to control for both network detectability and spurious connectivity issues prior to statistical analysis. To avoid absent and spurious connectivity (**figure 2**), scans that do not meet minimum quality thresholds are removed. Subsequently, group-level statistical reports describing the impact of data quality on inter-scan variability in connectivity are generated. To allow for the inspection of network shape (e.g. **figure 4B**) and to identify network-specific issues, these guidelines are carried on a per network basis.

Scans which fail on either standards of network detectability or spurious connectivity are removed using two scan-level measures (**figure 3**):

- **Network specificity:** The specificity of the network map relative to the canonical network is evaluated using the Dice overlap of the thresholded maps. A sufficient specificity implies both network detectability (i.e. see the absent category in **figure 2B**) and minimal spurious effects distorting network shape (see spurious scan category in **figure 2D**).
- **Confound temporal correlation:** To evaluate whether the network timecourse is dependent on known confounds (indicating spurious connectivity), the timecourse is correlated with confound sources evaluated from dual regression (after the careful identification of confound components described in **methods section 9**). This measure complements the Dice overlap by accounting for spurious effects on network amplitude, and allows to fully distinguish the spurious and absent category.

**Figure 3:**
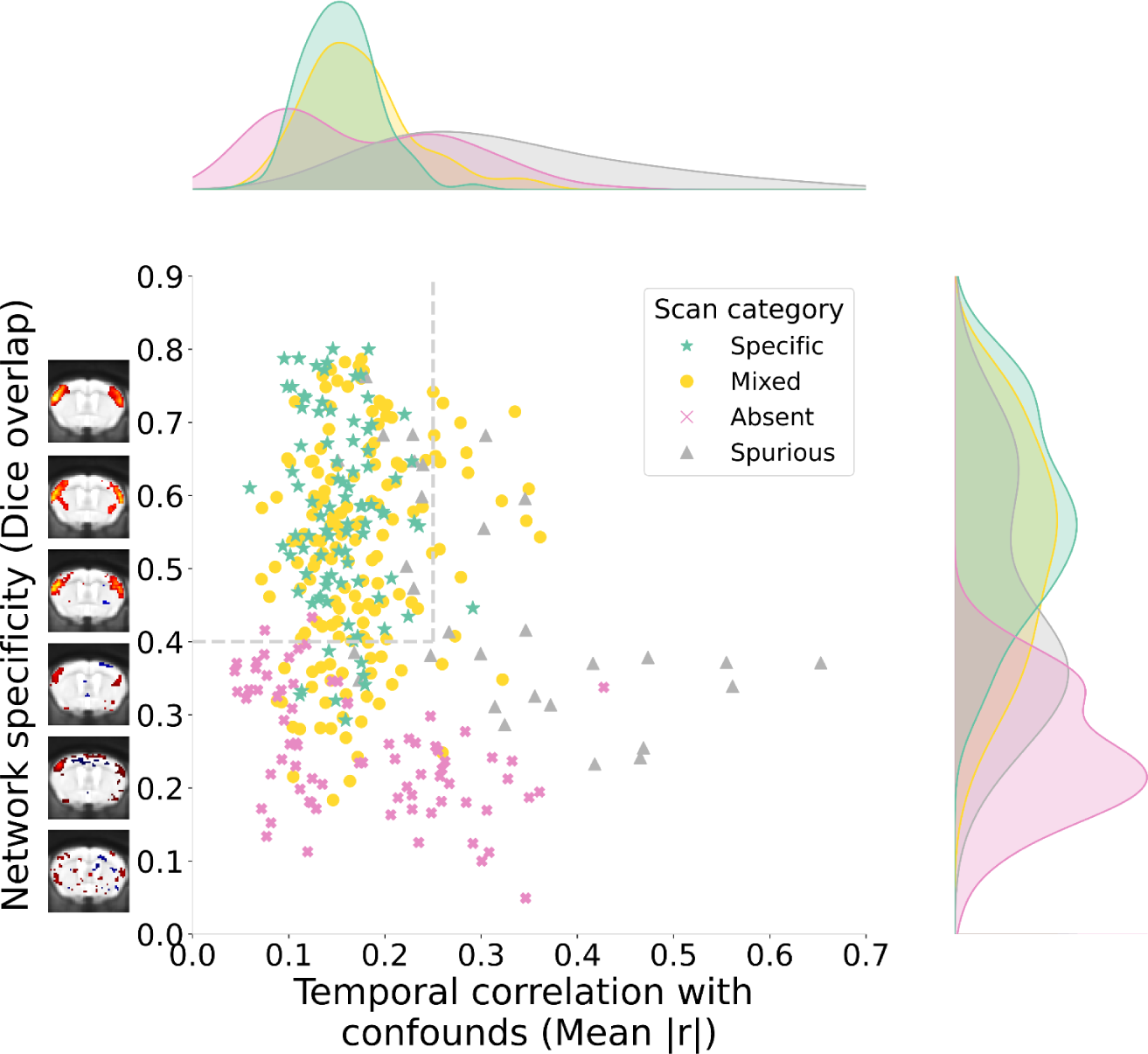
Individual scores on the measures of network specificity and confound temporal correlation across all scans from the 19 datasets investigated. The results shown correspond to the analysis of the somatomotor network with dual regression. Each dataset was assigned a predominant scan category (**sup. table 2**), and the scans are labelled accordingly. Network specificity is estimated by first thresholding the network map to identify the top 4% of voxels with highest connectivity, and then a Dice overlap is computed between the area covered by those voxels and the area covered by the corresponding canonical network map (**methods section 12**) thresholded in the same way. Both scan and canonical network maps are spatially smoothed with 0.3mm FWHM before thresholding. The temporal correlation with confounds is estimated as the mean absolute correlation between the timecourse of the network and the timecourse from each confound component obtained through dual regression. The gray dotted lines represent the selected thresholds for inclusion. The threshold for network specificity (Dice>0.4) is selected to delineate scans which reproduce network shape with little spurious features. A set of examples are shown along the axis of network specificity to represent corresponding increments in Dice overlap. For the confound temporal correlation, a threshold of 0.25 is selected to delineate scans in which clear spurious features were observed from the quality markers in figure 2.

We propose that scans which do not meet thresholds on either measure be removed from downstream statistical analysis. Network-specific thresholds should be selected on the basis of the quality markers from **figure 2** to exclude spurious and absent outcomes. Additionally, for dual regression analysis, scans which present outlier values in network amplitude are removed (**methods section 16**). This conservative approach was taken as extreme values of network amplitude may not be biologically plausible and can present additional evidence of spurious effects (Nickerson et al., 2017).

After removing scans, network detectability and spurious effects are evaluated at the group level based on inter-scan variability using the following metrics compiled in a dataset statistical report (**figure 4A**):

- **Specificity of network variability:** The inter-scan standard deviation in connectivity is computed voxelwise, creating a network variability map which is then compared to the canonical network map. In several datasets composed of scans with specific connectivity, we observed that network variability is largely specific to the anatomical extent of the network modelled. Conversely, datasets characterized by absent connectivity present random variability distributions, and heavily confounded datasets display spurious connectivity signatures (**figure 4B**). Evaluating the specificity of network variability thus allows to mitigate issues of absent or spurious connectivity which may persist at the group level despite scan-level quality control (**sup. figure 11 & 15**).
- **Confound effects:** Connectivity is correlated across scans with three complementary measures of confounds: mean framewise displacement, the variance explained from confound regression, and temporal degrees of freedom (tDOF). Mean framewise displacement is amongst the most commonly used metric (Ciric et al., 2018). However, it is limited to motion effects and does not localize confounding effects anatomically. Conversely, the variance modelled from confound regression, computed voxelwise, can reliably capture a wider variety of confound sources with anatomical specificity (**sup. figure 6 & 7**). Lastly, unequal tDOF between different scans, resulting for instance from frame censoring during confound correction, can introduce further statistical biases into the evaluation of connectivity strength. To mitigate confound effects, correlations should be minimized for all three metrics.

**Figure 4:**
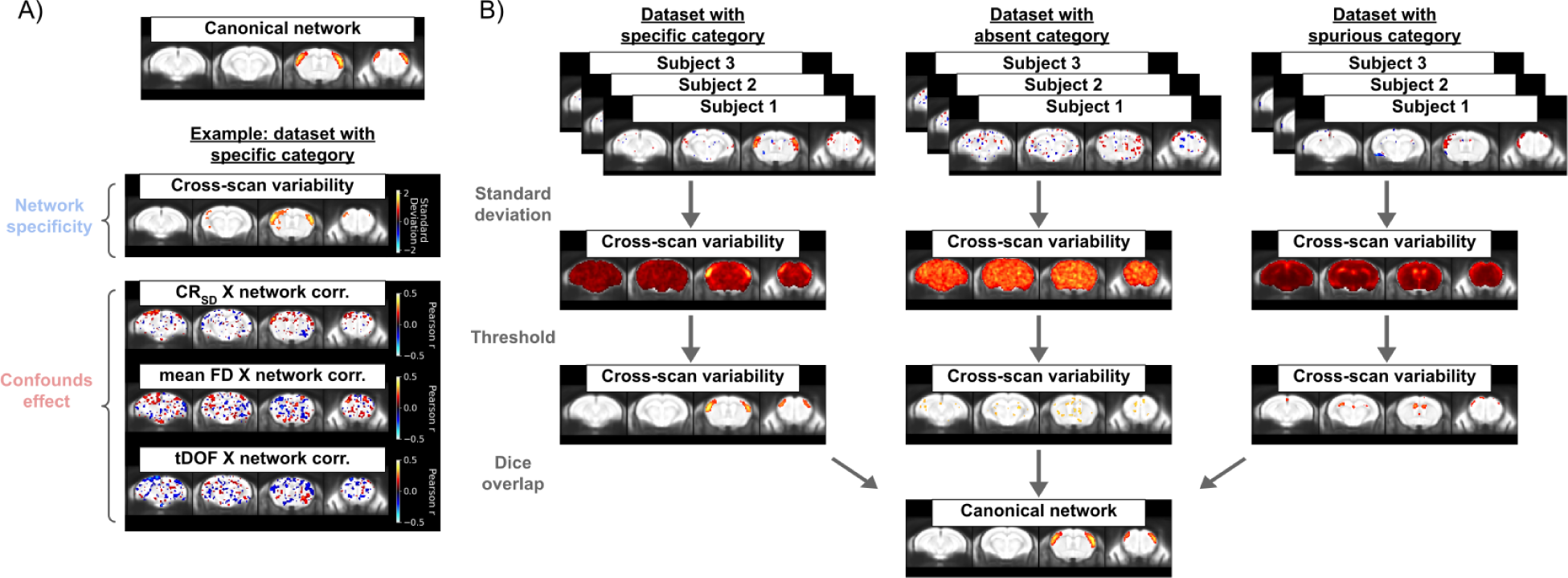
A) The group-level statistical report for quality control of network analysis. Here, an example is shown for a dataset which displayed desired outcomes for the somatomotor network using dual regression. The cross-scan variability is measured with the standard deviation at each voxel, and the resulting brain map is thresholded to include the top 4% of voxels with highest value. The predominant variability is specific to the canonical network (shown at the top, thresholded the same way). The correlation maps (thresholded at r>0.1) for each confound measures are then listed. For mean framewise displacement (mean FD) and temporal degrees of freedom (tDOF), a single scalar is derived for each scan, and correlated with connectivity across subjects for each voxel. Since the variance explained by confound regression (CR_SD_) is measured at each voxel (**sup. figure 6**), this measure is correlated voxelwise with connectivity, allowing to delineate effects which are anatomically specific (**sup. figure 15**). All brain maps in the report are spatially-smoothed with a 0.3mm FWHM kernel before display and thresholding. **B)** Schema illustrating the evaluation of the specificity of network variability and example cases for the specific, absent and spurious categories. After deriving thresholded maps of network variability and the canonical network, specificity can be summarized quantitatively by estimating the Dice overlap between the thresholded regions.

Scan-level thresholds must be applied together with the evaluation of group-level statistics as group-level metrics are inaccurate without meeting data quality assumptions at the scan-level, and group statistics inspecting inter-scan variability can reveal persisting issues pertaining to group analysis despite sensible scan-level features (**sup. table 6**). However, each quality control metric has limitations (**sup. table 6**) and certain issues are not always best captured through a simplified framework. Ideally, the interpretation of the quality control report should be supported by the inspection of the spatiotemporal diagnosis in individual scans, which provides a more comprehensive evaluation of dataset-specific issues.

#### Applications of the analysis quality control framework for improving data quality

Here we describe potential applications of these tools for improving data quality and orienting analytical design. We first explore how confound correction can be optimized according to the relevant quality issues, then discuss further applications for defining analysis design and improving acquisition protocols.

Ideally, the confound correction strategy is adapted to correct the specific issues present in a given dataset, given that excess correction can negatively impact data quality by removing network activity (Bright & Murphy, 2015). This was tested by comparing the impact of various correction methods available within RABIES on the scan-level and group-level quality. We observed important trade-offs across different aspects of data quality (e.g. reduction of confound correlation at the cost of network detectability), and important variability between datasets regarding the effectiveness of a given correction strategy (**sup. material section 3**). Thus, instead of applying the same strategy across datasets, we leveraged the spatiotemporal diagnosis and analysis quality control to define a principled approach to designing dataset-specific corrections which address relevant aspects of data quality (**methods section 17**). With this approach, correction strategies are combined incrementally, evaluating scan quality features and analysis quality control at each iteration, with the goal of maximizing network detectability and minimizing spurious effects. This process was conducted across the multi-site datasets and led to improvements in quality, both at the scan (36 more scans passing threshold for confound effects, 7 less passing for network specificity) and dataset level (3 more datasets passing threshold for network specificity and 2 more for confound effects) (**figure 5**). In particular, correlations with confounds were greatly reduced at the scan level, and network specificity was improved in datasets with spurious connectivity, suggesting correction of spurious effects on network shape. However, in datasets with low network detectability (i.e. absent connectivity), confound correction was unable to improve quality, suggesting limited network signal at data acquisition instead, which cannot be recovered by correcting confounds. Thus, this protocol offers important support for navigating the range of options for confound correction and mitigating spurious effects on connectivity, but alternatives must be considered for issues strictly targeting network detectability.

**Figure 5:**
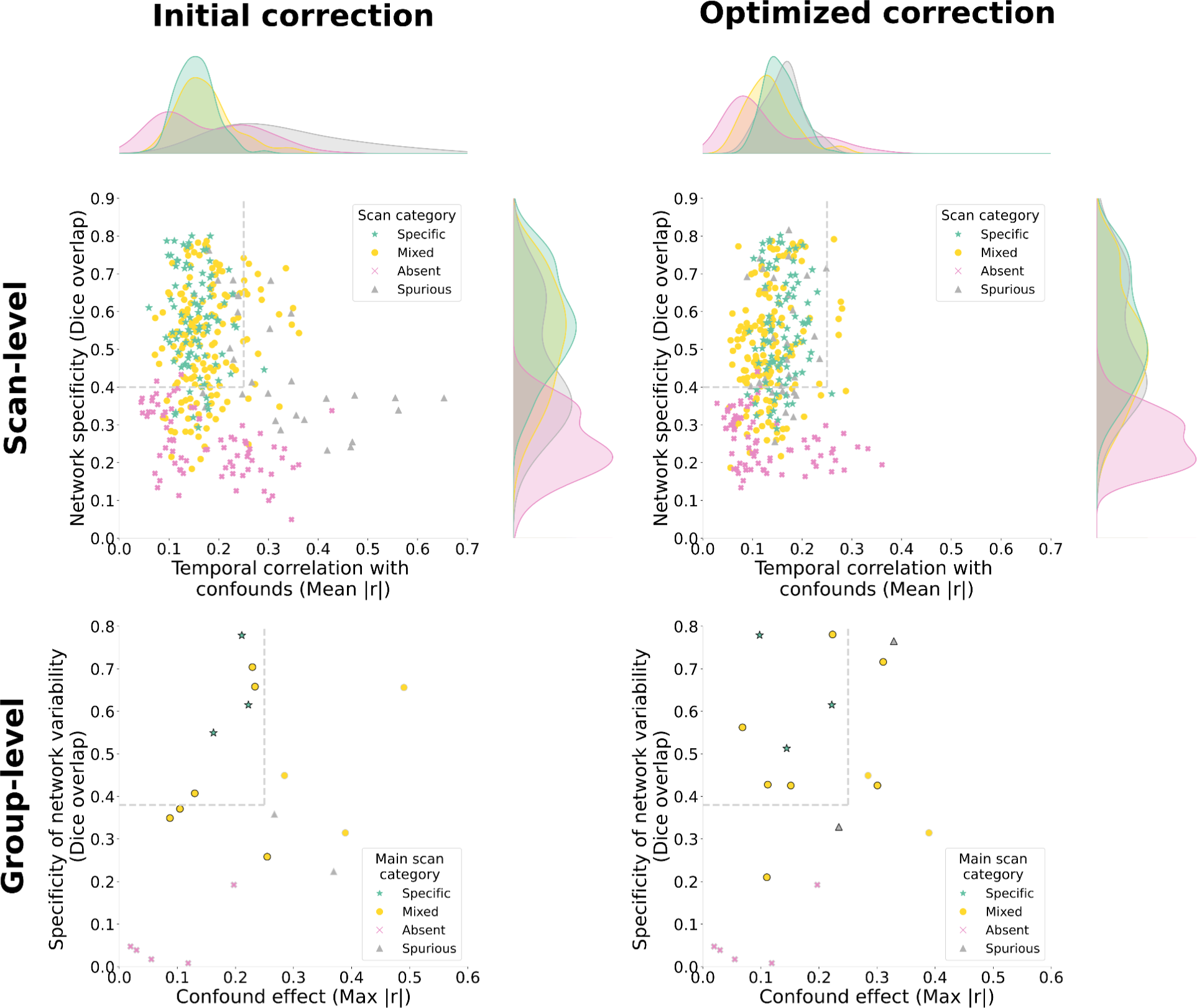
Impact of optimizing confound correction on quality control measures at the scan and group levels (results are shown for the somatomotor network measured with dual regression). At the top, scan-level measures are shown as in figure 3 before and after optimization. With optimization, the confound correlation are broadly reduced, whereas network specificity is only significantly improved in the spurious category. At the bottom, outcomes for group-level measures are summarized across all 19 datasets considered (each point is a dataset). The specificity of network variability is estimated using Dice overlap with the canonical network map as illustrated in figure 4B. For the confound effects, the mean correlation is calculated within the thresholded area of the canonical network map for each of the three confound metrics, and the maximum absolute mean correlation out of the three metrics is shown (we take the absolute value to include negative relationships). Similarly to the scan-level thresholds, the gray dotted lines represents ideal outcomes. The threshold for network variability (Dice>0.38) was determined upon visual inspection of the spatial maps to attempt distinguishing maps displaying spurious features. For the confound effects, a threshold of 0.25 was selected. Limitations to these thresholds are discussed in the supplementary discussion. Points are circled in black when at least 8 scans passed scan-level thresholds before computing group-level measures (points are circled in gray otherwise, and group-level measures are computed on all scans without applying scan-level thresholds). With the optimization of confound correction, the number of datasets passing this inclusion threshold of 8 scans increased from 9 to 12. In the absent category, we can observe that none of the 5 datasets could pass this threshold with or without optimization. The optimization increased the specificity of network variability in several datasets, where only three datasets who do not belong to the absent category do not meet the threshold. However, confound effects remain above threshold in several datasets.

Other important aspects of the experimental design which impact network quality are the analysis design and the acquisition protocol. The impact from data quality issues is influenced both by the technique selected for measuring connectivity and the specific brain network studied (**sup. figure 18 & 19**). We observed greater challenges with regards to network detectability with the default mode network than the somatomotor network, which could be explained by an increased susceptibility to physiological confounds provided its overlap with major blood vessels and ventricles (**sup. figure 10**), a narrower anatomical shape compared to the somatomotor network, an increased anesthesia susceptibility, or a reduced intrinsic activity of the network. Additionally, although seed-based connectivity does not require defining network maps a priori as with dual regression, this technique performed relatively worse on measures of network specificity. Thus, the network of interest and type of analysis may be best considered in the context of data quality, supported by network quality control guidelines. Finally, the RABIES tools can support accurately identifying underlying issues at acquisition (e.g. significant motion, physiological instability or low network detectability), and in turn, these observations may help users in designing adjustments to the acquisition protocol (e.g. improving head stability, increasing or lowering anesthesia administration) or other aspects of the experimental design. In cases where networks are detected at the scan level, but network variability is absent at the group level, more scans may need to be acquired, as this metric is dependent on sample size (**sup. figure 15B**).

Altogether, data quality assessment within RABIES supports several aspects of the analysis workflow. For best practices, these tools can be integrated throughout an experiment as it unfolds, orienting researchers in designing high quality experiments, and ensuring quality standards are met for the final analyses (**figure 6**).

**Figure 6:**
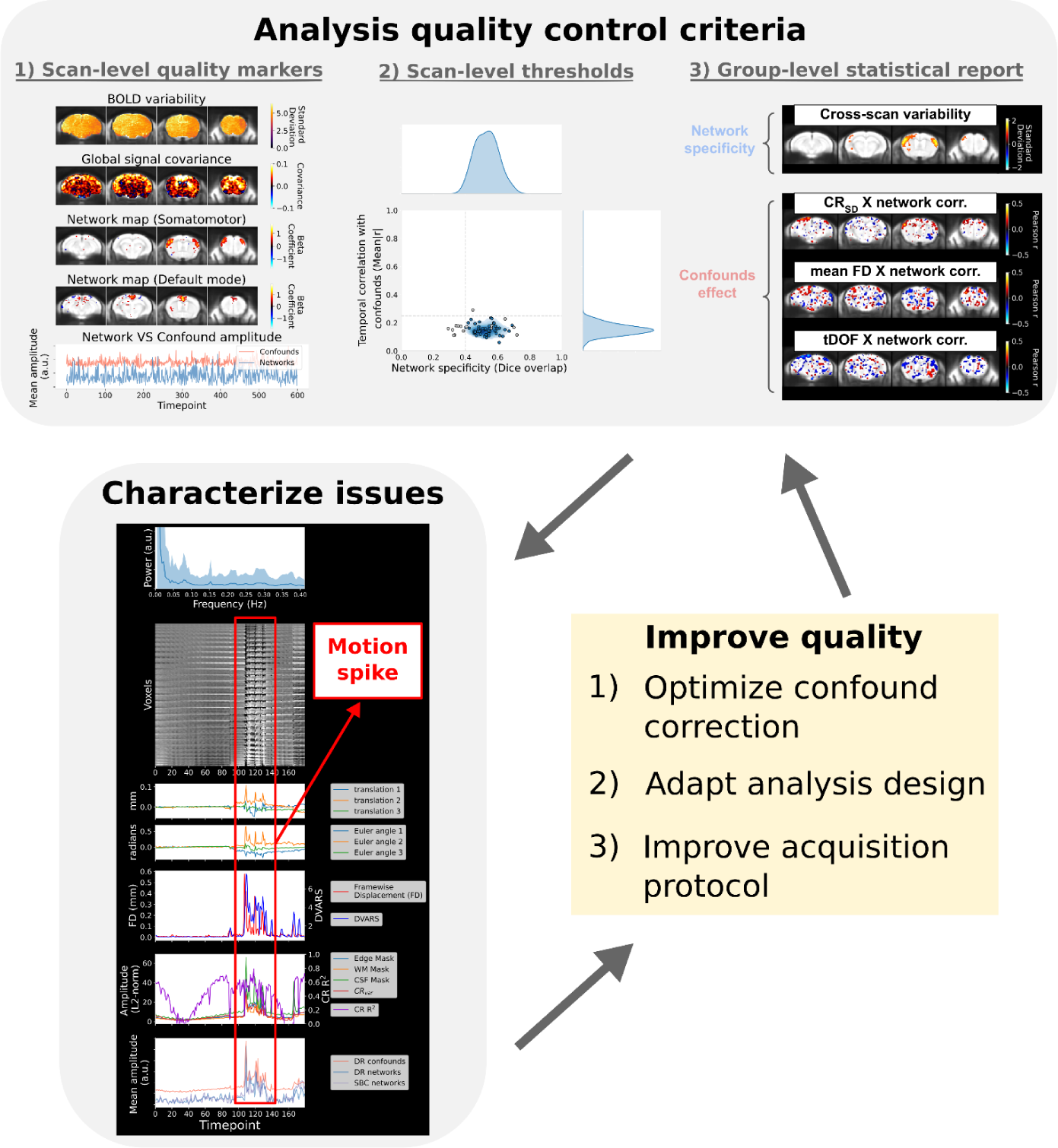
Schema summarizing the application of the RABIES data quality assessment tools. Quality control criteria are first assessed to meet the recommended standards for analysis. These consists of three complementary steps, namely the visual assessment of scan-level markers for specific connectivity, consulting the dataset distribution plot and associated files to evaluate scan-level quality thresholds (**methods section 13**), and consulting the group statistical report to evaluate expectations of network variability and group-wise confound correlation. If issues are found, these can be further characterized using the extended features from the spatiotemporal diagnosis, together with the aforementioned quality control reports, to identify plausible sources for the issue. Here, the temporal features from a scan presenting a motion spike is shown. Finally, the information provided can be leveraged to improve experimental design, either by selecting an appropriate confound correction strategy (**sup. figure 17**), by revising analysis approach, or by re-visiting the acquisition protocol. Following modifications, the quality control criteria can be re-assessed to evaluate improvements, and whether desired outcomes are achieved.

## DISCUSSION

We introduce a robust image processing and analysis pipeline adapted for specific challenges to rodent fMRI. While previous preprocessing pipelines were introduced for rodent fMRI (Celestine et al., 2020; Diao et al., 2021; Ioanas et al., 2021; Lee et al., 2021), the strengths of the proposed approach include a thorough validation across acquisition sites and species, spanning broad differences in image quality, together with the integration of state-of-the-art tools for confound correction, analysis and data quality assessment.

Motivated by the current variability in results from network analysis between mouse fMRI sites (Grandjean et al., 2020, 2023), we provide novel insights into the underlying sources of data quality divergences leading to downstream issues with network analyses. We put forward tools and guidelines for the quality control of network analysis, which support addressing potential data quality challenges and deriving informed decisions regarding experimental design. Establishing formal guidelines in those regards is challenging, and remains an ongoing topic of conversation in the human literature (Taylor et al., 2023). Hence, our approach is the first to leverage a wide array of tools for quality control (**sup. table 5**) and synthesize recommendations based on observations derived across a representative variety of rodent fMRI acquisition protocols. This is an important initial step towards providing standards for high quality rodent connectivity research, and their adoption by the wider community will improve comparability between studies, as well as foster future conversations towards addressing persisting challenges.

Despite the innovations of the RABIES software, certain limitations remain with regards to reproducibility within the current workflow. To address the substantial variability in image contrast and artefacts across rodent acquisition protocols, it remains necessary to provide flexible registration parameters which must be handled by the user. We provide a set of recommendations online to navigate the variety of registration challenges (https://rabies.readthedocs.io/en/stable/registration_troubleshoot.html). Also, the quality control of registration relies on subjective visual inspection, although this limitation is shared with human work (Craddock et al., 2013; Esteban et al., 2019). There are further concerns surrounding the design of confound correction, which could not generalize across datasets, drawing parallels with ongoing challenges discussed in the human literature regarding those decisions (Power et al., 2020; Satterthwaite et al., 2019). Here, we put forward a protocol which, informed by the reports from analysis quality control, allows identifying the most sensible strategies for correcting issues pertaining to a given dataset. Thus, instead of putting forward a single solution for confound correction, we support dataset-specific design and recommend harmonizing standards along quality control measures accounting for issues of network detectability and spurious connectivity. Similar to the quality control of preprocessing, the proposed guidelines for the quality control of network analysis rely on the visual interpretation of reports comprising several metrics. This was necessary, as generic quality control metrics have their shortcomings (**sup. table 6**), and ideally, should be complemented with the more detailed visual reports at the scan and group levels. Finally, although the quality control guidelines we present here can greatly support evaluating the main pitfalls in network analysis, these recommendations are not meant to be required irrespective of the scientific question and experimental design. The judgement of the experimenter remains paramount in determining which aspect applies to their study (for instance, network detectability may not always be expected, if studying the impact of anesthesia or inspecting a visual network in blind subjects). The guidelines are thus meant to offer a baseline which may apply to most standard resting-state network studies, as well as to instruct new researchers and RABIES users as to the main confounding factors which may impact analysis.

We provide the community with a reliable tool for conducting image processing and together with recommendations for analysis quality control, thereby allowing harmonization of computational practices across laboratories and fostering reproducible research. This advancement is timely in light of current inconsistencies in methodological and reporting practices for rodent fMRI. Previous initiatives, such as fMRIPrep, have already been transformative in those regards for human work. The RABIES analysis quality control framework will be crucial for the development of high quality data acquisition protocols (Grandjean et al., 2023). Providing these reports, automatically generated from RABIES, along with publications will improve transparency of results and prevent false positives/negatives. The adoption of RABIES for rodent fMRI will foster key practices and provide essential tools in addressing pressing issues in translational neuroscience.

## DATA AND CODE AVAILABILITY

Instructions and associated code to reproduce the results in this paper can be found in an online repository (https://github.com/Gab-D-G/RABIES_paper_repro) as well as associated supplementary files (https://doi.org/10.17605/OSF.IO/GT7EX). All MRI datasets can be found in their respective online repositories as provided in **sup. table 2**.

## ACKNOWLEDGMENTS

We thank the fMRIPrep team for their leadership in providing standards and guidelines in the development of open source neuroimaging softwares. We also thank the ICA-AROMA developper for making their code open access, which allowed us to integrate the algorithm with minor adaptations. We are grateful to all members of the mouse multicenter initiative from (Grandjean et al., 2020) as well as other members of the rodent fMRI community who publicly shared data. Without these recent initiatives, this work would not have been possible. We also thank Dr. Alessandro Gozzi and his laboratory and Mila Urosevic for providing critical feedback regarding the manuscript, Daniel Gallino for suggesting the software acronym, and the RABIES users who provided feedback to correct issues and enhance the distribution of the software. Finally, we acknowledge our funding sources. The Fonds de recherche du Québec – Nature et technologies (FRQNT) scholarship for providing salary support to G.D.-G., the Dutch Research Council grant OCENW.KLEIN.334 supports the work of JG and MMC receives salary support from the les Fonds de recherches du Québec – Santé (FRQS). Furthermore, MMCs research is supported by McGill University’s Healthy Brains for Healthy Lives (a Canada First Research Excellence Fund program), Canadian Institutes of Health Research, the National Sciences and Engineering Research Council of Canada, and a donation from the Toronto Dominion bank.

## METHODS

### 1) Core preprocessing pipeline

The preprocessing of functional magnetic resonance imaging (fMRI) scans prior to analysis consists of, at minimum, the anatomical alignment of scans to a common space, head realignment to correct for motion, and the correction of susceptibility distortions arising from the echo-planar imaging (EPI) acquisition of functional scans. The core preprocessing pipeline in Rodent Automated Bold Improvement of EPI Sequences (RABIES) carries each of these steps with state-of-the-art processing tools and techniques (detail of each step in **sup. table 1**).

To conduct common space alignment, structural images, which should be acquired along the EPI scans, are initially corrected for inhomogeneities and then registered together to allow the alignment of different MRI acquisitions. This registration is conducted by generating an unbiased data-driven template through the iterative linear and non-linear registration of each image to the dataset consensus average (using ANTs’ SyN registration algorithm (Avants et al., 2008) with Mattes Mutual Information and ANTs cross-correlation objective functions for linear and non-linear stages respectively), where the average gets updated at each iteration to provide an increasingly representative dataset template (https://github.com/CoBrALab/optimized_antsMultivariateTemplateConstruction; (Avants et al., 2011)). The finalized template after the last iteration provides a representative alignment of each MRI session to a template that shares the acquisition properties of the dataset (e.g. brain shape, field of view, anatomical contrast, …), making it a stable registration target for cross-subject alignment. This newly-generated unbiased template is then itself registered to an external reference atlas to provide both an anatomical segmentation and a common space comparable across studies defined from the provided reference atlas.

The remaining preprocessing involves the EPI images. A volumetric EPI image is first derived using a trimmed mean across the EPI frames, after an initial motion realignment step. Using this volumetric EPI as a target, the head motion parameters are estimated by realigning each EPI frame to the target using a rigid registration. To correct for EPI susceptibility distortions, the volumetric EPI is first subjected to an inhomogeneity correction step, and then registered non-linearly to the anatomical scan from the same MRI session, which allows to calculate the required geometrical transforms for recovering brain anatomy (Wang et al., 2017). Finally, after calculating the transformations required to correct for head motion and susceptibility distortions, both transforms are concatenated into a single resampling operation (avoiding multiple resampling) which is applied at each EPI frame, generating the preprocessed EPI timeseries in native space (Esteban et al., 2019). Preprocessed timeseries in common space are also generated by further concatenating the transforms allowing resampling to the reference atlas.

The preprocessing workflow of RABIES is illustrated in **figure 1A**, and each associated module is described in detail in the RABIES documentation (https://rabies.readthedocs.io/en/stable/). When structural images are not provided by the dataset, the volumetric EPI replaces the structural image during the generation of the unbiased template.

### 2) Datasets for preprocessing benchmarking

The robustness of RABIES preprocessing was evaluated by sampling a range of datasets from different sites to attempt capturing representative data samples across the rodent fMRI community. We accessed 20 open mouse fMRI datasets, 17 of which did not include structural scans, allowing for testing the workflow without structural scans. The effectiveness of the pipeline was also tested on 3 independent rat fMRI datasets. All datasets are openly accessible (**sup. table 2**), and acquisition details for each dataset can be found in their respective original publication. For mouse preprocessing using structural scans, we used the DSURQE ex-vivo MRI atlas (Dorr et al., 2008; Richards et al., 2011; Steadman et al., 2014; Ullmann et al., 2013), and the Fischer 344 in vivo atlas (Goerzen et al., 2020) was used for rat preprocessing. We generated a mouse EPI atlas for the preprocessing of mouse datasets lacking structural images (see **methods section 3** below).

### 3) Generation of a EPI atlas

We selected 5 mouse datasets (i.e. 117_Cryo_mediso_v, 7_Cryo_med_f1, 7_Cryo_med_f2, 7_Cryo_aw_f, 7_RT_med_f in **sup. table 2**) from the multi-site study by Grandjean and colleagues (Grandjean et al., 2020). We selected those datasets for their complete brain coverage, minimal susceptibility distortions and variable anatomical contrast. Using the RABIES workflow, each EPI scan was first corrected for inhomogeneity and an unbiased template was generated combining all 5 datasets. The unbiased template was then registered to the DSURQE mouse atlas using a non-linear registration (Avants et al., 2008) to propagate brain masks and anatomical labels onto the newly-generated EPI template. This EPI atlas is openly available online (https://doi.org/10.5281/zenodo.5118030), and used by default by RABIES as reference atlas when executing the EPI-only preprocessing workflow.

### 4) Preprocessing quality control

During preprocessing, a set of images are generated to conduct visual quality control of each failure-prone preprocessing step (**sup. figure 1**). To visualize the inhomogeneity correction of either the structural or the EPI image, the input MRI image is shown before and after the correction together with the overlap of the brain mask used for the final correction, thus allowing to visualize the quality of inhomogeneity correction. Failures can be noticed either if important inhomogeneities remain or if brain masking fails. Then for the QC of each registration step, the report shows the overlap between the moving image and the target image. These registration steps include alignment to the generated unbiased template, the alignment between the unbiased template and the reference common space atlas, and the registration between anatomical and functional images for susceptibility distortion correction.

The entire QC report from each dataset was visually inspected to record the number of failures at each registration step. Registration failures were considered when there was a clear misalignment of at least one brain structure. Minimal misalignments that subsisted because of EPI distortions and poor anatomical contrast were tolerated, given that very few EPIs have sufficient contrast to achieve specific anatomical alignment and complete correction for distortions. In **sup. table 2**, additional notes on preprocessing quality are provided to detail dataset-specific limitations (e.g. partial misalignment to the reference atlas in highly distorted regions). All QC reports inspected are provided along this paper in the supplementary material.

### 5) Confound correction workflow

The workflow for confound correction is structured following best practices found in human literature, largely based on recommendations from Power and colleagues (Power et al., 2014). Here, we detail the execution of the confound correction workflow, which requires specific considerations regarding the order and combination of multiple correction steps. These considerations aim to avoid inefficient corrections, or the re-introduction of confounds into the data at later steps (Lindquist et al., 2019). Several options are provided, and if selected, are applied in the following sequence:

1. **Frame censoring:** Frame censoring temporal masks are derived from framewise displacement and/or DVARS thresholds, and applied first on both BOLD timeseries before any other correction step to exclude signal spikes which may bias downstream corrections, in particular, detrending, frequency filtering and confound regression (Power et al., 2014).

a. **Censoring with framewise displacement:** Apply frame censoring based on a framewise displacement threshold. The frames that exceed the given threshold, together with 1 back and 2 forward frames will be masked out (Power et al., 2012).
b. **Censoring with DVARS:** The DVARS values are z-scored (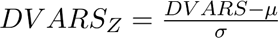, where is the mean DVARS across time, and the standard deviation), and frames with |*DVARS_Z_*| > 2.5 (i.e. above 2.5 standard deviations from the mean) are removed. z-scoring and outlier detection is repeated within the remaining frames, iteratively, until no more outliers are detected, to obtain a final set of frames post-censoring. This second censoring strategy was implemented to address residual spiking artefacts which were not well detected with a one-shot framewise displacement censoring approach (**sup. figure 8**).
2. **Detrending:** Linear (or quadratic) trends are removed from timeseries. Detrended timeseries 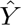 are obtained by performing ordinary-least square (OLS) linear regression,

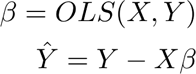

where Y is the timeseries and the predictors are *X* = [*intercept*, *time*, *time*^2^] (*time*^2^is included only if removing quadratic trends).
3. **ICA-AROMA:** Motion sources can be cleaned using a rodent-adapted ICA-AROMA classifier (Pruim et al., 2015) (**methods section 6**). ICA-AROMA is applied prior to frequency filtering to remove further effects of motion than can result in ringing after filtering (Carp, 2013; Pruim et al., 2015).
4. **Frequency filtering:** For highpass and/or lowpass filtering of timeseries, censored timepoints are first simulated while presenting the frequency profile of the timeseries to allow for the application of a butterworth filter.

a. **Simulating censored timepoints:** frequency filtering requires particular considerations when applied after frame censoring, since conventional filters cannot handle missing data (censoring results in missing timepoints). To address this issue, we implemented a method described in (Power et al., 2014) allowing the simulation of data points while preserving the frequency composition of the data. This method relies on an adaptation of the Lomb-Scargle periodogram, which allows estimating the frequency composition of the timeseries despite missing data points, and from that estimation, missing timepoints can be simulated while preserving the frequency profile (Mathias et al., 2004).
b. **Butterworth filter:** Following the simulation, frequency filtering is applied using a 3rd-order Butterworth filter, and timepoints can be removed at each edge of the timeseries to account for edge artefacts following filtering (Power et al., 2014). Edge artefacts were visualized in simulated data (**sup. figure 4**), and from those observations we conclude that removing 30 seconds at each edge and using a 3rd order filter are optimal for a highpass filter at 0.01 Hz. After frequency filtering, the temporal mask from censoring is re-applied to remove simulated timepoints.
5. **Confound regression**: For each voxel timeseries, a selected set of nuisance regressors (**sup. table 5**) are modelled using OLS linear regression and their modelled contribution to the signal is removed. Regressed timeseries 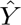 are obtained with

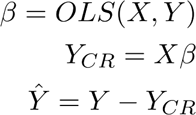

where *Y* is the timeseries, is the set of nuisance timecourses and *Y_CR_* is the confound timeseries predicted from the model at each voxel (*Y_CR_* is a time by voxel 2D matrix). Prior to the regression, the nuisance timecourses are subjected to the same frame censoring, detrending and frequency filtering which were applied to the BOLD timeseries to avoid the re-introduction of previously corrected confounds (Lindquist et al., 2019; Power et al., 2014).
6. **Intensity scaling:** Signal amplitude must be scaled to allow for comparison of network amplitude estimates between scans (Chen et al., 2017; Nickerson et al., 2017). The following options are available within RABIES:

a. **Grand mean:** Timeseries are divided by the mean intensity across the brain, and then multiplied by 100 to obtain percent BOLD deviations from the mean. The mean intensity of each voxel is derived from the linear coefficient (*β*) from the intercept computed during detrending.
b. **Voxelwise mean:** Same as grand mean, but each voxel is independently scaled by its own mean signal.
c. **Global standard deviation:** Timeseries are divided by the total standard deviation across all voxel timeseries.
d. **Voxelwise z-scoring:** Each voxel is divided by its standard deviation (i.e. z-scoring).
7. **Smoothing:** Timeseries are spatially smoothed using a Gaussian smoothing filter.

Importantly, each confound correction step (with the exception of linear detrending) is optional when using RABIES to allow for adapting correction strategy to specific dataset needs.

### 6) Rodent-adapted ICA-AROMA

The original code for the algorithm (https://github.com/maartenmennes/ICA-AROMA) was adapted to function without the hard-coded human priors for anatomical masking and parameter thresholds for component classification. Following an initial independent component analysis (ICA) decomposition of the data using FSL’s MELODIC algorithm (Beckmann & Smith, 2004), four features are extracted from each ICA component spatial map for classification: 1) the temporal correlation between the component and motion parameters, 2) the fraction of the frequency spectrum within the high frequency range, and the fraction of the component within 3) the cerebrospinal fluid (CSF) mask and 4) the brain edge mask. The component is classified as motion if the CSF or high frequency content fractions are above a given threshold, or if classified by a pre-trained linear classifier combining the brain edge fraction and motion correlation. To adapt the original algorithm with RABIES, the CSF mask is inherited from the rodent reference atlas and the edge mask is automatically generated from the brain mask, the threshold for high frequency content was increased as rodent can express higher BOLD frequencies, particularly under medetomidine (Grandjean et al., 2014), and the linear classifier was retrained. To select the new parameters, we manually classified motion and network components, derived from a set of scans from the REST-AWK group anesthetized under a medetomidine-isoflurane mixture, and selected parameters to successfully classify clear motion components while avoiding false classification of brain networks.

### 7) Standardization of variance voxelwise

Observing the distribution of signal variability across the brain (i.e. BOLD variability) allows one to identify confound signatures if present, whereas relatively uncorrupted scans present a predominantly homogeneous distribution of signal variability. To attempt mitigating the impact of confounds observed from the BOLD variability map (**figure 2**), we implemented a correction strategy to enforce an homogeneous variance distribution across voxels. To do so, timeseries are first scaled voxelwise by their standard deviation (yielding standardized variance), and then timeseries are re-scaled to preserve the original total standard deviation of the entire 4D timeseries (i.e. the global standard deviation does not change). This is to avoid changing the overall scale of image intensity, which is handled separately during confound correction (see **methods section 5**). Importantly, this is different from z-scoring, where timeseries are standardized to unit variance. Enforcing standardized variance distribution effectively downscales the contribution of voxels with higher variance, which typically corresponds to a confound signature.

### 8) Connectivity analysis within RABIES

Here we detail how each connectivity analysis available within RABIES can be conducted following the completion of the confound correction workflow.

- **Seed-based connectivity** (B. Biswal et al., 1995): An anatomical seed of interest is selected, and the mean timecourse within the seed is correlated (Pearson’s r) with each voxel timeseries to measure the brain connectivity map for that seed. In RABIES, this is repeated for each scan independently, to obtain a subject-level connectivity map that can then be used for hypothesis testing.
- **Whole-brain connectivity matrix** (B. B. Biswal et al., 2010): the whole brain parcellation during RABIES preprocessing is used to extract a mean timecourse at each parcel. Then, all parcel timecourses are cross-correlated (Pearson’s r) to obtain a whole-brain matrix describing the connectivity strength between every region pair. As with SBC, in RABIES, a connectivity matrix is computed for each scan.
- **Group independent component analysis (ICA)** (Beckmann & Smith, 2004): the timeseries from all scans are concatenated in common space, and ICA decomposition is conducted on those concatenated timeseries to identify spatial sources of covariance (components) prevalent in the entire group. Among those components, resting-state networks as well as sources of confounds can be identified (example in **sup. figure 5**). RABIES uses FSL’s MELODIC ICA algorithm (Beckmann & Smith, 2004).
- **Dual regression** (Beckmann et al., 2009; Nickerson et al., 2017): dual regression is a technique building on the group ICA algorithm which allows the modelling of previously detected group-level components back onto single scans. The dual regression algorithm consists of two consecutive linear regression steps. First, components from group ICA are regressed against an individual scan timeseries to obtain an associated timecourse for each component. Then, a second regression using these component timecourses are regressed again against the scan’s timeseries, allowing to finally derive scan-specific spatial maps for each corresponding ICA component. Those linear coefficients can allow for describing individual-specific features of resting-state networks, such as amplitude or anatomical shape. To include network amplitude, the timeseries of the first regression are variance-normalized prior to the second regression, to weight the variance explained of each component into the spatial maps (Nickerson et al., 2017). This method is applied for all dual regression analyses in this paper.

### 9) Datasets included for data quality characterization and guidelines

To explore signatures of confounds in mouse fMRI, we first relied on a dataset with a group of anesthetized mice under a medetomidine-isoflurane mixture equipped with mechanical ventilation (i.e. controlling for motion) (N=10) and a group of physically-restrained, freely-breathing, awake mice (N=9) (https://openneuro.org/datasets/ds001653/versions/1.0.2). The group of awake mice was conditioned to the restraint system for 1 week prior to scanning. fMRI (gradient echo EPI, 0.18×0.15 mm and 90×60 matrix in-plane, 28 0.4 mm slices, TR=1.2 s, 180 volumes) were acquired on a 11.7T Bruker animal scanner. For each subject, data was acquired over 2 sessions where 3 consecutive fMRI scans were acquired per session, giving a total of 60 and 53 functional scans for the anesthetized and awake group respectively (one awake scan was lost during preprocessing QC). Furthermore, to benchmark the quality control metrics and guidelines across a representative range of data quality found with mouse fMRI, we further leveraged a multicenter dataset previously harmonized by Grandjean and colleagues (17 acquisition sites, N=15 scans per site) (Grandjean et al., 2020). All datasets were preprocessed using only EPI scans, as the multicenter study did not provide structural scans.

### 10) Group-ICA priors for dual regression

The group-ICA components used as priors for dual regression (available online https://zenodo.org/record/5118030/files/melodic_IC.nii.gz) were obtained using FSL’s MELODIC algorithm (Beckmann & Smith, 2004). MELODIC was run with 30 components on the combined data from the anesthetized and awake mouse groups of the REST-AWK dataset, where confound correction consisted of highpass filtering at 0.01 Hz, FD and DVARS censoring, and confound regression was applied with the 6 rigid motion parameters together with white matter (WM)/CSF mask signals. The group-ICA maps were visually inspected, and inspired by suggestions from Zerbi and colleagues (Zerbi et al., 2015), each component was classified into a rodent brain network, confound source or other (**sup. figure 5**). More specifically, confound components display signatures of motion (i.e. edge effects), bilateral asymmetry, high loadings into WM and CSF tissues, wide-spread effects across the brain or affect only single slices. We opted for a conservative categorization of components into confounds by selecting those which primarily displayed anatomically recognizable confound features without network-like features. For network analyses in this study, we selected two components corresponding to the somatomotor and default mode networks as described in previous work (Grandjean et al., 2020; Zerbi et al., 2015).

### 11) Confound correction strategies investigated

To assess how confound correction can improve the quality of network analysis, the REST-AWK dataset and the 17 datasets from the multi-site study (Grandjean et al., 2020) were processed over a range of confound correction strategies. Datasets were initially corrected by applying a framewise displacement threshold of 0.05mm and regressing the 6 motion parameters to first establish a baseline with minimal correction.

We then evaluated the impact of the following additional corrections: voxelwise variance standardization (**methods section 7**), DVARS censoring, 24 motion parameters regression (Friston et al., 1996), WM and CSF signal regression, aCompCor regression of the first 5 principal components, global signal regression, highpass filtering and ICA-AROMA. Lowpass filtering was not considered in this study, as lowpass inflates temporal autocorrelations (Ebisuzaki, 1997) and impacts the measures of confound correlations at the scan level. Lowpass can be used in conjunction with the proposed QC framework in future studies, but would require re-evaluating an appropriate threshold separately.

In all conditions, we applied linear detrending, grand mean scaling and spatial smoothing using a 0.3 mm Full-Width at Half Maximum Gaussian filter. When highpass filtering was applied, 30 seconds were removed at each end of the timeseries to avoid edge artefacts. After censoring timepoints, scans which had less than ⅔ of the original number of frames were removed from further analysis.

### 12) Connectivity analyses investigated and definition of canonical network maps

In this study, we consider the analysis of two different mouse canonical networks, namely the somatomotor and default mode networks (Grandjean et al., 2020; Liska et al., 2015; Zerbi et al., 2015), and each network was measured with either dual regression or seed-based connectivity. For seed-based connectivity analysis, the somatomotor and default mode networks were mapped using a seed in the right primary somatosensory cortex and anterior cingulate cortex respectively. For dual regression, both networks were found among the group ICA components used as priors (**methods section 10**).

To conduct the proposed analysis quality control framework, we needed to define a reference brain map for each canonical network and each analysis technique. For dual regression, a representative map of each network was already defined from the group ICA priors. However, for seed-based connectivity analysis (**sup. figure 18 & 19**), we needed to compute a representative average brain map for each network from the set of scans across datasets. To do so, we selected a subset of the datasets with various acquisition parameters (for the somatomotor network: 7_Cryo_aw_f, 7_Cryo_mediso_v, 7_RT_halo_v, 94_RT_iso_v, REST-AWK mediso; for the default mode network: 7_Cryo_mediso_v, 7_RT_halo_v, 94_RT_iso_v, 117_Cryo_mediso_v, REST-AWK mediso) and performed well on quality metrics with dual regression analysis, when using the initial confound correction strategy (framewise displacement censoring + 6 motion parameters regression). The consensus network map was obtained by taking the median connectivity across scans. The selection of datasets was repeated independently for each network to select datasets succeeding quality control for the relevant network. The resulting consensus network maps were highly similar to the corresponding ICA components.

### 13) Dataset distribution report

To complement the group statistical report during analysis quality control (**figure 4**), as well as the selection of scan-level thresholds for network specificity and confound correlation (**figure 3**), an additional figure is generated to display the sample distribution across quality metrics (**sup. figure 9**). This report is automatically generated along the spatiotemporal diagnosis and group statistical reports and allows visualizing the relationship of both network specificity and amplitude with each measure of confound, i.e., the within scan temporal correlation with confounds, mean framewise displacement, the variance explained from confound regression and temporal degrees of freedom. For dual regression only, overall network amplitude is summarized for each scan by taking the L2-norm across the spatial map of network amplitude.

This report can be used to assess the proportion of scans which pass quality control thresholds, as well as evaluate the impact of modifying confound correction strategy on network specificity and temporal correlation with confounds. Additionally, visualizing the relationship between network amplitude and the three measures of confounds from the group statistical report allows inspecting the correlational structure between connectivity and confounds. In particular, it can be useful for detecting outliers, which are automatically identified using a threshold of 3.5 on the modified Z-score, as recommended in (Iglewicz & Hoaglin, 1993), for each measure independently (network amplitude/shape, and each confound measure). Z-scores are computed on the subset of scans which passed scan-level quality control thresholds and after removing outliers in network amplitude (**methods section 14**).

### 14) Outlier removal based on network amplitude

Scans presenting outlier values in overall network amplitude from the distribution plot were automatically removed, as this likely indicates spurious connectivity (e.g. **sup. figure 10**). After removing scans which did not pass quality control, outliers were automatically detected and removed using the modified z-score, with a threshold of 3.5 (Iglewicz & Hoaglin, 1993). If outliers were detected and removed, z-scores were re-computed on the remaining scans, and new outliers were removed. This process was repeated iteratively until no more outliers were detected to obtain a group distribution without outliers.

### 15) CR optimization protocol

We designed a stepwise protocol for optimizing confound correction on a per dataset basis, which refers to a table relating data quality features to corresponding correction strategy (**sup. figure 17A**). For all analyses, we selected the following scan-level quality control thresholds: minimum of 0.4 Dice overlap for network specificity and maximum of 0.25 correlation for confound effects (**figure 3**). The procedure applied for each dataset was as follow:

1. The dataset is initially corrected only using framewise displacement censoring and regression of 6 motion parameters, and the initial data quality reports are generated.
2. Evaluation of data quality reports:

a. Using the spatiotemporal diagnosis, the 4 main data quality markers described in **figure 2** are inspected across scans to identify potential issues.
b. The dataset distribution report is inspected to assess how many scans pass the quality control thresholds, and idenitfy outliers. If fewer than 8 scans passed, the group statistical report was not considered.
c. Inspect the group statistical report to notice absent or spurious signatures in the network variability map and evaluate the extent of confound correlation (the average correlation within the network is saved in an attached CSV file).
3. An appropriate CR strategy is selected based on the observations from the data quality reports, following the order of priority and instructions in **sup. figure 17A**.
4. One additional correction is applied at a time to evaluate the impact of individual corrections. The quality outcomes are re-evaluated as in 2), and the correction is kept only if improving quality outcomes.
5. Repeat 3)-4) until no quality issues are left, or CR options are exhausted

When evaluating the quality reports, the somatomotor - dual regression analysis was prioritized, but other analyses were also considered for improvements. The optimized set of corrections defined for each dataset is documented in **sup. figure 17B**.

## REFERENCES

Avants, B. B., Epstein, C. L., Grossman, M., & Gee, J. C. (2008). Symmetric diffeomorphic image registration with cross-correlation: evaluating automated labeling of elderly and neurodegenerative brain. Medical Image Analysis, 12(1), 26–41.

Avants, B. B., Tustison, N. J., Song, G., Cook, P. A., Klein, A., & Gee, J. C. (2011). A reproducible evaluation of ANTs similarity metric performance in brain image registration. NeuroImage, 54(3), 2033–2044.

Beckmann, C. F., Mackay, C. E., Filippini, N., & Smith, S. M. (2009). Group comparison of resting-state FMRI data using multi-subject ICA and dual regression. NeuroImage, 47(Suppl 1), S148.

Beckmann, C. F., & Smith, S. M. (2004). Probabilistic independent component analysis for functional magnetic resonance imaging. IEEE Transactions on Medical Imaging, 23(2), 137–152.

Biswal, B. B., Mennes, M., Zuo, X.-N., Gohel, S., Kelly, C., Smith, S. M., Beckmann, C. F., Adelstein, J. S., Buckner, R. L., Colcombe, S., Dogonowski, A.-M., Ernst, M., Fair, D., Hampson, M., Hoptman, M. J., Hyde, J. S., Kiviniemi, V. J., Kötter, R., Li, S.-J., … Milham, M. P. (2010). Toward discovery science of human brain function. Proceedings of the National Academy of Sciences of the United States of America, 107(10), 4734–4739.

Biswal, B., Zerrin Yetkin, F., Haughton, V. M., & Hyde, J. S. (1995). Functional connectivity in the motor cortex of resting human brain using echo-planar MRI. Magnetic Resonance in Medicine: Official Journal of the Society of Magnetic Resonance in Medicine / Society of Magnetic Resonance in Medicine, 34(4), 537–541.

Botvinik-Nezer, R., Holzmeister, F., Camerer, C. F., Dreber, A., Huber, J., Johannesson, M., Kirchler, M., Iwanir, R., Mumford, J. A., Adcock, R. A., Avesani, P., Baczkowski, B. M., Bajracharya, A., Bakst, L., Ball, S., Barilari, M., Bault, N., Beaton, D., Beitner, J., … Schonberg, T. (2020). Variability in the analysis of a single neuroimaging dataset by many teams. Nature, 582(7810), 84–88.

Bright, M. G., & Murphy, K. (2015). Is fMRI “noise” really noise? Resting state nuisance regressors remove variance with network structure. NeuroImage, 114, 158–169.

Carp, J. (2013). Optimizing the order of operations for movement scrubbing: Comment on Power et al [Review of *Optimizing the order of operations for movement scrubbing: Comment on Power et al*]. NeuroImage, 76, 436–438.

Celestine, M., Nadkarni, N. A., Garin, C. M., Bougacha, S., & Dhenain, M. (2020). Sammba-MRI: A Library for Processing SmAll-MaMmal BrAin MRI Data in Python. Frontiers in Neuroinformatics, 14, 24.

Chen, G., Taylor, P. A., & Cox, R. W. (2017). Is the statistic value all we should care about in neuroimaging? NeuroImage, 147, 952–959.

Chuang, K.-H., & Nasrallah, F. A. (2017). Functional networks and network perturbations in rodents. NeuroImage, 163, 419–436.

Ciric, R., Rosen, A. F. G., Erus, G., Cieslak, M., Adebimpe, A., Cook, P. A., Bassett, D. S., Davatzikos, C., Wolf, D. H., & Satterthwaite, T. D. (2018). Mitigating head motion artifact in functional connectivity MRI. Nature Protocols, 13(12), 2801–2826.

Craddock, C., Sikka, S., Cheung, B., Khanuja, R., Ghosh, S. S., Yan, C., Li, Q., Lurie, D., Vogelstein, J., Burns, R., & Others. (2013). Towards automated analysis of connectomes: The configurable pipeline for the analysis of connectomes (c-pac). Frontiers in Neuroinformatics, 42, 10–3389.

Diao, Y., Yin, T., Gruetter, R., & Jelescu, I. O. (2021). PIRACY: An Optimized Pipeline for Functional Connectivity Analysis in the Rat Brain. Frontiers in Neuroscience, 15, 602170.

Dorr, A. E., Lerch, J. P., Spring, S., Kabani, N., & Henkelman, R. M. (2008). High resolution three-dimensional brain atlas using an average magnetic resonance image of 40 adult C57Bl/6J mice. NeuroImage, 42(1), 60–69.

Ebisuzaki, W. (1997). A Method to Estimate the Statistical Significance of a Correlation When the Data Are Serially Correlated. Journal of Climate, 10(9), 2147–2153.

Esteban, O., Markiewicz, C. J., Blair, R. W., Moodie, C. A., Isik, A. I., Erramuzpe, A., Kent, J. D., Goncalves, M., DuPre, E., Snyder, M., Oya, H., Ghosh, S. S., Wright, J., Durnez, J., Poldrack, R. A., & Gorgolewski, K. J. (2019). fMRIPrep: a robust preprocessing pipeline for functional MRI. Nature Methods, 16(1), 111–116.

Friston, K. J., Williams, S., Howard, R., Frackowiak, R. S. J., & Turner, R. (1996). Movement-Related effects in fMRI time-series. In Magnetic Resonance in Medicine (Vol. 35, Issue 3, pp. 346–355). 10.1002/mrm.1910350312

Goerzen, D., Fowler, C., Devenyi, G. A., Germann, J., Madularu, D., Chakravarty, M. M., & Near, J. (2020). An MRI-Derived Neuroanatomical Atlas of the Fischer 344 Rat Brain. Scientific Reports, 10(1), 6952.

Gorgolewski, K., Burns, C. D., Madison, C., Clark, D., Halchenko, Y. O., Waskom, M. L., & Ghosh, S. S. (2011). Nipype: a flexible, lightweight and extensible neuroimaging data processing framework in python. Frontiers in Neuroinformatics, 5, 13.

Gorgolewski, K. J., Auer, T., Calhoun, V. D., Craddock, R. C., Das, S., Duff, E. P., Flandin, G., Ghosh, S. S., Glatard, T., Halchenko, Y. O., Handwerker, D. A., Hanke, M., Keator, D., Li, X., Michael, Z., Maumet, C., Nichols, B. N., Nichols, T. E., Pellman, J., … Poldrack, R. A. (2016). The brain imaging data structure, a format for organizing and describing outputs of neuroimaging experiments. Scientific Data, 3, 160044.

Grandjean, J., Canella, C., Anckaerts, C., Ayrancı, G., Bougacha, S., Bienert, T., Buehlmann, D., Coletta, L., Gallino, D., Gass, N., Garin, C. M., Nadkarni, N. A., Hübner, N. S., Karatas, M., Komaki, Y., Kreitz, S., Mandino, F., Mechling, A. E., Sato, C., … Gozzi, A. (2020). Common functional networks in the mouse brain revealed by multi-centre resting-state fMRI analysis. NeuroImage, 205, 116278.

Grandjean, J., Corcoba, A., Kahn, M. C., Upton, A. L., Deneris, E. S., Seifritz, E., Helmchen, F., Mann, E. O., Rudin, M., & Saab, B. J. (2019). A brain-wide functional map of the serotonergic responses to acute stress and fluoxetine. Nature Communications, 10(1), 350.

Grandjean, J., Desrosiers-Gregoire, G., Anckaerts, C., Angeles-Valdez, D., Ayad, F., Barrière, D. A., Blockx, I., Bortel, A., Broadwater, M., Cardoso, B. M., Célestine, M., Chavez-Negrete, J. E., Choi, S., Christiaen, E., Clavijo, P., Colon-Perez, L., Cramer, S., Daniele, T., Dempsey, E., … Hess, A. (2023). A consensus protocol for functional connectivity analysis in the rat brain. Nature Neuroscience, 26(4), 673–681.

Grandjean, J., Schroeter, A., Batata, I., & Rudin, M. (2014). Optimization of anesthesia protocol for resting-state fMRI in mice based on differential effects of anesthetics on functional connectivity patterns. NeuroImage, 102 Pt 2, 838–847.

Hutchison, R. M., Hutchison, M., Manning, K. Y., Menon, R. S., & Everling, S. (2014). Isoflurane induces dose-dependent alterations in the cortical connectivity profiles and dynamic properties of the brain’s functional architecture. Human Brain Mapping, 35(12), 5754–5775.

Iglewicz, B., & Hoaglin, D. (1993). Volume 16: how to detect and handle outliers, The ASQC basic references in quality control: statistical techniques, Edward F. Mykytka. Ph. D., Editor.

Ioanas, H.-I., Marks, M., Zerbi, V., Yanik, M. F., & Rudin, M. (2021). An optimized registration workflow and standard geometric space for small animal brain imaging. NeuroImage, 241, 118386.

Kurtzer, G. M., Sochat, V., & Bauer, M. W. (2017). Singularity: Scientific containers for mobility of compute. PloS One, 12(5), e0177459.

Lake, E. M. R., Ge, X., Shen, X., Herman, P., Hyder, F., Cardin, J. A., Higley, M. J., Scheinost, D., Papademetris, X., Crair, M. C., & Constable, R. T. (2020). Simultaneous cortex-wide fluorescence Ca2+ imaging and whole-brain fMRI. Nature Methods, 17(12), 1262–1271.

Lee, S.-H., Broadwater, M. A., Ban, W., Wang, T.-W. W., Kim, H.-J., Dumas, J. S., Vetreno, R. P., Herman, M. A., Morrow, A. L., Besheer, J., Kash, T. L., Boettiger, C. A., Robinson, D. L., Crews, F. T., & Shih, Y.-Y. I. (2021). An isotropic EPI database and analytical pipelines for rat brain resting-state fMRI. NeuroImage, 243, 118541.

Lindquist, M. A., Geuter, S., Wager, T. D., & Caffo, B. S. (2019). Modular preprocessing pipelines can reintroduce artifacts into fMRI data. Human Brain Mapping, 40(8), 2358–2376.

Liska, A., Galbusera, A., Schwarz, A. J., & Gozzi, A. (2015). Functional connectivity hubs of the mouse brain. NeuroImage, 115, 281–291.

Mandino, F., Cerri, D. H., Garin, C. M., Straathof, M., van Tilborg, G. A. F., Chakravarty, M. M., Dhenain, M., Dijkhuizen, R. M., Gozzi, A., Hess, A., Keilholz, S. D., Lerch, J. P., Shih, Y.-Y. I., & Grandjean, J. (2020). Animal Functional Magnetic Resonance Imaging: Trends and Path Toward Standardization. Frontiers in Neuroinformatics, 13, 78.

Mathias, A., Grond, F., Guardans, R., Seese, D., Canela, M., & Diebner, H. H. (2004). Algorithms for spectral analysis of irregularly sampled time series. Journal of Statistical Software, 11(1), 1–27.

Mills, B. D., Grayson, D. S., Shunmugavel, A., Miranda-Dominguez, O., Feczko, E., Earl, E., Neve, K. A., & Fair, D. A. (2018). Correlated Gene Expression and Anatomical Communication Support Synchronized Brain Activity in the Mouse Functional Connectome. The Journal of Neuroscience: The Official Journal of the Society for Neuroscience, 38(25), 5774–5787.

Nickerson, L. D., Smith, S. M., Öngür, D., & Beckmann, C. F. (2017). Using dual regression to investigate network shape and amplitude in functional connectivity analyses. Frontiers in Neuroscience, 11, 115.

Parkes, L., Fulcher, B., Yücel, M., & Fornito, A. (2018). An evaluation of the efficacy, reliability, and sensitivity of motion correction strategies for resting-state functional MRI. NeuroImage, 171, 415–436.

Power, J. D., Barnes, K. A., Snyder, A. Z., Schlaggar, B. L., & Petersen, S. E. (2012). Spurious but systematic correlations in functional connectivity MRI networks arise from subject motion. NeuroImage, 59(3), 2142–2154.

Power, J. D., Lynch, C. J., Adeyemo, B., & Petersen, S. E. (2020). A Critical, Event-Related Appraisal of Denoising in Resting-State fMRI Studies. Cerebral Cortex, 30(10), 5544–5559.

Power, J. D., Mitra, A., Laumann, T. O., Snyder, A. Z., Schlaggar, B. L., & Petersen, S. E. (2014). Methods to detect, characterize, and remove motion artifact in resting state fMRI. NeuroImage, 84, 320–341.

Power, J. D., Plitt, M., Laumann, T. O., & Martin, A. (2017). Sources and implications of whole-brain fMRI signals in humans. NeuroImage, 146, 609–625.

Pruim, R. H. R., Mennes, M., van Rooij, D., Llera, A., Buitelaar, J. K., & Beckmann, C. F. (2015). ICA-AROMA: A robust ICA-based strategy for removing motion artifacts from fMRI data. NeuroImage, 112, 267–277.

Richards, K., Watson, C., Buckley, R. F., Kurniawan, N. D., Yang, Z., Keller, M. D., Beare, R., Bartlett, P. F., Egan, G. F., Galloway, G. J., Paxinos, G., Petrou, S., & Reutens, D. C. (2011). Segmentation of the mouse hippocampal formation in magnetic resonance images. NeuroImage, 58(3), 732–740.

Satterthwaite, T. D., Ciric, R., Roalf, D. R., Davatzikos, C., Bassett, D. S., & Wolf, D. H. (2019). Motion artifact in studies of functional connectivity: Characteristics and mitigation strategies. Human Brain Mapping, 40(7), 2033–2051.

Satterthwaite, T. D., Elliott, M. A., Gerraty, R. T., Ruparel, K., Loughead, J., Calkins, M. E., Eickhoff, S. B., Hakonarson, H., Gur, R. C., Gur, R. E., & Wolf, D. H. (2013). An improved framework for confound regression and filtering for control of motion artifact in the preprocessing of resting-state functional connectivity data. NeuroImage, 64, 240–256.

Satterthwaite, T. D., Wolf, D. H., Loughead, J., Ruparel, K., Elliott, M. A., Hakonarson, H., Gur, R. C., & Gur, R. E. (2012). Impact of in-scanner head motion on multiple measures of functional connectivity: relevance for studies of neurodevelopment in youth. NeuroImage, 60(1), 623–632.

Schlegel, F., Sych, Y., Schroeter, A., Stobart, J., Weber, B., Helmchen, F., & Rudin, M. (2018). Fiber-optic implant for simultaneous fluorescence-based calcium recordings and BOLD fMRI in mice. Nature Protocols, 13(5), 840–855.

Steadman, P. E., Ellegood, J., Szulc, K. U., Turnbull, D. H., Joyner, A. L., Henkelman, R. M., & Lerch, J. P. (2014). Genetic effects on cerebellar structure across mouse models of autism using a magnetic resonance imaging atlas. Autism Research: Official Journal of the International Society for Autism Research, 7(1), 124–137.

Taylor, P. A., Glen, D. R., Reynolds, R. C., Basavaraj, A., Moraczewski, D., & Etzel, J. A. (2023). Editorial: Demonstrating quality control (QC) procedures in fMRI. Frontiers in Neuroscience, 17. 10.3389/fnins.2023.1205928

Ullmann, J. F. P., Watson, C., Janke, A. L., Kurniawan, N. D., & Reutens, D. C. (2013). A segmentation protocol and MRI atlas of the C57BL/6J mouse neocortex. NeuroImage, 78, 196–203.

Wang, S., Peterson, D. J., Gatenby, J. C., Li, W., Grabowski, T. J., & Madhyastha, T. M. (2017). Evaluation of Field Map and Nonlinear Registration Methods for Correction of Susceptibility Artifacts in Diffusion MRI. Frontiers in Neuroinformatics, 11, 17.

Zerbi, V., Floriou-Servou, A., Markicevic, M., Vermeiren, Y., Sturman, O., Privitera, M., von Ziegler, L., Ferrari, K. D., Weber, B., De Deyn, P. P., Wenderoth, N., & Bohacek, J. (2019). Rapid Reconfiguration of the Functional Connectome after Chemogenetic Locus Coeruleus Activation. Neuron, 103(4), 702–718.e5.

Zerbi, V., Grandjean, J., Rudin, M., & Wenderoth, N. (2015). Mapping the mouse brain with rs-fMRI: An optimized pipeline for functional network identification. NeuroImage, 123, 11–21.

