## Supplementary material for "Rodent Automated Bold Improvement of EPI Sequences (RABIES): A standardized image processing and data quality platform for rodent fMRI"

#### Section 1: A high-dimensional spatiotemporal diagnosis for data quality characterization in single scans

Leveraging the significant human literature on the impact of confounds on network analysis, we identified metrics that provide information on the relationship between the underlying BOLD signal and confounds that may bias this signal. These include temporal features and spatial features that RABIES combines into an automatically generated report for each scan derived after the confound correction step, which we term the *spatiotemporal diagnosis* (**sup. table 5; sup. figure 6**). For preliminary investigations, we selected the REST-AWK dataset that groups mouse fMRI scans with known contribution of motion artefacts (9 awake) and other scans with very minimal contributions from motion (10 anesthetized) acquired on the same 11.7T scanner. We leveraged these known differences in acquisition to support characterizing data quality features observed in the diagnosis across different scans. Below we detail the interpretation of each feature included in the diagnosis, and contrast examples of corrupted and uncorrupted scans to highlight the successful distinction between desirable BOLD sources and confounds using the spatiotemporal diagnosis, as illustrated in **sup. figure 6**:

A) **Power spectrum:** the frequency power spectrum (averaged across voxels) is displayed to assess the dominant frequency profile.
B) **Carpet plot:** the entire fMRI timeseries are displayed in a time by voxel 2D matrix. This allows to visualize global fluctuations in signal intensity, which can be a proxy for various global artefacts (Power et al., 2017).
C) **The translation and rotation head motion parameters:** those are the 6 rigid body parameters estimated during preprocessing, and allow tracking of head position across scan duration.
D) **The framewise displacement and the temporal shifts in global signal from the root-mean-square of the timeseries' temporal derivative (DVARS)** (Power et al., 2012)**:** Framewise displacement quantifies movement between consecutive frames, which reveals the timing and amplitude of spontaneous motion, whereas DVARS reveals shifts in global fMRI signal intensities. In the corrupted scan from **sup. figure 6**, we can clearly observe the important impact of motion spikes, which corrupt every temporal features including brain network timecourses modelled in **F)**. In contrast, the uncorrupted scan has a comparatively homogeneous carpet plot, framewise displacement and DVARS spikes are absent, and network timecourses are minimally aligned with motion indices.
E) **Markers of confounded fMRI signal fluctuation with anatomical masks and confound regression:** A representative timecourse is derived within a set of anatomical masks (edge, white matter and CSF masks) using the L2-norm across voxels. Each of these anatomical regions is susceptible to motion (Pruim et al., 2015), and the white

matter and CSF signal can reveal physiological confounds (Ciric et al., 2018), thus offering broader insights into potential confound sources. Furthermore, the diagnosis leverages the confound regression step, where nuisance timecourses (e.g. the 6 realignment parameters) are fitted against the fMRI signal at each voxel to obtain a modelled timecourse representing the confounded portion of the fMRI signal. To obtain an average confound timecourse across the brain, we compute the L2-norm across voxels. The proportion of variance explained by confound regression is also provided. These features allow both to visualize confound effects, and evaluate whether confound regression appropriately modelled confounds detected from other temporal features.

F) **Mean amplitude of network VS confound timecourses:** The averaged timecourse between network analyses and confound sources are compared. This is discussed in the main text (**figure 2**).

**G) Spatial distribution in signal variability ($BOLD_{SD}$):** The first spatial feature of the diagnosis is the signal variability (standard deviation) at each voxel. In **sup. figure 6**, we observe that signal variability is largely homogeneous without the major influence of confounds, but displays instead the anatomical signature of motion (i.e. anatomical edges) in the corrupted scan. This map thus offers an index of whether significant confounds are contributing to the signal.

**H) Confound regression variance explained ($CR_{SD}$):** Here, the variance explained from confound regression is quantified at each voxel by taking the standard deviation from the modelled confound timecourse (see **E)**). This allows to contrast spatially the amplitude of confound effects. $CR_{SD}$ is the only spatial feature revealing the expected signature of motion along anatomical edges for both uncorrupted and corrupted examples (although more prominent in the corrupted scan). This suggests that this feature can both reliably and specifically reflect the presence of confounds, better so than alternative features.

**I) Confound regression variance explained proportion:** Similar to $CR_{SD}$, but showing instead the proportion of variance explained ($R^2$).

**J) Global signal covariance:** The correlation with the global signal was used previously to identify globals sources of confounds (Power et al., 2017). Here we select instead the covariance to preserve the inhomogeneous distribution of voxel variance, which is relevant for the detection of confounds in **G)**. By visualizing the spatial contrast in the covariance of each voxel with the global signal timecourse, we can clearly observe the mixed nature of the global signal, which is known to capture the effect from both confounds (Power et al., 2017) and neural activity (Schölvinck et al., 2010). Indeed, in the uncorrupted scan, the global signal has a predominant contrast along the cortical gray matter, with anatomical features reminiscent of brain networks. On the other hand, in the corrupted scan, the global signal clearly reflects the spatial contrast of the motion confound, whereas applying confound correction effectively recovers a predominant cortical contrast. This feature can thus offer an index of whether confounding sources or network activity are dominant in the data.

**K) Network spatial maps:** Finally, the diagnosis shows the spatial network maps fitted using dual regression (or seed-based analysis) from the selected set of brain networks of interest (in this case the somatomotor and default mode networks). These fits provide insights into the quality of network analysis, and how they may affect downstream statistical analyses (see main text).

Outside of motion spikes, we could derive similar conclusions for two alternative sources of confound related to slow global fluctuations (**sup. figure 7**). By relying on the two spatial features of signal variability, i.e. $BOLD_{SD}$ and $CR_{SD}$, we find evidence that one of the confounds is associated to a ventricular origin and the other to a vascular origin, but the temporal features reveal that only the latter significantly influences network estimates from dual regression. In all of these confound cases, the temporal features reveal key mechanistic interactions between the presence of each confound and other aspects of the data, including network analysis. Subsequently, complementing the temporal information, $CR_{SD}$ can identify the specific source of confounds by inspecting its anatomical features within the brain. Finally, the $BOLD_{SD}$ and the global signal covariance can reveal whether signal variability is dominated by confound sources or network activity, and whether confounds persist after applying a correction. These observations demonstrate the complementarity of temporal and spatial features in the interpretation of signal sources, and together, these features allow the diagnosis of most major sources of confounds and can support their correction in rodent fMRI data.

#### Section 2: Discussion on the expectation of network variability specificity

As part of the quality control of network analysis, we put forward the recommendation that inter-subject variability in network connectivity should be predominant within the core anatomical delineation of the canonical network. This recommendation is grounded in observations presented in this manuscript, where datasets with relatively low evidence of confounds and clear network detectability tend to express those features of network variability at the group level (both for dual regression and seed-based connectivity analyses, and for the somatomotor and default mode networks). We argue that benchmarking network variability is important for accurately addressing certain data quality shortcoming that would be missed otherwise. We recognize, however, that the nature of inter-individual variability in network connectivity is not fully understood. In particular, it is not clear how to interpret cases where network variability is absent despite sufficient network detectability at the scan level and no evidence of spurious effects (**sup. figure 15A**). Such cases raise the question of whether the specific network variability should be necessarily expected.

Providing conclusive evidence on this issue would require a comprehensive assessment of the relationship between network variability as observed through BOLD fMRI and underlying neural metabolism. While we do not aim to reach such conclusions in this study, we emphasize several considerations. First, the specificity of network variability relates strongly to sample size (**sup. figure 15B**), and thus, absent variability could be indicative of shortcomings in statistical power. Sample sizes were relatively low for most datasets, which may explain the failure to detect such variability for a subset of the datasets. Second, cross-scan variability can transition from absent to specific following the optimization of confound correction (**sup. figure 15C**). We would thus recommend attempting improving confound correction before concluding that variability cannot be detected. Finally, we are providing here a generic measure of variability (standard deviation across scans) which can be applied on any set of scans, but alternatives could be considered in function of the experimental design. For instance, in the context of studying group differences, inspecting the spatial contrast of group differences (i.e. the group difference reproduces network features) can be a better indicator that the statistical analyses relate to the network of interest. Thus, we do not suggest that the built-in approach within RABIES is the only valid approach for assessing that statistical analyses relate to the network of interest. We do argue however that, provided the susceptibility of fMRI connectivity analyses to false findings, it is preferable to provide evidence that connectivity results relate to expected anatomical features from the network of interest.

#### Section 3: The impact of each confound correction strategies across datasets

To evaluate the ability of confound correction to improve analysis quality outcomes and investigate whether specific strategies can provide generalizable improvements across datasets, the main strategies available within RABIES were all tested systematically across the 19 datasets considered in part 2 of the manuscript (**sup. figure 16**). More specifically, datasets were initially processed with the regression of 6 motion parameters and censoring with framewise displacement, then one additional correction was applied. The impact of the correction was evaluated across quality metrics part of the quality control guidelines (**figure 3 & 4**). Outcomes were also evaluated when removing initial corrections (using only censoring, or no correction applied) to inspect potential impacts of the baseline correction and benchmark the ability of quality control metrics in detecting problematic quality outcomes.

Overall, there was important variability in the impact of additional corrections across datasets. Although certain corrections predominantly yielded improvements in quality metrics (i.e. standard, WM/CSF and aCompCor), no single correction led to uniform increase in the number of scans passing network specificity and confound correlation thresholds across all datasets. The application of no correction (raw) or only framewise displacement censoring led to an almost uniform decrease in quality, thus justifying the recommendation of regressing 6 motion parameters and applying censoring as a baseline. The impact of each correction, as reported in **sup. figure 16** (together with similar reports for other connectivity analyses), supported defining recommendations for optimizing confound correction in **sup. figure 17** according to the quality metrics of network specificity and confound correlation.

### SUPPLEMENTARY TABLES

| Processing module | Description |
|---|---|
| Structural inhomogeneity correction | The image is denoised with non-local mean denoising (Manjón et al., 2010) (for EPIs, denoising was carried beforehand during 3D EPI generation) followed by iterative correction for intensity inhomogeneities (Sled et al., 1998). Initial masking is achieved via intensity thresholding, giving an initial correction of the image, and a registration is then conducted to register a brain mask for a final round of correction. |
| Unbiased template generation | Generates a dataset-specific unbiased template from the input structural images. Through a set of iterations, images are registered to a consensus average generated from the overlap of all scans at the previous iteration. Registrations are progressively more flexible, executing 2 iterations each for a rigid, similarity, affine and finally non-linear template generation. The final iteration provides the individual transforms to align each scan to the unbiased template. This template generation process creates a robust target for the alignment of MRI sessions sharing the same acquisition properties, which will minimize registration inconsistencies between sessions, as opposed to the direct registration to an external template. The algorithm is implemented in https://github.com/CoBrALab/optimized_antsMultivariateTemplateConstruction. |
| Atlas registration | A affine and non-linear registration is conducted to align the unbiased template with the structural atlas reference, hence providing the transforms to propagate associated anatomical masks and parcellations. |
| Generate 3D EPI | The 4D raw EPI file is used to generate a representative volumetric 3D EPI. This volume later becomes the target for motion realignment and the estimation of susceptibility distortions through registration to the structural image. Two iterations of motion realignment to an initial median of the volumes are conducted, then a trimmed mean is computed on the realignment volumes, ignoring 5% extreme, and this average becomes the reference image. The final image is then corrected using non-local means denoising (Manjón et al., 2010). |
| Functional inhomogeneity correction | Same as structural inhomogeneity correction, but conducted on the 3D EPI reference. |
| Susceptibility distortion correction | The input volumetric EPI image is registered non-linearly to an associated structural MRI image. The non-linear transform estimates the correction for EPI susceptibility distortions (Wang et al., 2017). |
| Head motion estimation | This workflow estimates motion during fMRI acquisition. To do so, each EPI frame is registered to a volumetric target reference image with a rigid registration using ANTs' antsMotionCorr algorithm (Avants et al., 2009). This results in the measurement of 3 Euler angles in radians and 3 translations in mm (from ITK's Euler3DTransform https://itk.org/Doxygen/html/classitk_1_1Euler3DTransform.html) at each time frame, which are then stored into an output CSV file. |
| Frame-wise resampling | This module carries out the resampling of the original EPI timeseries into preprocessed timeseries. This is accomplished by applying at each frame a combined transform which accounts for previously estimated motion correction and susceptibility distortion correction, together with the alignment to common space if the outputs are desired in common space. All transforms are concatenated into a single resampling operation to mitigate interpolation effects from repeated resampling. This workflow also carries the resampling of brain masks and labels from the reference atlas onto the preprocessed EPI timeseries. |

**Supplementary table 1:** Core preprocessing modules within RABIES.

| Online link | Dataset name | Species | Structural - Inhomogeneity correction | Functional - Inhomogeneity correction | Unbiased template generation | Atlas registration | Susceptibility distortion correction | Additional notes on registration quality control | Predominant scan quality category (figure 2) |
|---|---|---|---|---|---|---|---|---|---|
| https://openneuro.org/datasets/ds001720/versions/1.0.2 | Multisite - 117_Cryo_iso_f | Mouse | - | 15/15 | 15/15 | Succesful | - | | Spurious |
| https://openneuro.org/datasets/ds001720/versions/1.0.2 | Multisite - 117_Cryo_mediso_v | Mouse | - | 15/15 | 15/15 | Succesful | - | | Specific |
| https://openneuro.org/datasets/ds001720/versions/1.0.2 | Multisite - 47_RT_iso_f | Mouse | - | 15/15 | 15/15 | Succesful | - | | Mixed |
| https://openneuro.org/datasets/ds001720/versions/1.0.2 | Multisite - 7_Cryo_aw_f | Mouse | - | 15/15 | 15/15 | Succesful | - | | Mixed |
| https://openneuro.org/datasets/ds001720/versions/1.0.2 | Multisite - 7_Cryo_med_f1 | Mouse | - | 15/15 | 15/15 | Succesful | - | | Absent |
| https://openneuro.org/datasets/ds001720/versions/1.0.2 | Multisite - 7_Cryo_med_f2 | Mouse | - | 15/15 | 15/15 | Succesful | - | Some heavy distortions near the ear canals couldn't be completely accounted for by nonlinear registration. | Absent |
| https://openneuro.org/datasets/ds001720/versions/1.0.2 | Multisite - 7_Cryo_med_f3 | Mouse | - | 15/15 | 14/15 | Succesful | - | The posterior of the brain wasn't well aligned for one subject because of the field of view. | Mixed |
| https://openneuro.org/datasets/ds001720/versions/1.0.2 | Multisite - 7_Cryo_mediso_v | Mouse | - | 15/15 | 15/15 | Succesful | - | | Specific |
| https://openneuro.org/datasets/ds001720/versions/1.0.2 | Multisite - 7_RT_halo_v | Mouse | - | 15/15 | 15/15 | Succesful | - | | Mixed |
| https://openneuro.org/datasets/ds001720/versions/1.0.2 | Multisite - 7_RT_iso_f | Mouse | - | 15/15 | 15/15 | Succesful | - | | Absent |
| https://openneuro.org/datasets/ds001720/versions/1.0.2 | Multisite - 7_RT_med_f | Mouse | - | 15/15 | 15/15 | Succesful | - | | Absent |
| https://openneuro.org/datasets/ds001720/versions/1.0.2 | Multisite - 94_Cryo_iso_f | Mouse | - | 15/15 | 15/15 | Succesful | - | | Spurious |
| https://openneuro.org/datasets/ds001720/versions/1.0.2 | Multisite - 94_Cryo_med_f | Mouse | - | 15/15 | 15/15 | Succesful | - | EPI distortions on the top of the brain couldn't be aligned with the target atlas. | Mixed |
| https://openneuro.org/datasets/ds001720/versions/1.0.2 | Multisite - 94_Cryo_mediso_v | Mouse | - | 15/15 | 15/15 | Succesful | - | | Mixed |
| https://openneuro.org/datasets/ds001720/versions/1.0.2 | Multisite - 94_RT_iso_v | Mouse | - | 15/15 | 15/15 | Succesful | - | | Mixed |
| https://openneuro.org/datasets/ds001720/versions/1.0.2 | Multisite - 94_RT_mediso_f1 | Mouse | - | 15/15 | 15/15 | Succesful | - | EPI distortions near the ear canals couldn't be completely accounted for by nonlinear registration. | Absent |
| https://openne | Multisite - | Mouse | - | 15/15 | 15/15 | Succesful | - | | Mixed |

| | | | | | | | | | |
|---|---|---|---|---|---|---|---|---|---|
| uro.org/datasets/ds001720/versions/1.0.2 | 94_RT_mediso_f2 | | | | | | | | |
| | **Total mice (Multisite dataset):** | Mouse | - | 255/255 | 254/255 | 17/17 | - | | - |
| https://openneuro.org/datasets/ds001653/versions/1.0.2 | REST-AWK | Mouse | 38/38 | 114/114 | 38/38 | Succesful | 114/114 | Inconsistent alignment in ventral areas below hippocampi. | - |
| https://openneuro.org/datasets/ds001653/versions/1.0.2 | REST-AWK - bold-only | Mouse | - | 113/114 | 113/113 | Succesful | - | | mediso: specific awake: Mixed |
| https://openneuro.org/datasets/ds004125/versions/1.0.0 | Montreal, Douglas CIC | Mouse | 20/20 | 19/19 | 19/20 | Succesful | 19/19 | | - |
| https://openneuro.org/datasets/ds001890/versions/1.0.1 | Singapore, 3xTGAD | Mouse | 52/52 | 52/52 | 52/52 | Succesful | 52/52 | An hydrocephalic mouse was excluded. | - |
| | **Total mice (with anat):** | Mouse | 110/110 | 185/185 | 109/110 | 3/3 | 185/185 | | - |
| https://openneuro.org/datasets/ds001981/versions/1.0.3 | Rat Guttingen | Rat | 23/24 | 23/23 | 23/23 | Succesful | 22/23 | EPI distortions near the ear canals couldn't be completely accounted for by nonlinear registration. | - |
| https://www.nitrc.org/projects/rat_rsfmri | Rat NITRC | Rat | 65/65 | 64/64 | 64/65 | Succesful | 64/64 | A ghosting artefacts caused one failure in the anatomical images. | - |
| https://openneuro.org/datasets/ds003646/versions/1.0.0 | Rat CAMRI | Rat | - | 143/143 | 143/143 | Succesful | - | | - |
| | **Total rats:** | Rat | 88/89 | 230/230 | 230/231 | 3/3 | 87/88 | | - |

**Supplementary table 2:** 23 rodent fMRI datasets and results from the quality control of preprocessing. Also, the predominant category for scan data quality markers, as described in **figure 2**, is listed for each dataset included for part 2 of the results. This was determined through visual inspection of the data quality markers in individual scans for each dataset.

| Registration challenge | Solution within RABIES |
|---|---|
| Parameter selection for the inhomogeneity correction and registration algorithms are dependent on brain size, and are thus inconsistent across rodent species. | We developed a scale-independent registration algorithm, wrapped around antsRegistration (Avants et al., 2008), which automatically adapts parameters to species-specific physical dimensions. The provided commonspace target is used to compute brain dimensions, and automatically define the parameters for the inhomogeneity correction and registration operations (https://github.com/CoBrALab/minc-toolkit-extras/blob/master/ants_generate_iterations.py) |
| Poor anatomical contrast led to several EPI masking failures during inhomogeneity correction. | 1. We created an EPI reference atlas using mice from different sites with whole-brain coverage and varying anatomical contrast (**methods section 3**). Using this atlas as reference for workflows lacking structural images improved substantially the quality of masking during EPI inhomogeneity correction and of alignment from EPI-generated unbiased templates to the reference atlas.<br>2. We created an option to skip registration during inhomogeneity correction, and instead rely only on a Otsu thresholding (Otsu, 1979) strategy for masking. The number of thresholds derived from the Otsu method can be manually selected to fit the intensity distribution of the brain for a given dataset. |
| Incomplete brain coverage or the blurring of brain boundaries results in misalignment during template generation and atlas registration. | Three innovations were required:<br>1. Using a staged template construction with increasing orders of registration (rigid, similariy, affine and finally non-linear registration) (https://github.com/CoBrALab/optimized_antsMultivariateTemplateConstruction) helped ensure robust initial alignment of images before computing downstream nonlinear template. This was most crucial for datasets with incomplete brain coverage.<br>2.<br>3. During inhomogeneity correction, a preliminary brain mask is computed (see **sup. table 1**). Following preliminary alignment for the unbiased template generation, these brain masks can be combined into a consensus brain mask (through majority vote in commonspace). This consensus mask can then support registration by computing the registration similarity metric only within the area defined by the mask, and ignoring other areas. This masking strategy allowed improving alignment during unbiased template generation and subsequent atlas registration.<br>4. Finally, a brain extraction option for registration was added, where at each registration step, brain masks are used to remove outside tissues and enhance brain edge contrast. As in 2. above, the preliminary brain mask from inhomogeneity correction was provided for EPI brain extraction. |
| For datasets lacking anatomical scans, the EPI-generated unbiased template frequently failed alignment with the DSURQE structural atlas. | In mice, when conducting preprocessing without anatomical scans, the EPI-generated unbiased template is registered to the EPI reference atlas mentioned above instead. |

**Supplementary table 3:** Innovations required to offer generalizable registration for rodent fMRI.

| Dataset name | EPI-only preprocessing | Structural - Inhomogeneity correction | Functional - Inhomogeneity correction | EPI - Otsu threshold | Atlas registration | Susceptibility distortion registration | Use EPI masking for unbiased template generation | Apply brain extraction | Minimum number of timepoints | Removing first 5 timepoints |
|---|---|---|---|---|---|---|---|---|---|---|
| Multisite - 117_Cryo_iso_f | TRUE | - | Rigid registration | 2 | Non-linear registration | - | TRUE | FALSE | 300/450 | TRUE |
| Multisite - 117_Cryo_mediso_v | TRUE | - | Rigid registration | 2 | Non-linear registration | - | TRUE | FALSE | 400/600 | FALSE |
| Multisite - 47_RT_iso_f | TRUE | - | Rigid registration | 2 | Non-linear registration | - | TRUE | TRUE | 196/295 | FALSE |
| Multisite - 7_Cryo_aw_f | TRUE | - | Rigid registration | 2 | Non-linear registration | - | TRUE | FALSE | 260/390 | FALSE |
| Multisite - 7_Cryo_med_f1 | TRUE | - | Rigid registration | 2 | Non-linear registration | - | TRUE | FALSE | 266/400 | TRUE |
| Multisite - 7_Cryo_med_f2 | TRUE | - | Rigid registration | 2 | Non-linear registration | - | TRUE | FALSE | 200/300 | TRUE |
| Multisite - 7_Cryo_med_f3 | TRUE | - | Rigid registration | 2 | Non-linear registration | - | TRUE | FALSE | 266/400 | FALSE |
| Multisite - 7_Cryo_mediso_v | TRUE | - | Rigid registration | 2 | Non-linear registration | - | TRUE | FALSE | 666/1000 | FALSE |
| Multisite - 7_RT_halo_v | TRUE | - | Rigid registration | 2 | Non-linear registration | - | TRUE | FALSE | 266/400 | FALSE |
| Multisite - 7_RT_iso_f | TRUE | - | No reg | 4 | Non-linear registration | - | TRUE | TRUE | 100/150 | FALSE |
| Multisite - 7_RT_med_f | TRUE | - | Rigid registration | 2 | Non-linear registration | - | TRUE | FALSE | 333/500 | TRUE |
| Multisite - 94_Cryo_iso_f | TRUE | - | Rigid registration | 2 | Non-linear registration | - | TRUE | FALSE | 260/390 | FALSE |
| Multisite - 94_Cryo_med_f | TRUE | - | Rigid registration | 2 | Non-linear registration | - | TRUE | FALSE | 266/400 | TRUE |
| Multisite - 94_Cryo_mediso_v | TRUE | - | Rigid registration | 2 | Non-linear registration | - | TRUE | TRUE | 240/360 | FALSE |
| Multisite - 94_RT_iso_v | TRUE | - | Rigid registration | 2 | Non-linear registration | - | TRUE | FALSE | 266/400 | FALSE |
| Multisite - 94_RT_mediso_f1 | TRUE | - | Rigid registration | 2 | Non-linear registration | - | TRUE | FALSE | 100/150 | FALSE |
| Multisite - 94_RT_mediso_f2 | TRUE | - | Rigid registration | 2 | Non-linear registration | - | TRUE | TRUE | 400/600 | TRUE |
| REST-AWK | FALSE | Rigid registration | Rigid registration | 2 | Non-linear registration | Non-linear registration | FALSE | FALSE | - | - |
| REST-AWK - bold-only | FALSE | Non-linear registration | Rigid registration | 2 | Non-linear registration | Non-linear registration | FALSE | FALSE | - | - |
| Montreal, Douglas CIC | FALSE | Non-linear registration | Rigid registration | 2 | Non-linear registration | Non-linear registration | FALSE | FALSE | - | - |
| Singapore, 3xTGAD | TRUE | - | Rigid registration | 2 | Non-linear registration | - | TRUE | FALSE | 120/180 | |
| Rat Guttingen | FALSE | Non-linear registration | Rigid registration | 2 | Non-linear registration | Non-linear registration | FALSE | FALSE | - | - |
| Rat NITRC | FALSE | No reg | Rigid registration | 3 | Non-linear registration | Non-linear registration | FALSE | FALSE | - | - |
| Rat CAMRI | TRUE | - | Rigid registration | 2 | Non-linear registration | - | FALSE | FALSE | - | - |

**Supplementary table 4:** Preprocessing parameters across datasets. The EPI - Otsu threshold and the EPI masking during unbiased template generation are discussed in **sup. table 3**.

| Category | Name | Definition |
|---|---|---|
| Nuisance regressor | 6 motion parameters | Corresponds to 3 rotations (Euler angles in radians) and 3 translations (in mm) measured for head motion realignment at each timeframe. Prior to the regression, the motion regressors are also subjected to the same frame censoring, detrending and frequency filtering which were applied to the BOLD timeseries to avoid the re-introduction of previously corrected confounds, as recommend in (Power et al., 2014) and (Lindquist et al., 2019). |
| | 24 motion parameters | Corresponds to the 6 motion parameters ($mot6$) together with their temporal derivatives, and 12 additional parameters are obtained by taking the squared terms (i.e. Friston 24 parameters (Friston et al., 1996)) $$mot24_t = [mot6_t, (mot6_t - mot6_{t-1}), (mot6_t)^2, (mot6_t - mot6_{t-1})^2]$$ with $mot24_t$ representing the list of 24 regressors for timepoint $t$. As with $mot_6$, the 24 regressors are additionally subjected to censoring, detrending and frequency filtering if applied on BOLD. |
| | WM/CSF/vascular /global signal | The mean signal is computed within the corresponding brain mask (white matter (WM), cerebrospinal fluid (CSF), vascular or whole-brain mask) from the partially cleaned timeseries (i.e. after frequency filtering, steps 1-4 in **methods section 5**). |
| | aCompCor - Percentage | Principal component timecourses are derived from timeseries within the combined WM and CSF masks (aCompCor technique (Muschelli et al., 2014)). From the timeseries within the WM/CSF masks $Y_{WM/CSF}$, a principal component analysis (PCA) decomposition is conducted to derive $$Y_{WM/CSF} = W_{aCompCor}C^T$$ with $C$ corresponding to a set of spatial principal components, and $W$ to their associated loadings across time. The set of first components explaining 50% of the variance are kept, and their loadings $W_{aCompCor}$ provide the set of aCompCor nuisance regressors. PCA is conducted on the partially cleaned timeseries (i.e. after frequency filtering, steps 1-4 in **methods section 5**). |
| | aCompCor - 5 components | aCompCor as defined above, but the first 5 components are kept instead of a set explaining 50% of the variance. |
| Temporal diagnostic features (**sup. figure 6**) | Power spectral density | The frequency power spectrum is computed for each voxel timecourse, and the average (and standard deviation) across voxels is displayed. |
| | Carpet plot | All timeseries are displayed into a 2D matrix format, where the rows are all the brain voxels and the columns are the timepoints. |
| | Translation and rotation parameters | Corresponds to 3 rotations (Euler angles in radians) and 3 translations (in mm) measured for head motion realignment at each timeframe. |
| | Framewise displacement (FD) | For each timepoint, corresponds to the displacement (mean across the brain voxels) between the current and the next frame. For each brain voxel within the referential space for head realignment (i.e. the 3D EPI which was provided as reference for realignment (**sup. table 1**)) and for each timepoint, the inverse transform of the head motion parameters (from the corresponding timepoint) is applied to obtain the voxel position pre-motion correction. Framewise displacement can then be computed for each voxel by computing the Euclidean distance between the positions pre-motion correction for the current and next timepoints. Thus, the mean framewise displacement $FD_t$ at timepoint $t$ is computed as $$FD_t = \frac{1}{n}\sum_{i=1}^{n}\sqrt{(x_{i,t+1} - x_{i,t})^2 + (y_{i,t+1} - y_{i,t})^2 + (z_{i,t+1} - z_{i,t})^2}$$ using the 3D $x$,$y$ and $z$ spatial coordinates in mm for timepoints $t$ and $t+1$ and for each voxel indices $i$. Framewise displacement for the last frame (which has no future timepoint) is set to 0. |
| | DVARS | Represents the estimation of temporal shifts in global signal at each timepoint, measured as the root-mean-square of the timeseries' temporal derivative |

| | | |
|---|---|---|
| | | $$DVARS_t = \sqrt{\frac{1}{n}\sum_{i=1}^{n}(Y_{i,t} - Y_{i,t-1})^2}$$ where $Y_{i,t}$ corresponds to the BOLD signal in brain voxel $i$ at timepoint $t$. The first timepoint is set to 0 (has no previous timepoint). |
| | Edge/WM/CSF mask | The L2-norm across voxels within a mask, at each timepoint. |
| | $CR_{var}$ | The variance estimated by confound regression is computed for each timepoint. This is done by taking the L2-norm $CR_{var} = \|\|Y_{CR}\|\|_2$ across voxels at each timepoints, where $Y_{CR}$ is the predicted confound timeseries (see **methods section 5**). |
| | CR $R^2$ | Represents the proportion of variance explained (and removed) by confound regression. This is obtained with $CR_{R^2} = 1 - \frac{Var(\hat{Y})}{Var(Y)}$ at each timepoint, where $Y$ and $\hat{Y}$ are the timeseries pre- and post-regression respectively, and $Var(x) = \frac{1}{n}\sum_{i=1}^{n}(x_i - \mu_x)^2$ calculates the variance, with $\mu$ as the mean. |
| | Mean amplitude | A set of timecourse are averaged as $\frac{1}{n}\sum_{i=1}^{n}\|X_i\|$, where $X_i$ is the timecourse $i$. Timecourses can correspond to either of the following sets:<br>● **DR confounds:** timecourses from the first stage of dual regression, using confound components (**sup. figure 5**).<br>● **DR networks:** network timecourses from the first stage of dual regression.<br>● **SBC networks:** network timecourses derived from the set of seeds provided. |
| Spatial diagnostic features (**sup. figure 6**) | $BOLD_{SD}$ | The temporal standard deviation is computed for each voxel from the BOLD timeseries. |
| | $CR_{SD}$ | The temporal standard deviation computed on each voxel from the predicted confound timeseries during confound regression (i.e. $Y_{CR}$, **methods section 5**). |
| | CR $R^2$ | The proportion of variance explained by confound regression at each voxel. This is obtained with $CR_{R^2} = 1 - \frac{Var(\hat{Y})}{Var(Y)}$ at each voxel, where $Y$ and $\hat{Y}$ are the timeseries pre- and post-regression respectively, and $Var(x) = \frac{1}{n}\sum_{i=1}^{n}(x_i - \mu_x)^2$ is variance of $x$, with $\mu$ as the mean. |
| | Global signal covariance ($GS_{cov}$) | The covariance between the global signal and the timeseries at each voxel is measured as $GS_{cov} = \frac{1}{n}\sum_{t=1}^{n} Y_t \times GS_t$, where $GS_t = \frac{1}{n}\sum_{i=1}^{n} Y_i$, i.e. the mean across all brain voxels for a given timepoint. |
| | DR network X | The linear coefficients resulting from the second regression with dual regression, corresponding to a network amplitude map (for the Xth network specified for analysis). |
| | SBC network X | The voxelwise correlation coefficients (pearson's r) estimated with seed-based connectivity (for the Xth seed provided for analysis). |
| Scan-level quality control metrics (**sup. figure 9**) | Network amplitude | The overall network amplitude is summarized by computing the L2-norm of the dual regression network amplitude map of linear coefficients. |
| | Network specificity | The network map (seed-based or dual regression) and the corresponding canonical network map are thresholded to include the top 4% of voxels with highest connectivity, and the overlap of the thresholded area is computed using Dice overlap. |
| | Dual regression confound correlation | The timecourse for a single network (from a seed or dual regression) is correlated with the timecourse from each confound component modelled through dual regression, then the absolute mean correlation is computed to obtain the average amplitude of confound correlations for this specific network analysis. |

| | | |
|---|---|---|
| | Total $CR_{SD}$ | The global standard deviation of the 4D confound timeseries (i.e. $Y_{CR}$, **methods section 5**). |
| | Mean framewise displacement | The mean framewise displacement computed across time (only including frames after censoring applied for confound correction). |
| | Temporal degrees of freedom | The remaining degrees of freedom post-confound correction are calculated as $tDOF = Original\ number\ of\ timepoints - Number\ of\ censored\ timepoints - Number\ of\ AROMA\ components\ removed - Number\ of\ regressors$ |

**Supplementary table 5:** Definition of metrics computed within RABIES for nuisance regression, spatiotemporal diagnosis, or quality control metrics computed at the scan level.

| Quality control metric | Importance | Limitation |
|---|---|---|
| Scan-level network specificity | Evaluates that the spatial features of the network are appropriately recovered in each scan. Accounts for network detectability issues, and spurious effects on network shape. | Does not detect spurious effects that only influence network amplitude (and not shape), and high scan-level network specificity does not imply specific network variability (**sup. figure 11**). |
| Scan-level temporal correlation with confounds | Controls for spurious connectivity, including network amplitude effects, at the scan level. In the case of strong confounding effects driving network amplitude with little impact on network shape, this measure can best capture spurious connectivity (**sup. figure 12**). | ● The selection of confound components is subjective (see classification in **methods section 10**), and requires expertise to conduct reliably.<br>● If network activity is predominant in a given scan with little confounds, dual regression can overfit confound components to network activity, providing false confound correlations (**sup. figure 13**). This cautions against interpreting low correlation values, and restricts the application of this metric to the exclusion of highly corrupted scans.<br>● Since lowpass filtering increases temporal autocorrelation (Ebisuzaki, 1997), defining an accurate inclusion threshold is more challenging when combined with this filtering technique. |
| Group-level specificity of network variability | Meeting the scan-level network specificity thresholds is insufficient to ensure that cross-subject variability is specific to the network, as inter-scan variability can differ significantly from the core network features observed in individual scans (**sup. figure 11**). | Divergences in network detectability between subjects as well as spurious effects on network amplitude with minimal impact on shape can both drive cross-scan variability in a network-specific manner. Thus, to ensure that network variability reflects biological variability of interest instead of divergences in data quality, it must be complemented with other metrics accounting for those effects. |
| Group-level confound correlation | Detects residual relationships between inter-scan variability in connectivity and confounds which can remain after accounting for scan-level thresholds (**sup. figure 14**). | ● Group-level correlations do not necessarily capture large confound effects observable in individual scans (**sup. figure 12**).<br>● What constitutes a 'concerning' correlation size should be considered in the context of the study, depending on the effect size of interest (i.e. is the effect size of interest much higher or similar to the effect size of confounds?). It is difficult to establish a standardized correlation threshold, as effect sizes can be inflated in low sample size (Marek et al., 2022).<br>● The $CR_{SD}$ measure of confound will depend on the input parameters for computing confound regression, and the variance explained by confound regression is influenced by baseline random noise (e.g. scanner thermal noise), and thus may be influenced by factors which are not strickly motion or physiological confounds. |

**Supplementary table 6:** Complementarity and limitations of the selected set of quality control metrics.

### SUPPLEMENTARY FIGURES

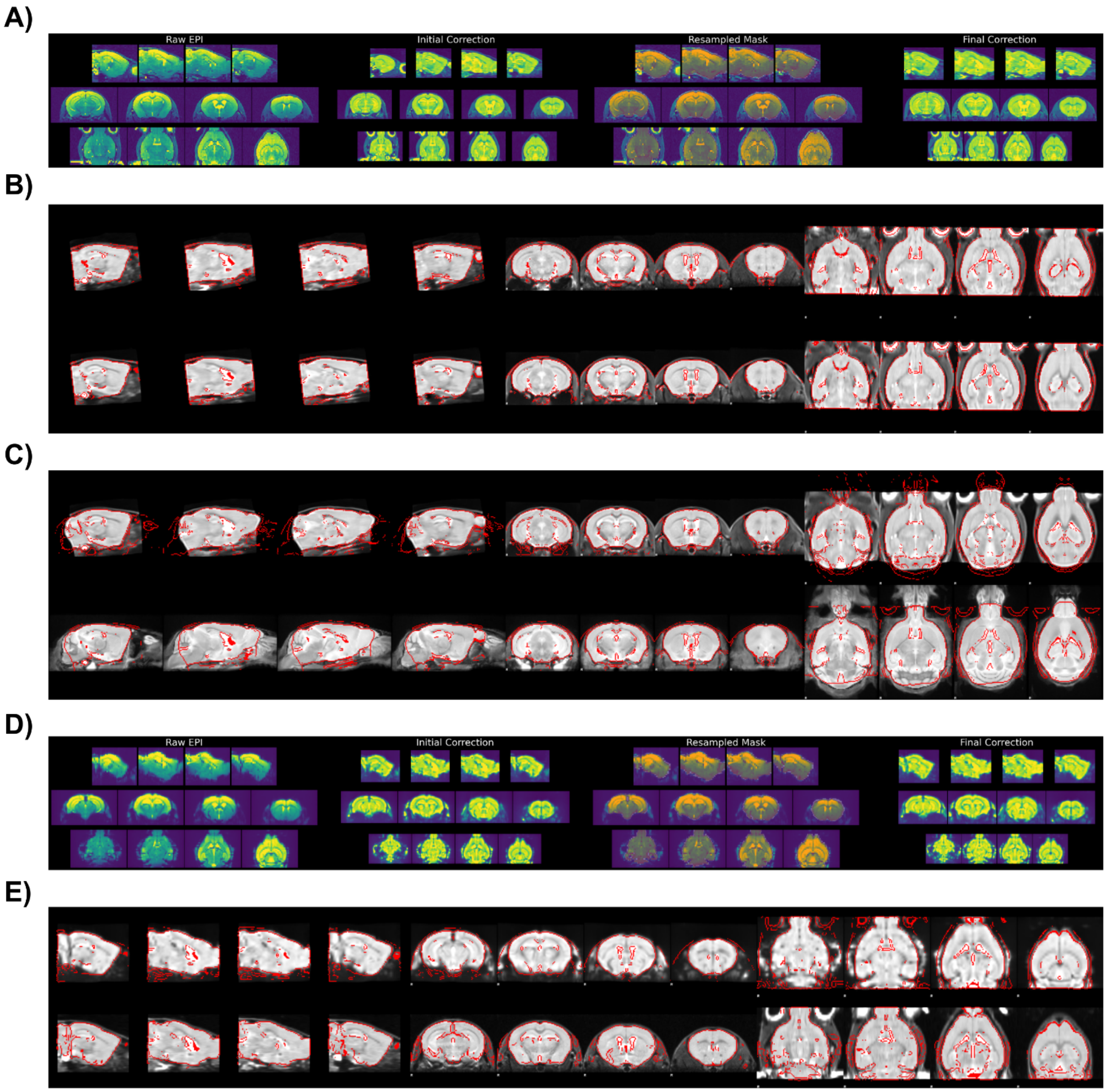


**Supplementary figure 1:** Examples of automatically-generated outputs for preprocessing quality control. **A)** The masking of the anatomical scan for inhomogeneity correction. The four columns show 1) the raw image before correction, 2) the image after an initial correction prior to registration, 3) the mask brain mask obtained for the final correction and 4) the final inhomogeneity correction. The sagittal, coronal and transverse slices are shown from top to bottom for each column. **B)** Quality control for the registration of the inhomogeneity-corrected anatomical scan (top row) to the dataset-generated unbiased average (bottom row). Anatomical edges were automatically detected using FSL's slicer (https://fsl.fmrib.ox.ac.uk/fsl/fslwiki/Miscvis) and overlapped over the opposite image to allow the evaluation of anatomical correspondence post-registration. **C)** Same as B), for the unbiased average template (top) and the external atlas (bottom). **D)** same as A), but for the EPI image. **E)** Same as B), but for the inhomogeneity-corrected EPI (top) and the inhomogeneity-corrected anatomical scan (bottom) from the same scanning session.

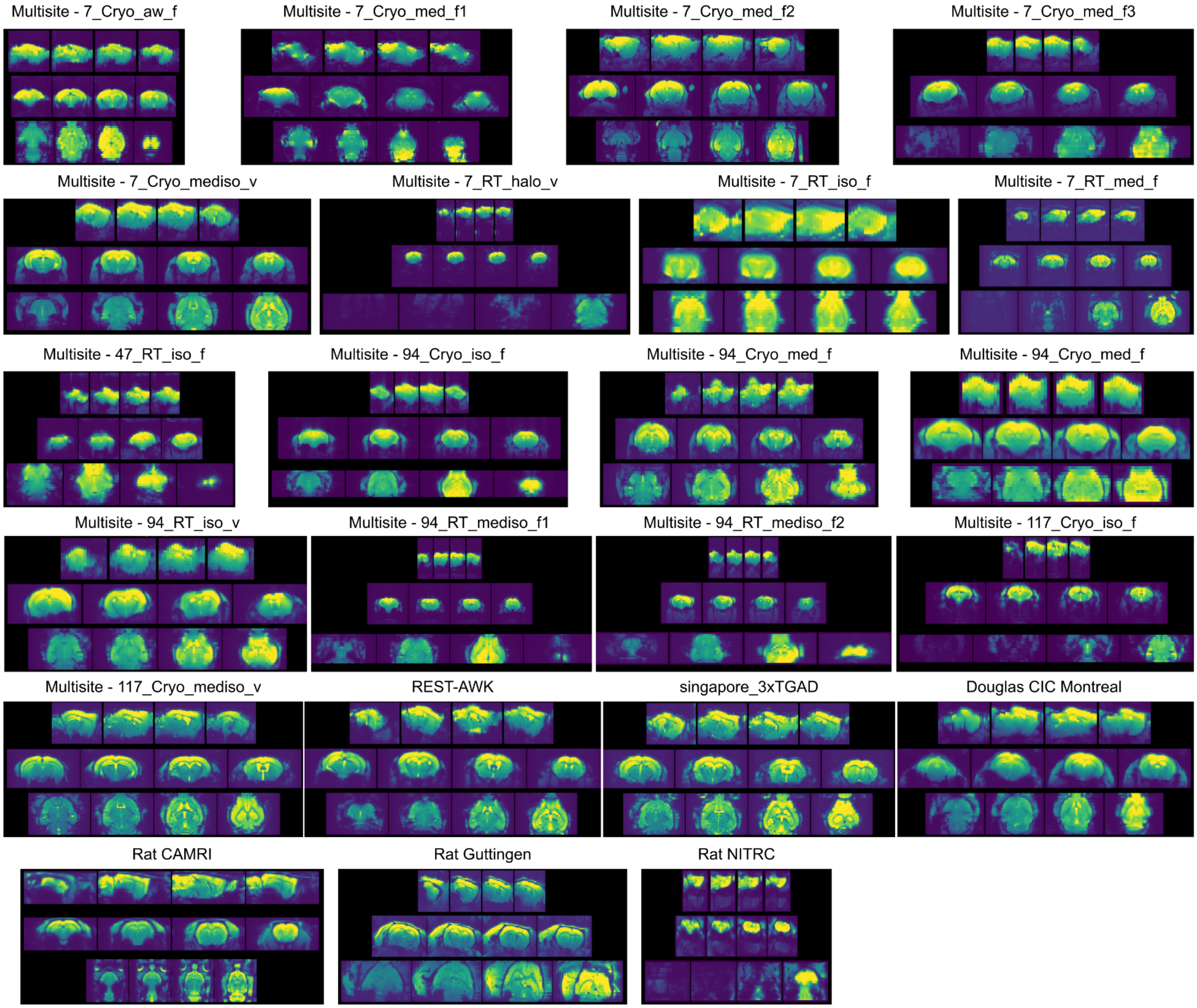


**Supplementary figure 2:** One example raw EPI scan is shown for each of the 23 different rodent fMRI dataset that were used for benchmarking the robustness of preprocessing quality. For each example, slices across the sagittal, coronal and transverse planes are displayed from the top to the bottom rows respectively. Datasets vary substantially in terms of anatomical contrast, intensity inhomogeneities, susceptibility distortions, brain coverage and in the presence of external tissue or ghosting artefacts.

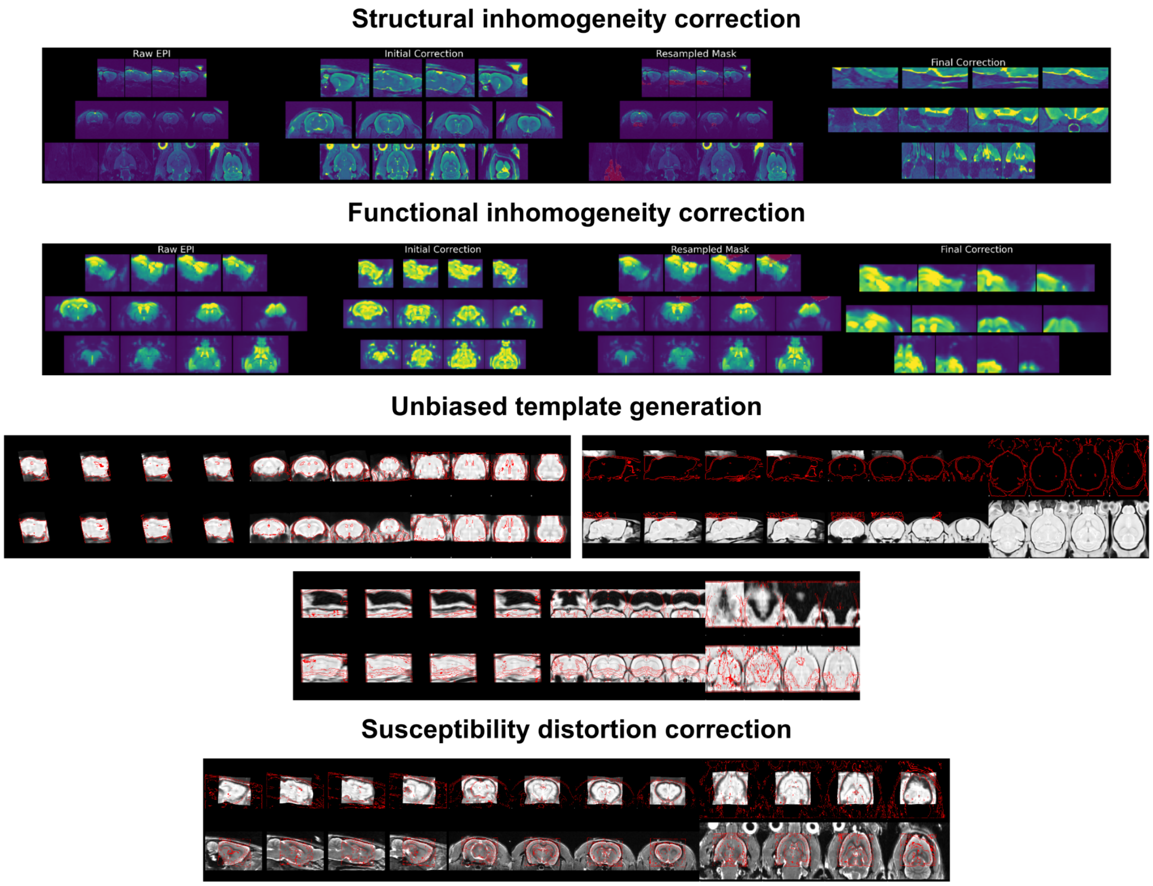


**Supplementary figure 3:** all the failures identified during preprocessing quality control.

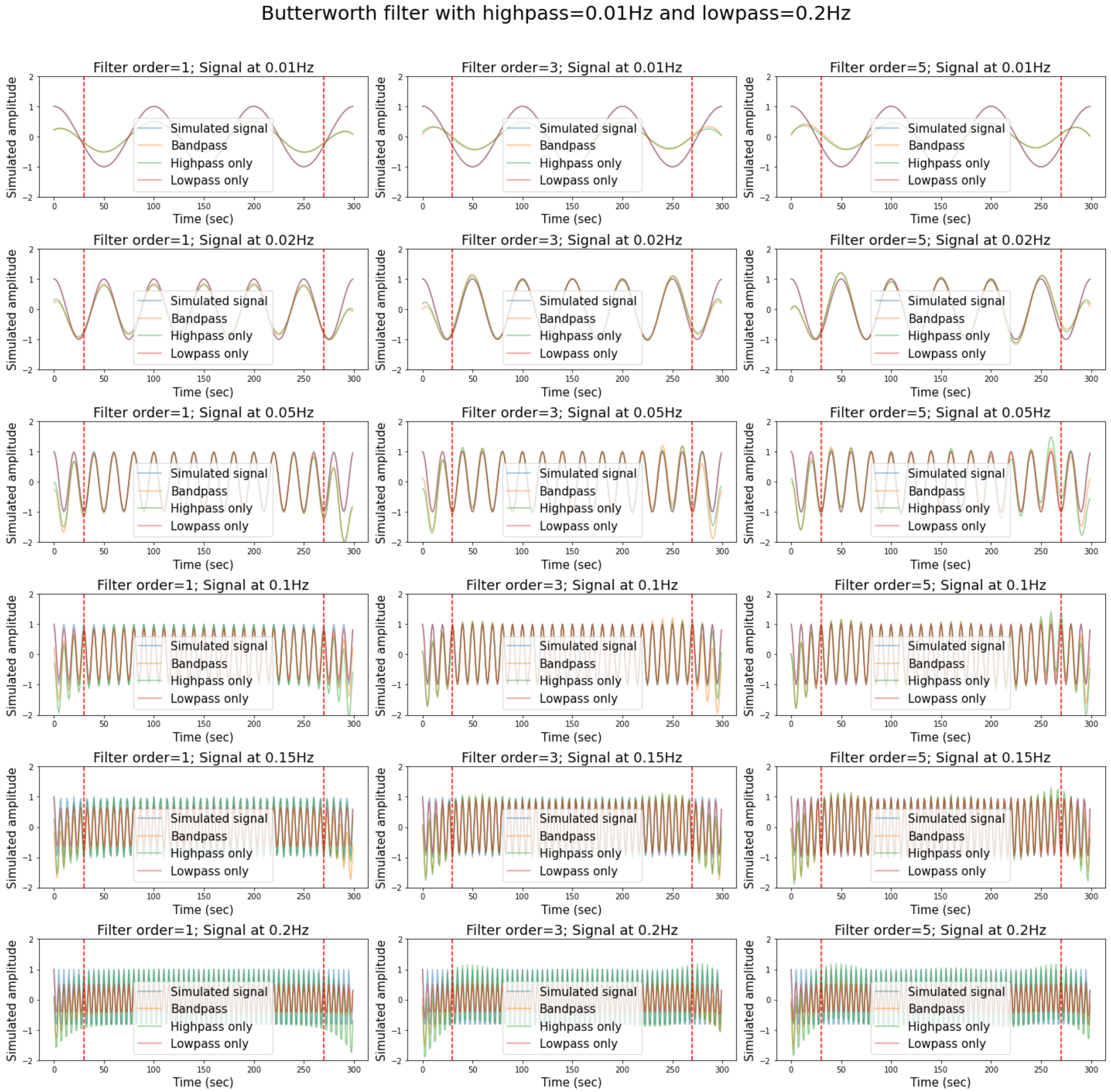


**Supplementary figure 4:** Visualizing the impact of butterworth bandpass filters on simulated data. For each simulation, a cosine wave (sampled at 1Hz for 300 seconds) is generated at a given frequency (0.01Hz, 0.02Hz, 0.05Hz, 0.1Hz, 0.15Hz, or 0.2Hz), then either highpass at 0.01Hz, lowpass at 0.2Hz or bandpass at 0.01Hz-0.2Hz is applied using a butterworth filter with a given filter order (1$^{st}$, 3$^{rd}$ or 5$^{th}$ order). The dotted red lines represent the selected time cutoff of 30 seconds at each edge to remove edge artefacts.

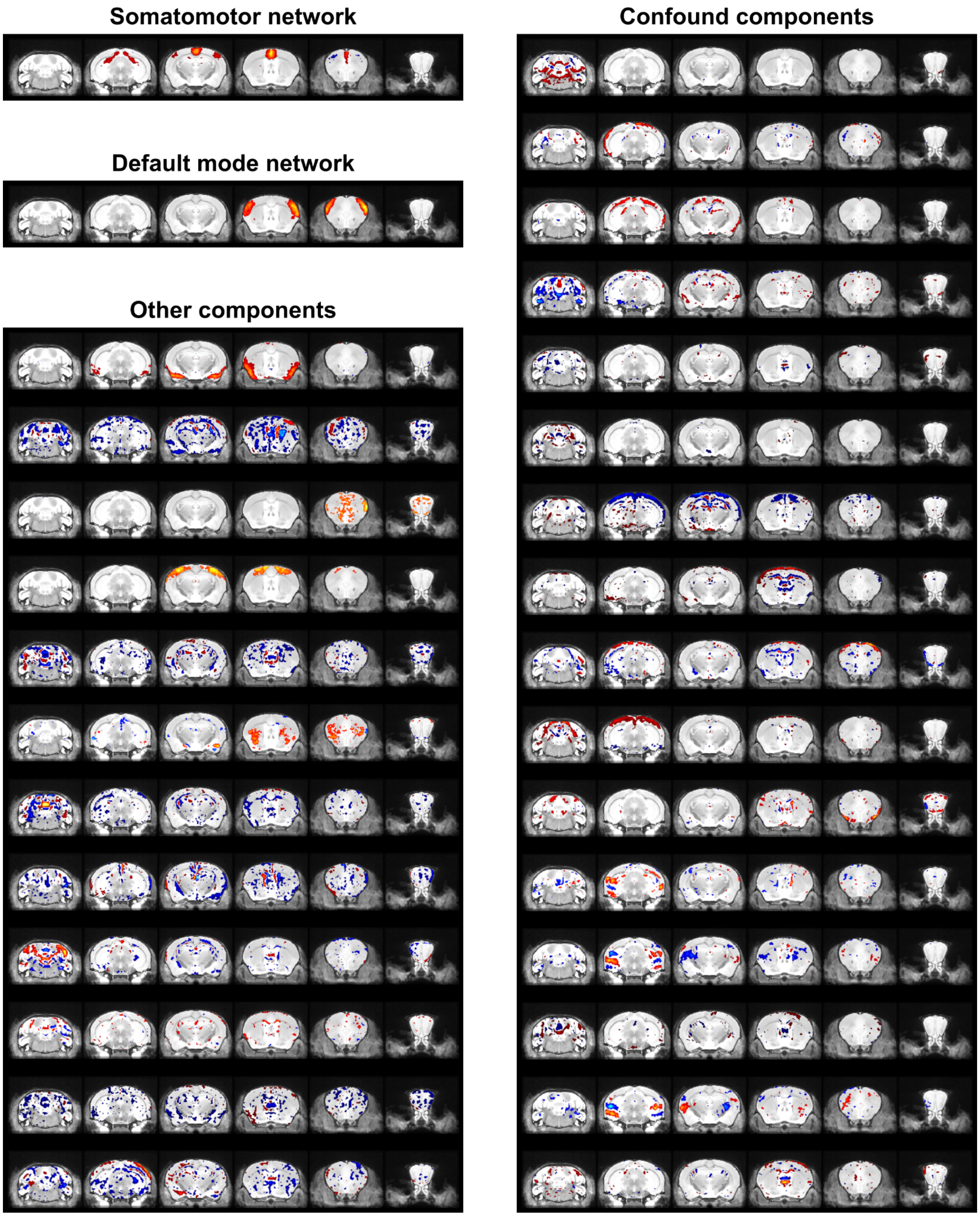


**Supplementary figure 5:** Components derived from group-ICA decomposition on the REST-AWK dataset using 30 components. The dataset was corrected for confounds prior to group-ICA using FD and DVARS censoring, highpass filtering at 0.01hz and confound regression using the 6 rigid motion parameters together with WM/CSF mask signals. The components were manually classified upon visual inspections. Each brain map was thresholded to include the top 4 % of voxels with highest component weights and is displayed on the coronal plane. The nifti file with the ICA components can be accessed online (https://zenodo.org/record/5118030/files/melodic_IC.nii.gz).

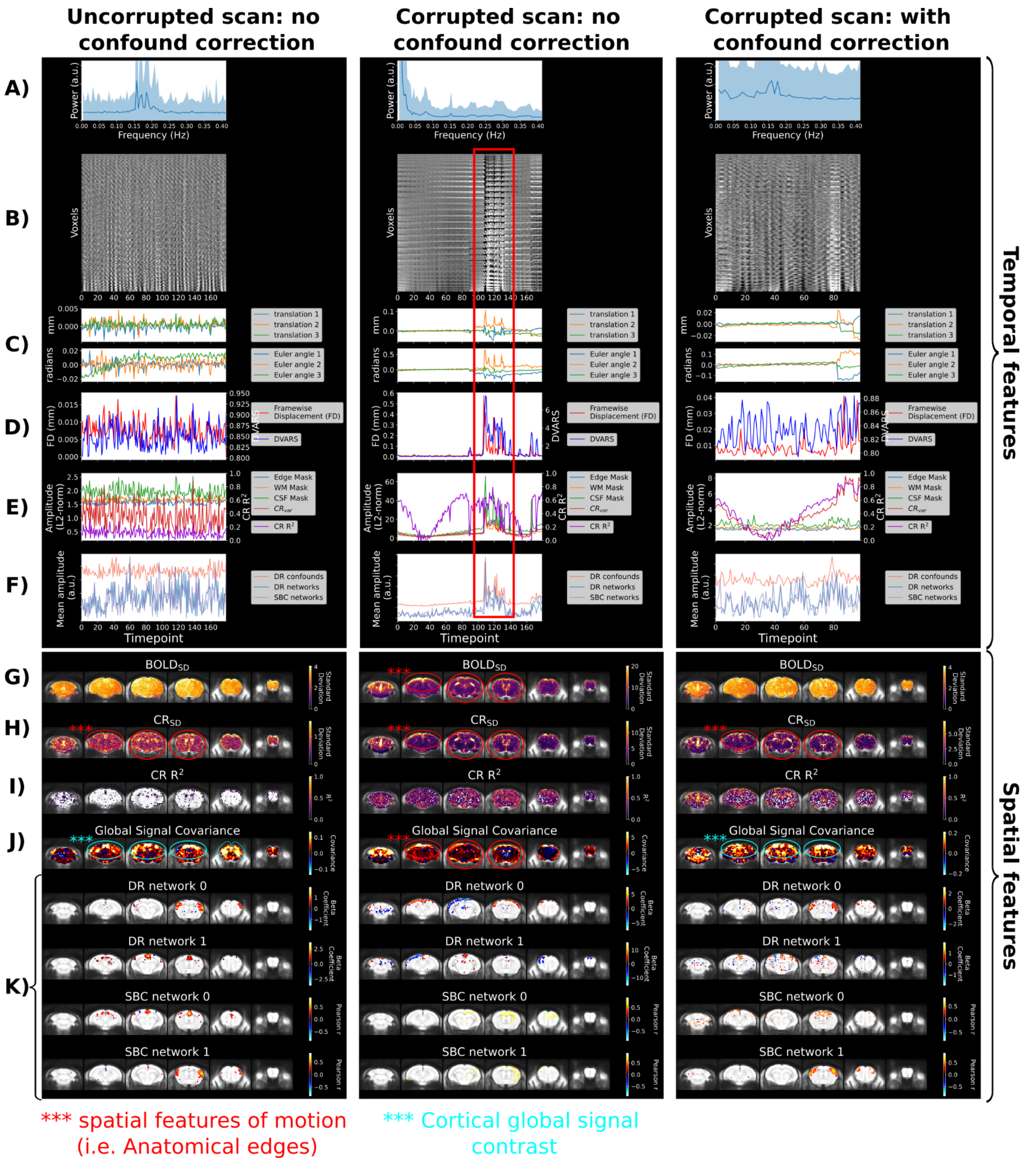


**Supplementary figure 6:** The spatiotemporal diagnosis generated by RABIES for 3 exemplified cases: an uncorrupted scan with no confound correction applied (first column), a motion-corrupted scan without confound correction (second column) and the same corrupted scan after applying confound correction (third column). The confound correction for the third example consisted of frame censoring using framewise displacement and DVARS combined, followed by confound regression of the 6 motion realignment parameters. For each diagnosis, the top half represents the set of temporal features, and the bottom half the set of spatial features. Each spatial map is represented along 6 cortical slices, overlapped onto the anatomical template in common space. The network maps from dual regression (DR) or seed-based connectivity (SBC) are thresholded to include the top 4% of the voxels with the highest values. In this report, DR network 0 and SBC network 1 correspond to the somatomotor network, whereas DR network 1 and SBC network 0 correspond to the default mode network. Each aspect of the spatiotemporal diagnosis is discussed in **sup. material section 1** and metrics are defined in **sup. table 5**.

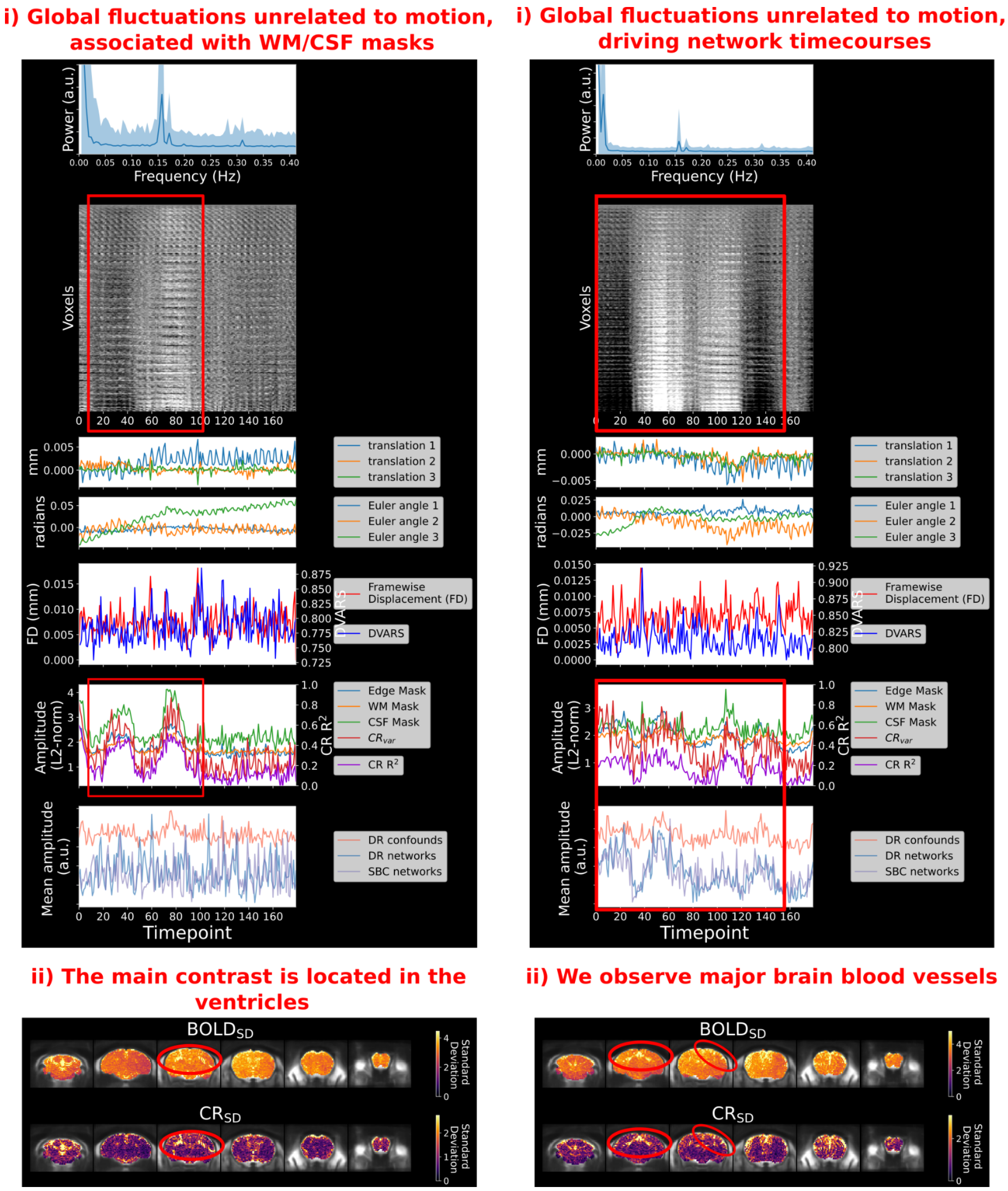


**Supplementary figure 7:** The spatiotemporal diagnosis can identify confounds of physiological origins. **A)** This scan displays slow global signal fluctuations which are minimally affecting the network timecourses. With the spatial maps of $BOLD_{SD}$ and $CR_{SD}$, we can observe that signal fluctuations are most pronounced over the ventricles. **B)** This scan also shows slow global signal fluctuations, but here the network timecourses are driven by the confound. The spatial maps of $BOLD_{SD}$ and $CR_{SD}$ reveal that signal fluctuations are strongest over major brain vessels, including the sagittal sinus and indications of cortical penetrating vessels. Both scans were not subjected to any confound correction for this figure.

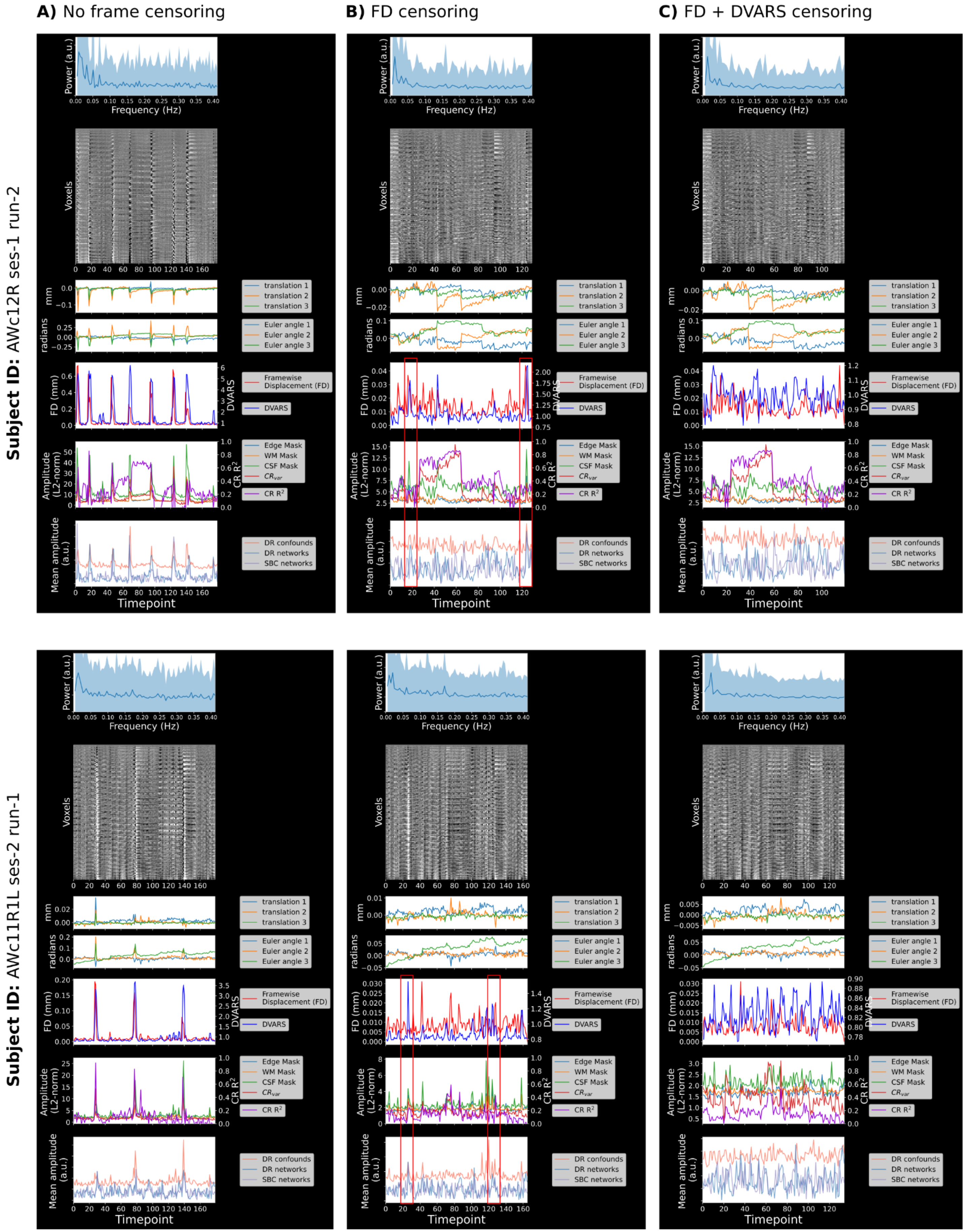


**Supplementary figure 8:** Two example scans where there are residual spike confounds even after framewise displacement (FD) censoring. For two different subjects (top and bottom rows), the temporal diagnoses are shown **A)** without applying any censoring, **B)** with FD censoring and **C)** with both FD and DVARS censoring. In each case the 6 motion parameters were also regressed. For each example, we can observe that when only FD censoring is applied, residual DVARS spikes are correlated with CSF signal and the dual regression confound components, as well as the network timecourses, suggesting corruption of network fits. These confounds are removed by the additional application of DVARS spike censoring.

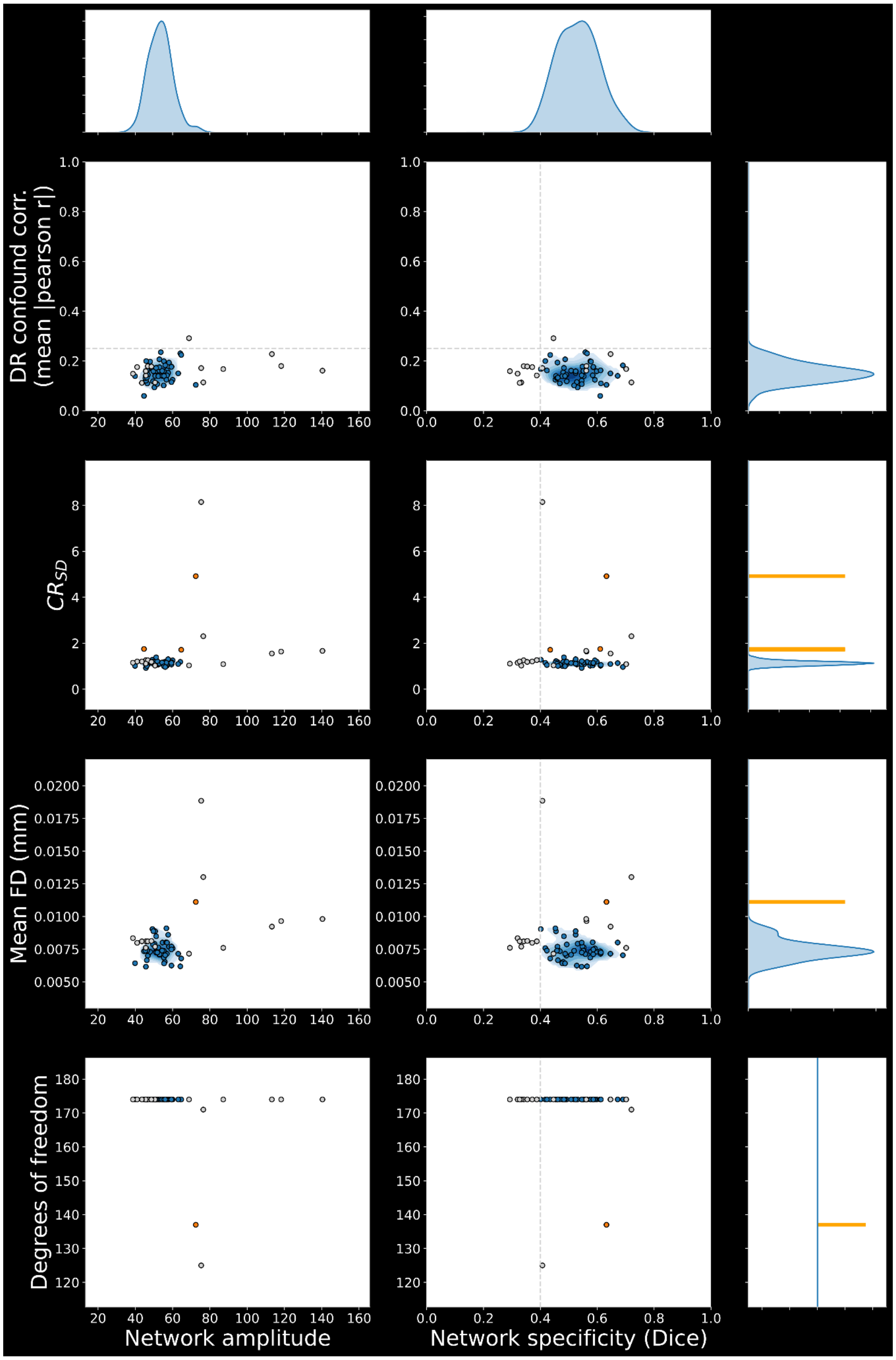


**Supplementary figure 9:** Example of a dataset distribution report (somatomotor network from the REST-AWK mediso dataset). Measures of network specificity and amplitude are contrasted with measures of confounds across samples (each point is a scan). Scan-level thresholds (**figure 3**) and removal of outliers in network amplitude (**methods section 14**) was applied, and the removed scans are shown in gray. For each metric separately, remaining outliers are highlighted in orange, whereas the dataset distribution is estimated from the remaining set of scans in blue. Scan-level thresholds for network specificity and dual regression (DR) confound correlation are shown as gray dotted lines along their respective axes. The derivation for each measure is detailed in **sup. table 5**.

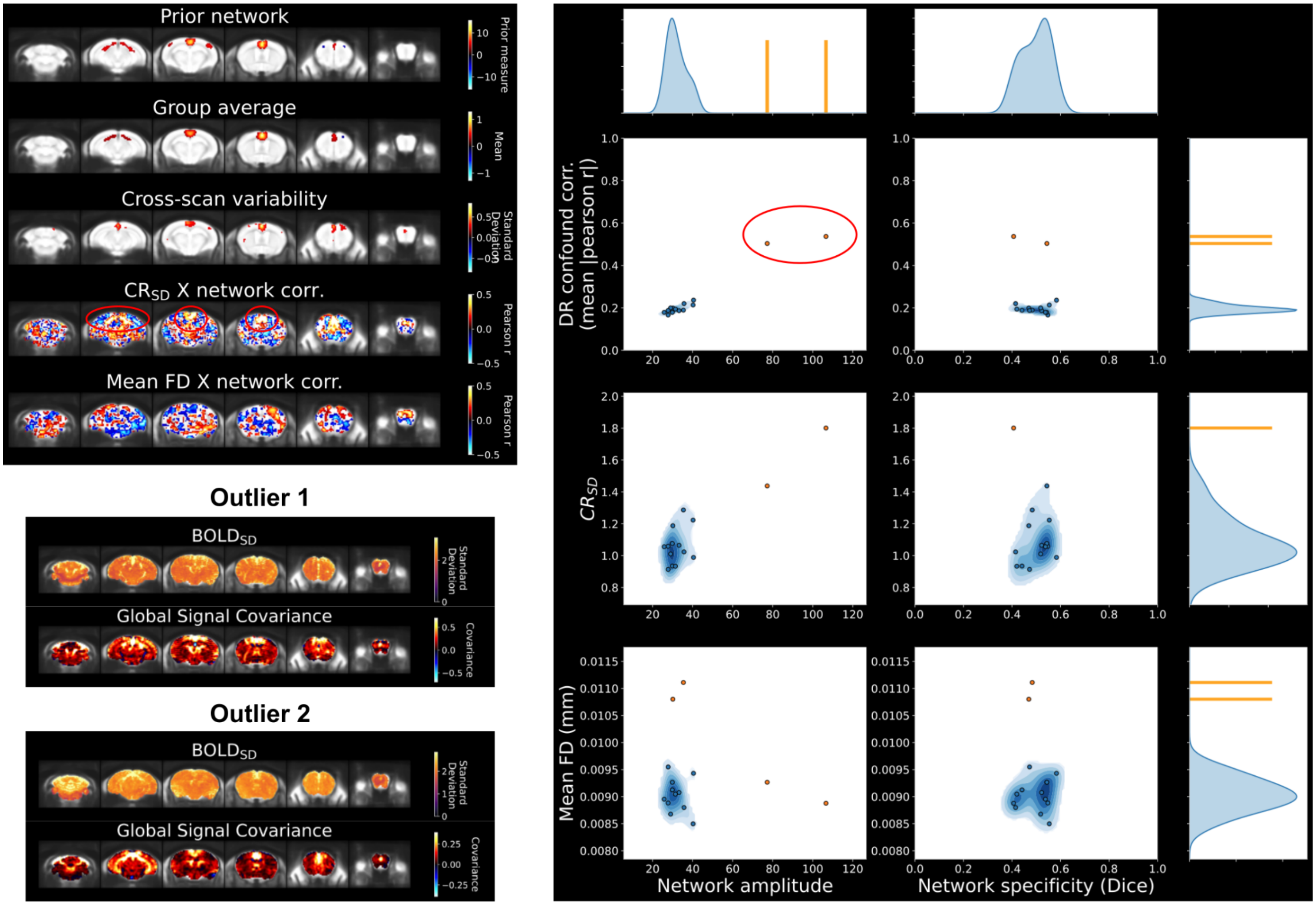


**Supplementary figure 10:** In an analysis of the default mode network with the 7_Cryo_mediso_v dataset, a group-wise correlation with the $CR_{SD}$ measure of confounds (top left report) is explained by two outliers with spurious network amplitude (see distribution report on the right). Inspecting the spatiotemporal diagnosis then allows to identify the presence of a vascular confound for both outliers (shown on bottom left), reproducing the anatomical signature of major brain vessels as observed in both the $BOLD_{SD}$ and global signal covariance map.

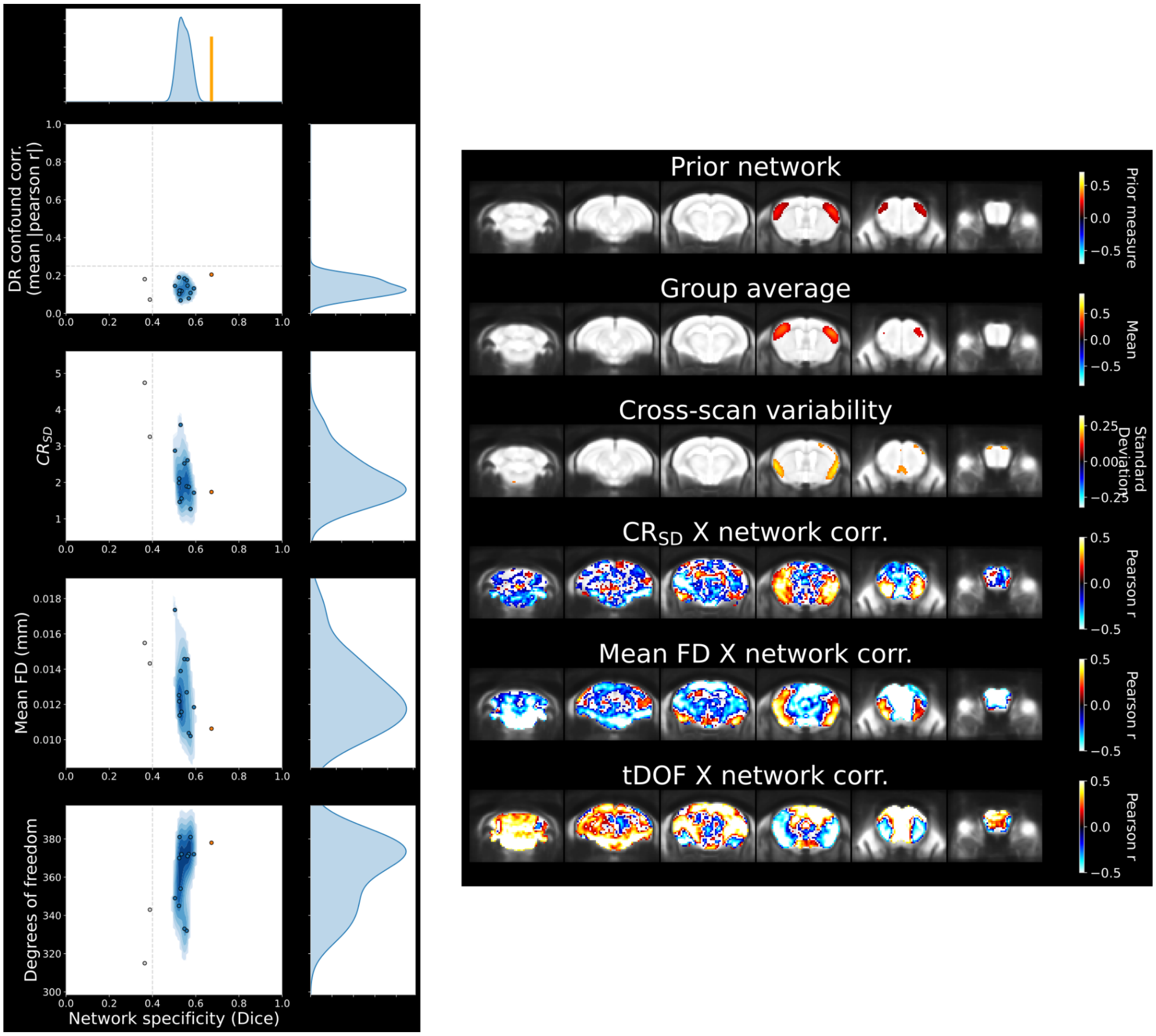


**Supplementary figure 11:** The group-level network variability can capture spurious connectivity which would not be detected otherwise. An example is shown for the 7_Cryo_aw_f dataset analysis of the somatomotor network with seed-based connectivity. Despite passing both scan-level thresholds (distribution report on the left), cross-scan variability is mainly driven by a spurious signature in ventral areas below the network (group statistical report on the right). Although confound correlations also high in this ventral area, the highest average correlation evaluated within the network is of 0.07 with $CR_{SD}$. Thus, assessing the specificity of network variability is crucial to detecting spurious effects.

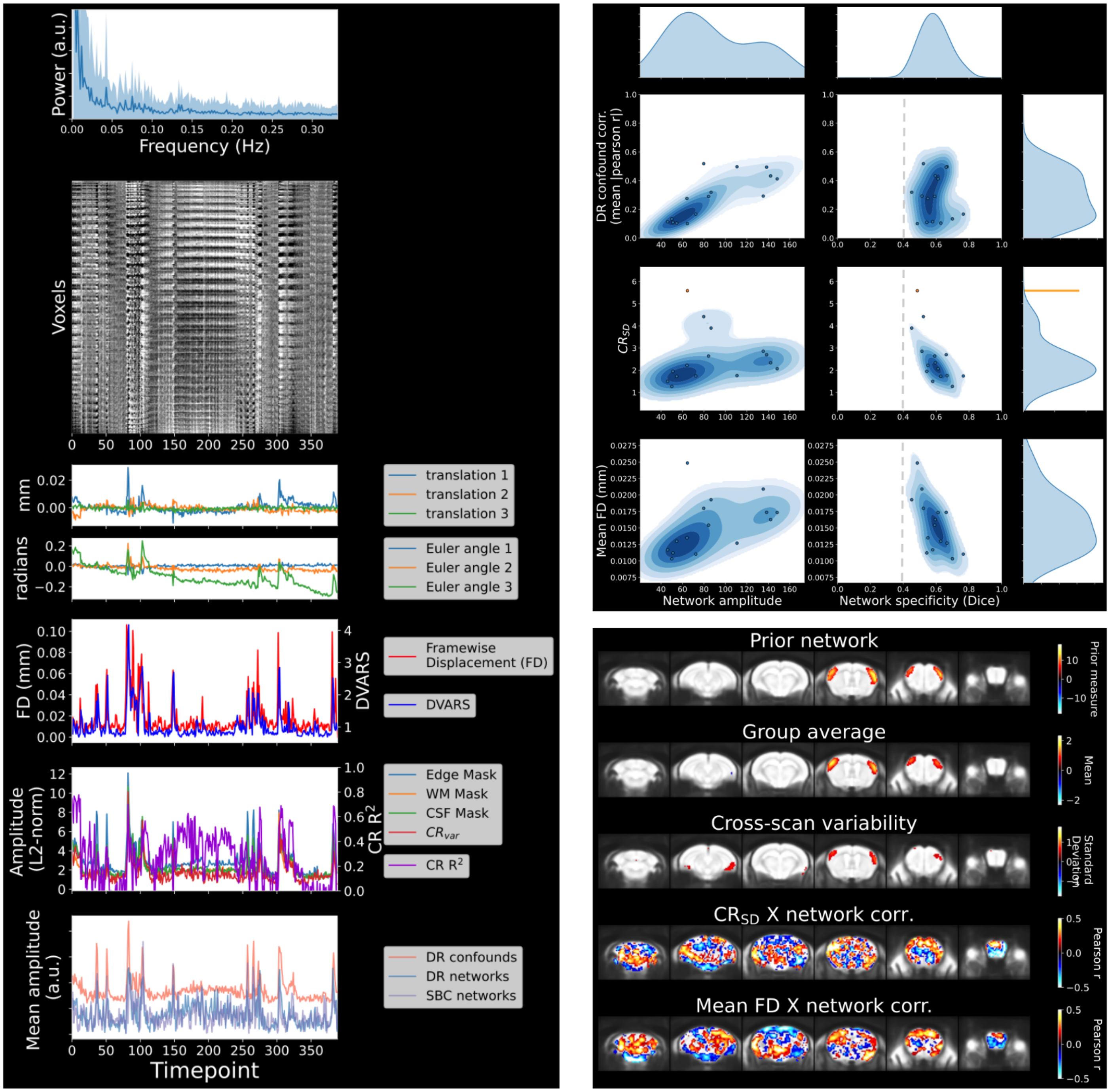


**Supplementary figure 12:** Example where spurious connectivity is only well assessed from scan-level confound correlation (7_Cryo_aw_f dataset with no confound correction applied). Inspecting the temporal features of individual scans reveals strong impact of framewise displacement on network connectivity in many scans (one example shown on the left). With scan-level measures (distribution report on the top right), we can assess that network amplitude is clearly impacted (high DR confound correlation associated with higher network amplitude), but with minimal impact on network shape (network specificity is above threshold for all scans). The group statistical report (on the bottom right) does not capture well the extent of the spurious effects, as cross-scan variability reproduces network features and confound correlations are relatively low (average correlation of 0.22 for $CR_{SD}$ and 0.17 for mean FD).

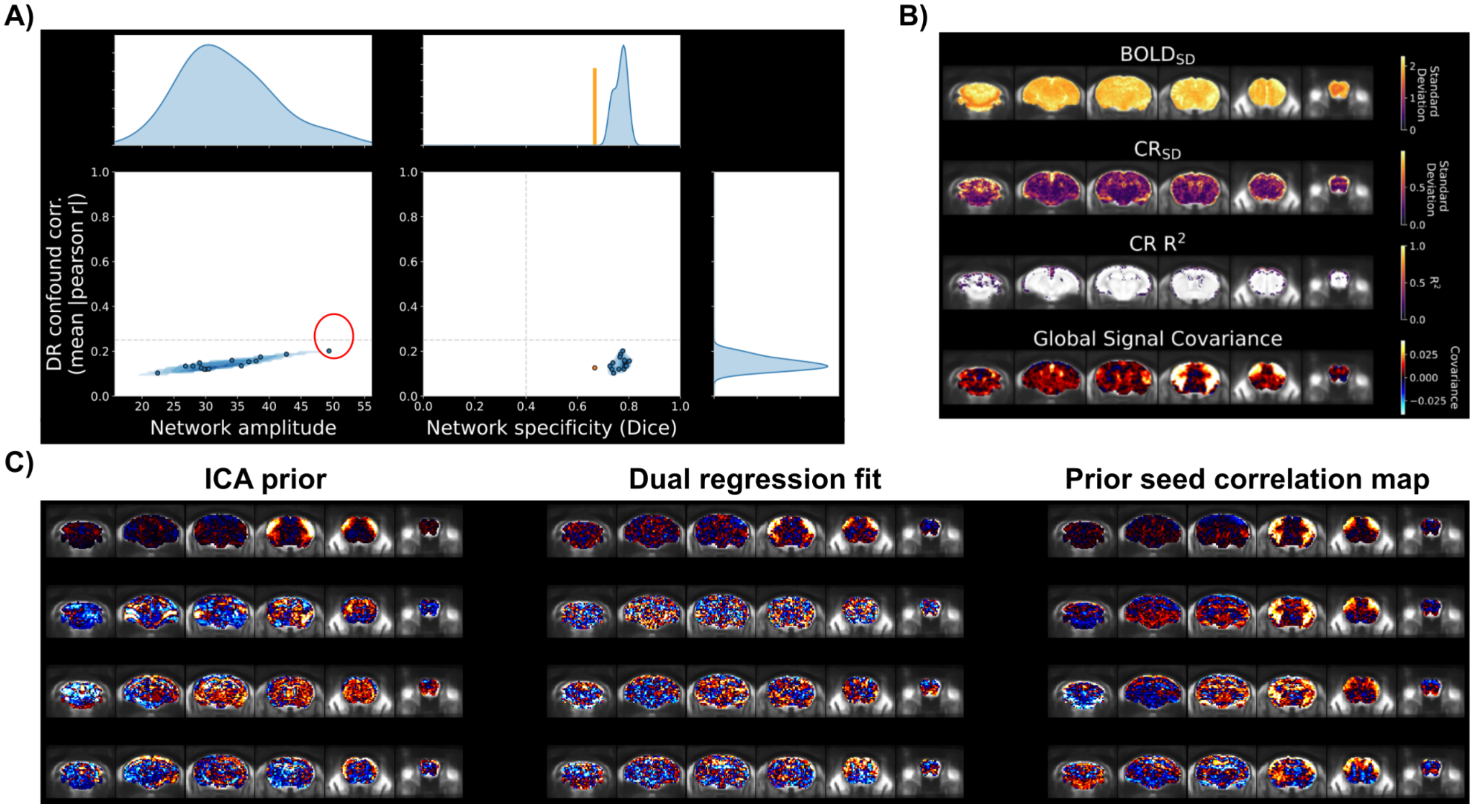


**Supplementary figure 13:** Scan-level confound correlation can overfit to network activity at low correlations. **A)** The dataset distribution report from the 7_Cryo_mediso_v dataset after confound correction optimization for the somatomotor network. Scan-level confound correlation is low, but is linearly associated with network amplitude across scans (correlation of 0.9). **B)** Spatial features from the scan with the highest confound correlation. (circled in red in **A)**) The global signal contrast is clearly predominantly associated with the somatomotor network. **C)** Demonstration of dual regression overfitting in this subject. On the left, the ICA component maps for the somatomotor network is shown together with the three confound components which had highest temporal correlation with the network. In the middle column, the spatial maps derived from the dual regression are shown for each corresponding ICA prior. In the column on the right, the timecourse from each component was correlated voxelwise (as if conducting seed-based connectivity), and the resulting correlation maps are shown. All confound components produce a correlation map which reproduces features of the somatomotor network. This demonstrates a low specificity of the dual regression fits for these confound components, which more likely driven (at least partially) by the predominant network activity.

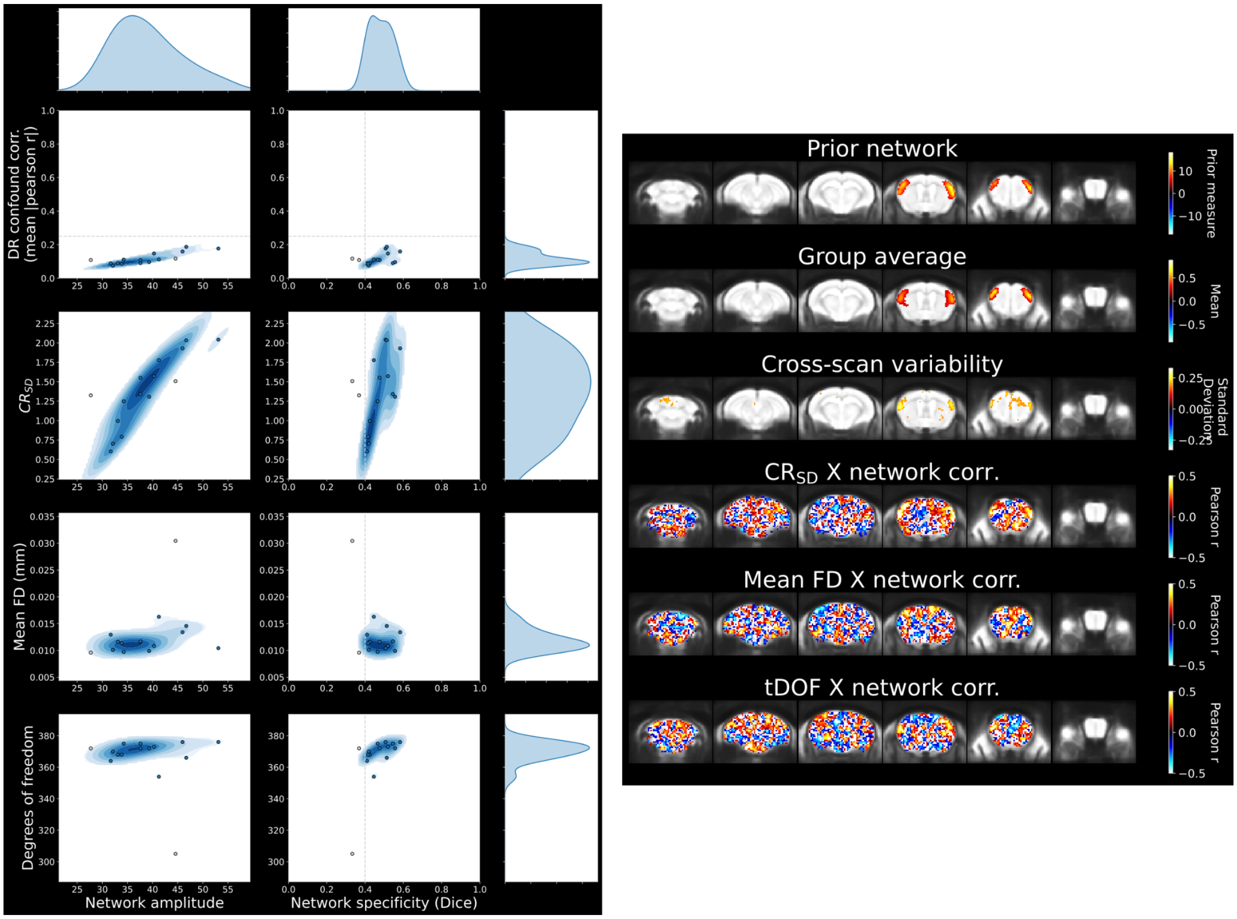


**Supplementary figure 14:** Group-level confound correlation can reveal subtle effects which are not well captured otherwise. This is observed in the 94_Cryo_med_f dataset after optimizing confound correction. The group statistics and distribution reports for this example are shown. All quality control criteria are respected, except for a moderate correlation with confounds as detected through $CR_{SD}$ (average group correlation of 0.3 within the area of the network).

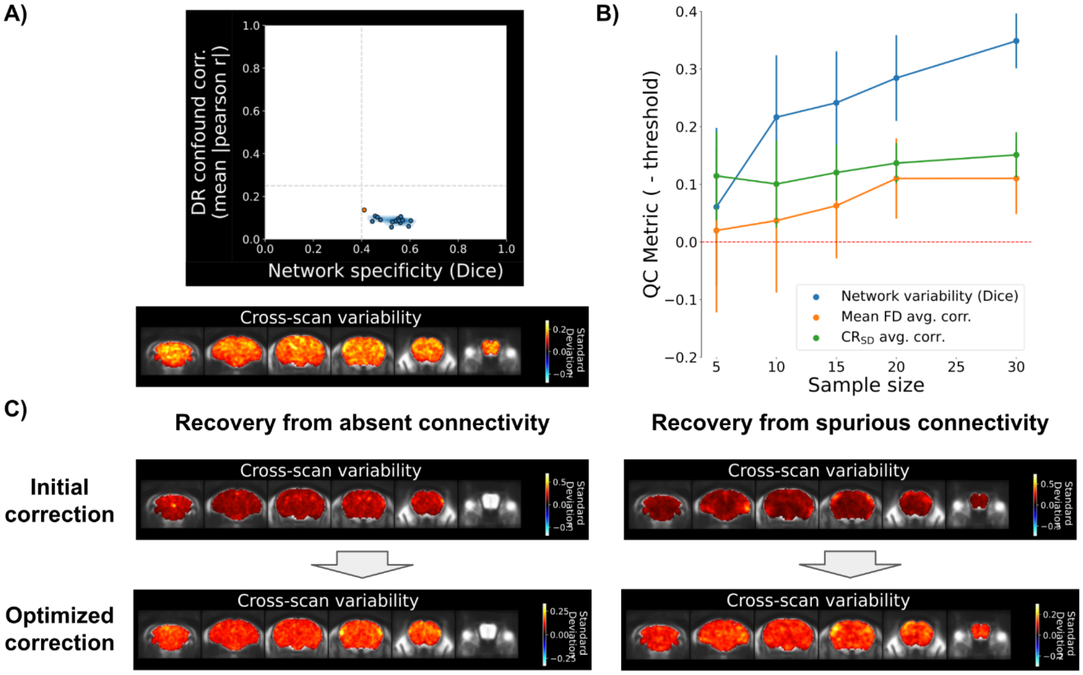


**Supplementary figure 15: A)** Example of a case where network variability was not well defined (Dice overlap of 0.2 with the canonical network) despite sufficient scan-level network detectability (dual regression analysis of the somatomotor network with the 7_RT_halo_v dataset after optimization for confound correction). **B)** The effect of sample size was evaluated on each quality metric from the group statistical report in the REST-AWK anesthetized (mediso) group (tDOF was not considered, since there was negligible variation in this dataset). The dataset was randomly subsampled at varying sample sizes (5, 10, 15, 20 and 30) for 50 iterations at each sample size, and the analysis quality report was computed on each subsample. Each point corresponds to the average across the 50 iterations and its associated error bar (standard deviation). **C)** Two examples where network variability was recovered by appropriate confound correction (94_Cryo_med_f on the left, 7_Cryo_aw_f on the right).

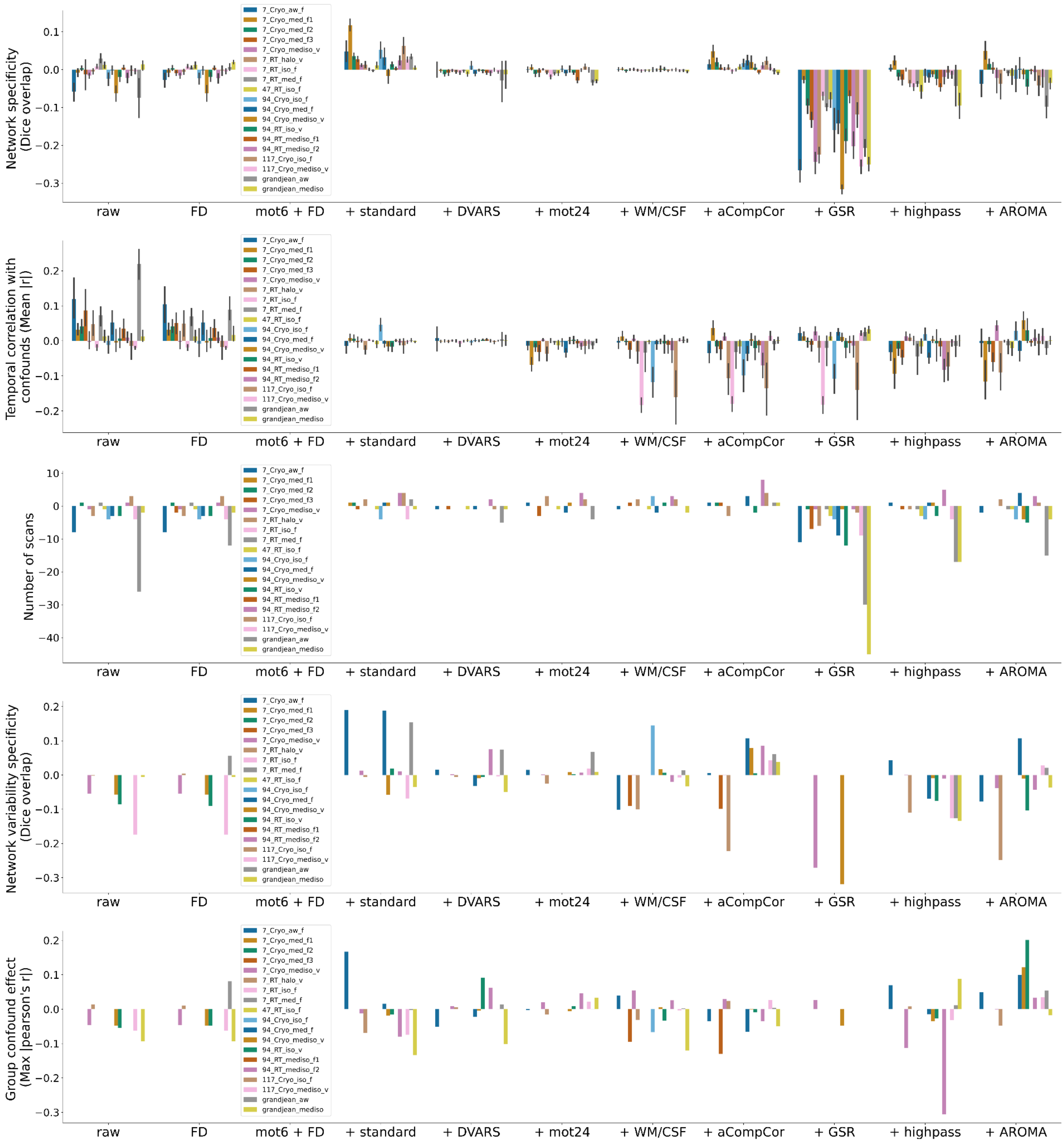


**Supplementary figure 16:** Impact of varying confound correction on quality control measures across datasets. For each dataset, measures include the scan-level assessment of network specificity and confound correlation (**figure 3**), the number of scans which passed those two thresholds, and at the dataset-level, the specificity network variability (Dice overlap with the canonical network maps), and the highest mean confound correlation within the area of the network obtained with either mean framewise displacement, $CR_{SD}$, or tDOF. The bar graphs display the difference from the baseline correction applying 6 motion parameters regression and framewise displacement censoring (all measures are thus 0 for this confound correction variant). Note that an increase in network specificity or the number of scans and a decrease in confound correlation are considered an improvement in quality (and vice-versa). All measures were computed on the somatomotor network (**see supplementary files for other connectivity analyses**). The corrections correspond to raw: no correction, FD: only FD censoring, mot6 + FD: 6 motion parameters and FD censoring, + standard: adding standardization of variance (**methods section 7**), + DVARS: adding censoring with DVARS, + mot24: using 24 motion parameters instead, + WM/CSF: adding WM/CSF nuisance regressors, + aCompCor: adding aCompCor nuisance regressors, + GSR: adding global signal regression, + highpass: adding highpass filter at 0.01Hz, + AROMA: adding ICA-AROMA.

## A)

| Order of priority | Feature description | Justification for order of priority | Data quality report to consult | Associated confound correction (listed in order of priority) | Additional considerations |
|---|---|---|---|---|---|
| 1) DVARS spikes | Spikes in the DVARS timecourse are affecting the network timecourse. | Spikes should be addressed first, as frame censoring is conducted first in the correction workflow. | Scan diagnosis: temporal features (**sup. figure 6**) | DVARS censoring | **AROMA:** ICA-AROMA is usually prioritized before the regression options, as it may yield substantial improvements and better address confound removal. The dimensionality of the ICA decomposition must be inspected, and should not be unreasonably high (we aimed for ~10-20 components per scan in this study). If the dimensionality is high, AROMA was re-attempted with 20 components.<br><br>**24 motion parameters:** This option was only considered if there was clear movement in the 6 motion parameters and noticeable signatures of motion in other quality features.<br><br>**Standardize variance:** this option was only considered if there were signatures of confounds in the $BOLD_{SD}$ map. It is also considered after regression strategies, as this correction does not remove the confound, but only reduce their potential contribution (i.e. reduce the variance of most corrupted voxels). |
| 2) Global drift in carpet plot | Slow global signal fluctuations are observed in the carpet plot. | If there are clear drifts dominating signal fluctuation, highpass is likely the best candidate correction. | Scan diagnosis: temporal features (**sup. figure 6**) | highpass | |
| 3) High-frequency spike | A spike in the frequency spectrum is observed at high frequencies. | If there is a clear spike in the high frequency spectrum, it can be best mitigated with lowpass filter. | Scan diagnosis: temporal features (**sup. figure 6**) | lowpass (not considered in this study) | |
| 4) Disturbances in global signal covariance | Non-neural sources are driving the contrast in global signal covariance (e.g. **figure 2D**). | Global signal disturbances are often the main descriptor of data quality issues which will impact various aspects of connectivity estimates. Recovering ideal contrast (e.g. **figure 2A**) is prioritized. | Scan diagnosis: spatial features (**sup. figure 6**) | 1. ICA-AROMA<br>2. aCompCor<br>3. WM/CSF<br>4. 24 motion parameters<br>5. Global signal regression<br>6. Standardize variance | |
| 5) Scan-level temporal correlation with confound | Some scans don't pass the threshold for confound correlation. | This measure best captures important spurious effects at the scan level (**sup. figure 12**). Addressing these issues first is likely to improve the interpretability of other quality control measures. | Distribution report (**sup. figure 9**) | 1. aCompCor<br>2. WM/CSF<br>3. Global signal regression | |
| 6) Scan-level network specificity | Some scans don't pass the threshold for network specificity. | Scan-level thresholds are addressed before the group statistics below to gather as many scans as possible. | Distribution report (**sup. figure 9**) | 1. aCompCor<br>2. Standardize variance | |
| 7) Group-level network variability | The network variability map is not specific to the network. | This is addressed before the group correlation, as the group correlation may be unreliable before network specificity is ensured (**sup. figure 11**). | Group statistical report (**figure 4A**) | 1. aCompCor<br>2. 24 motion parameters<br>3. Standardize variance | |
| 8) Group-level confound correlation | There are significant group-level confound correlations. | Confound correlation at the group level are inspected last. | Group statistical report (**figure 4A**) | 1. 24 motion parameters (for mean FD)<br>2. Remove outliers (e.g. **sup. figure 9**) | |

## B)

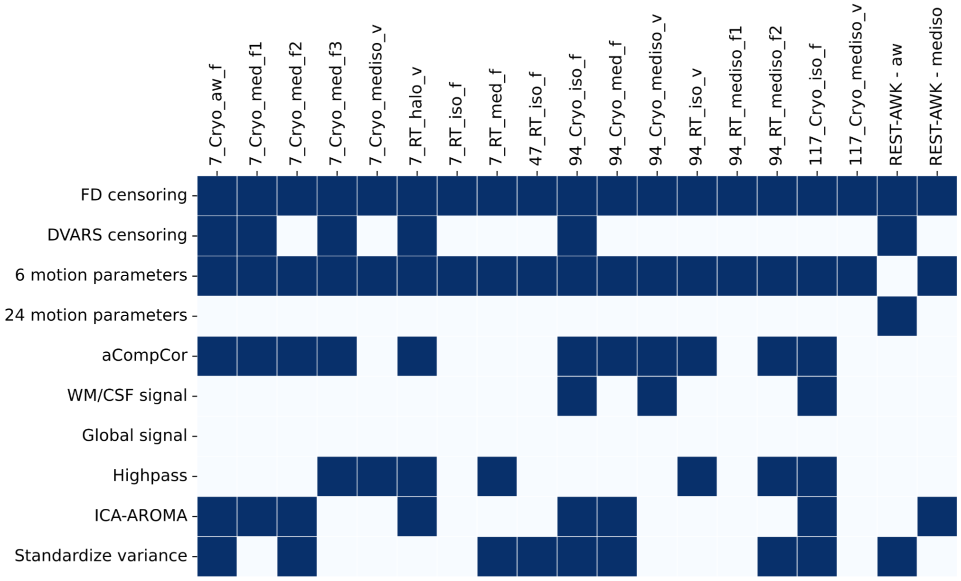


**Supplementary figure 17: A)** Supporting table for selecting and prioritizing adequate confound correction strategies. These recommendations were designed during the initial testing of all correction strategies across datasets (**sup. material section 3**). For a given correction strategy, cases which led to improvements were further investigated using the spatiotemporal diagnosis and quality control reports to identify features which may predict improvements, and the resulting recommendations are listed in table. **B)** The optimized confound correction strategy designed for each dataset.

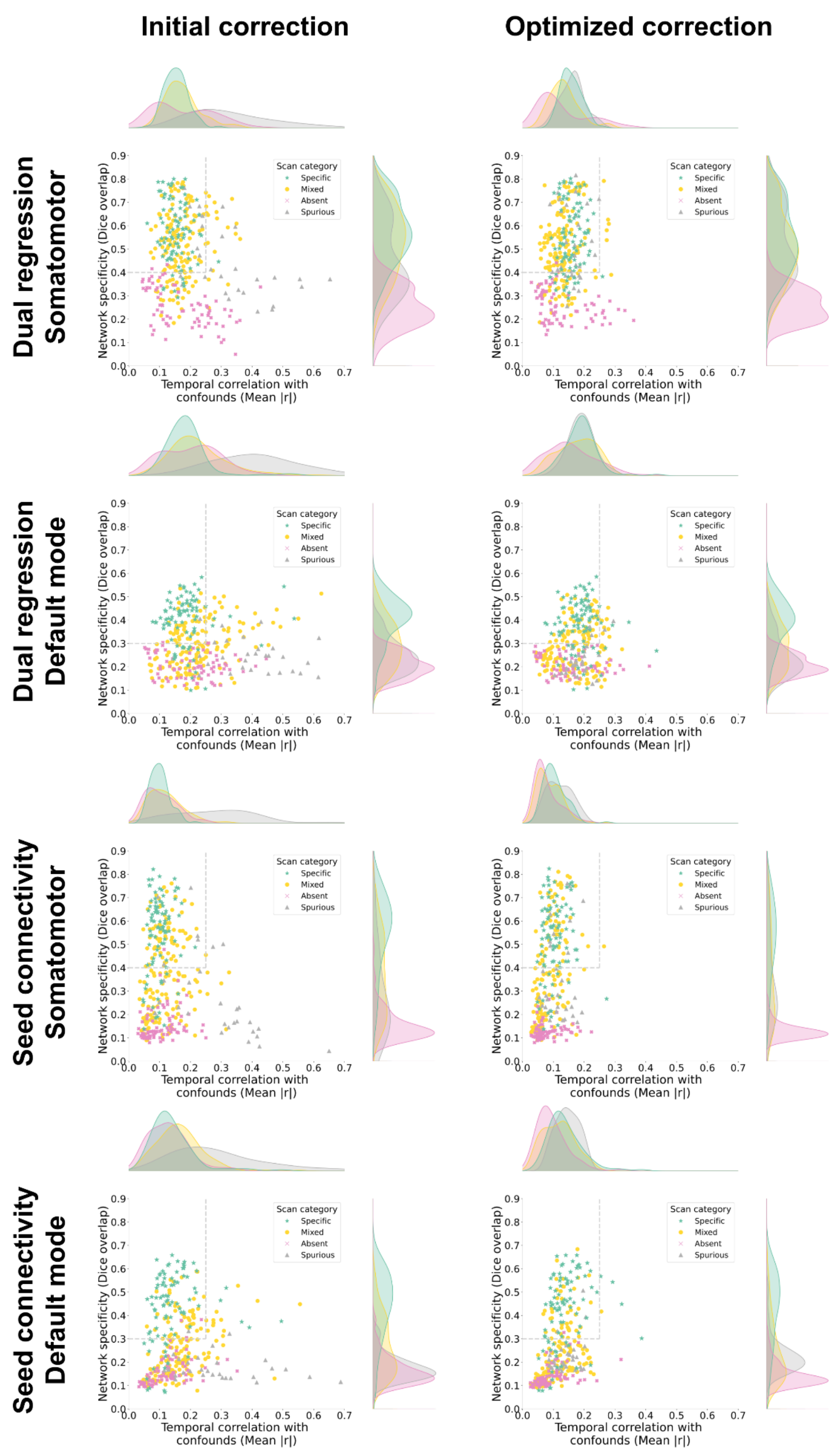


**Supplementary figure 18:** Repetition of **figure 3**, with initial and optimized confound correction, but for 4 different analyses: dual regression of the somatomotor network (found in the main text), dual regression of the default mode network, seed connectivity of the somatomotor network and seed connectivity of the default mode network. We can observe lower network specificity for the default mode network for both dual regression and seed connectivity methods. This could be a property of the evaluation of Dice overlap on this network (the threshold was lowered at 0.3 for this network instead of 0.4 based on observations from the specific category), but there was a smaller proportion of the scans expressing plausible detectability compared to the somatomotor network. Seed connectivity, compared to dual regression, had more variable outcomes across scans for network specificity. Confound effects were comparable across all analyses, with a clear reduction in those effects following confound correction optimization.

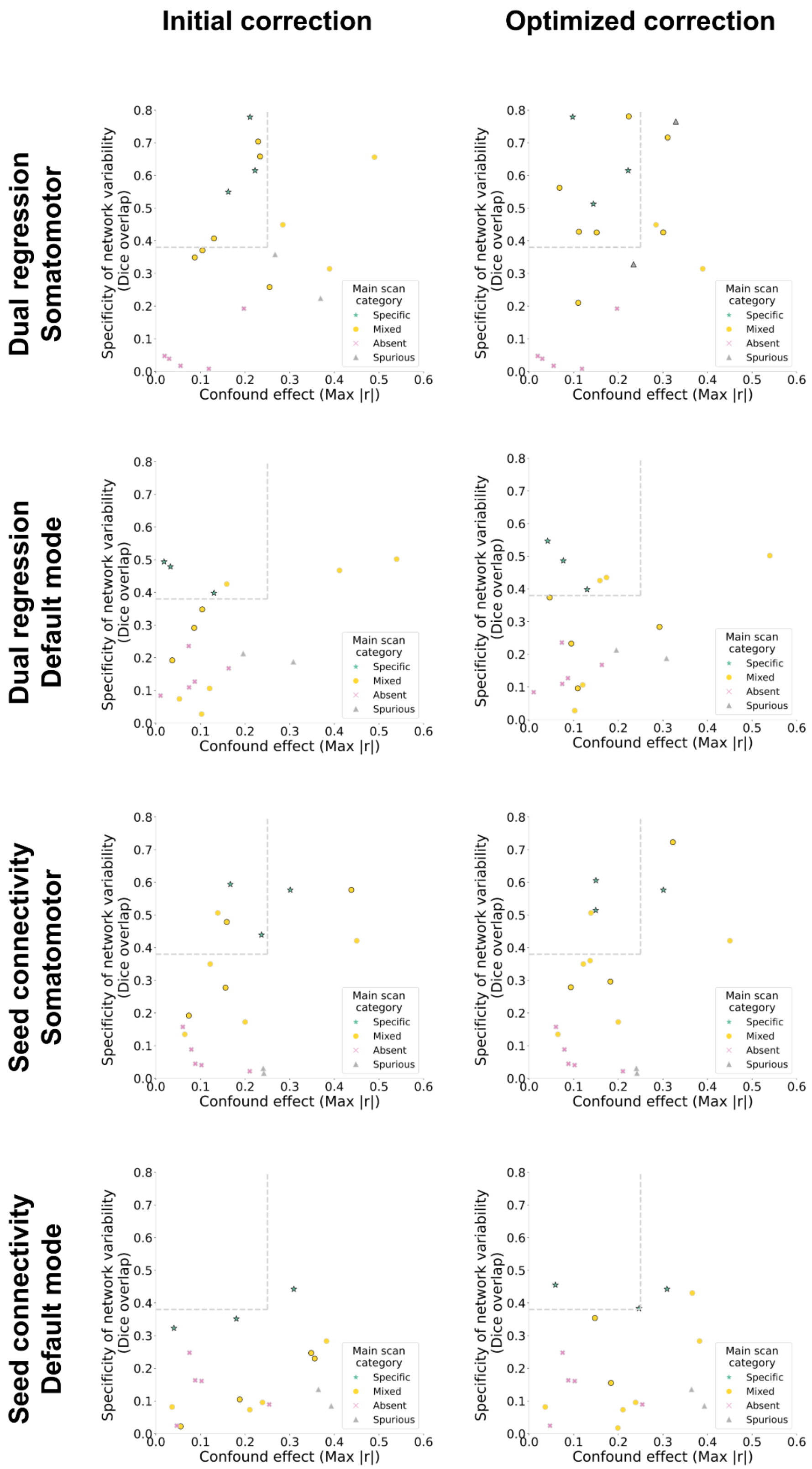


**Supplementary figure 19:** Repetition of the **figure 5**, with the summary of outcomes for the group statistical report before and after optimization of confound correction, but the four different connectivity analyses investigated (as with **sup. figure 18**). The specificity of network variability is overall lower for the default mode network, and comparatively lower for seed connectivity. In these alternative analyses, fewer datasets are meeting the ideal thresholds for quality control, primarily as a consequence of lower network variability, which may indicate a need for larger sample sizes (**sup. figure 15**).

### REFERENCES

Avants, B. B., Epstein, C. L., Grossman, M., & Gee, J. C. (2008). Symmetric diffeomorphic image registration with cross-correlation: evaluating automated labeling of elderly and neurodegenerative brain. *Medical Image Analysis*, *12*(1), 26–41.

Avants, B. B., Tustison, N., & Song, G. (2009). Advanced normalization tools (ANTS). *The Insight Journal*, *2*, 1–35.

Ciric, R., Rosen, A. F. G., Erus, G., Cieslak, M., Adebimpe, A., Cook, P. A., Bassett, D. S., Davatzikos, C., Wolf, D. H., & Satterthwaite, T. D. (2018). Mitigating head motion artifact in functional connectivity MRI. *Nature Protocols*, *13*(12), 2801–2826.

Ebisuzaki, W. (1997). A Method to Estimate the Statistical Significance of a Correlation When the Data Are Serially Correlated. *Journal of Climate*, *10*(9), 2147–2153.

Friston, K. J., Williams, S., Howard, R., Frackowiak, R. S. J., & Turner, R. (1996). Movement-Related effects in fMRI time-series. In *Magnetic Resonance in Medicine* (Vol. 35, Issue 3, pp. 346–355). https://doi.org/10.1002/mrm.1910350312

Lindquist, M. A., Geuter, S., Wager, T. D., & Caffo, B. S. (2019). Modular preprocessing pipelines can reintroduce artifacts into fMRI data. *Human Brain Mapping*, *40*(8), 2358–2376.

Manjón, J. V., Coupé, P., Martí-Bonmatí, L., Collins, D. L., & Robles, M. (2010). Adaptive non-local means denoising of MR images with spatially varying noise levels. *Journal of Magnetic Resonance Imaging: JMRI*, *31*(1), 192–203.

Marek, S., Tervo-Clemmens, B., Calabro, F. J., Montez, D. F., Kay, B. P., Hatoum, A. S., Donohue, M. R., Foran, W., Miller, R. L., Hendrickson, T. J., Malone, S. M., Kandala, S., Feczko, E., Miranda-Dominguez, O., Graham, A. M., Earl, E. A., Perrone, A. J., Cordova, M., Doyle, O., … Dosenbach, N. U. F. (2022). Reproducible brain-wide association studies require thousands of individuals. *Nature*, *603*(7902), 654–660.

Muschelli, J., Nebel, M. B., Caffo, B. S., Barber, A. D., Pekar, J. J., & Mostofsky, S. H. (2014). Reduction of motion-related artifacts in resting state fMRI using aCompCor. *NeuroImage*, *96*, 22–35.

Otsu, N. (1979). A threshold selection method from gray-level histograms. *IEEE Transactions on*

*Systems, Man, and Cybernetics*, *9*(1), 62–66.

Power, J. D., Barnes, K. A., Snyder, A. Z., Schlaggar, B. L., & Petersen, S. E. (2012). Spurious but systematic correlations in functional connectivity MRI networks arise from subject motion. *NeuroImage*, *59*(3), 2142–2154.

Power, J. D., Mitra, A., Laumann, T. O., Snyder, A. Z., Schlaggar, B. L., & Petersen, S. E. (2014). Methods to detect, characterize, and remove motion artifact in resting state fMRI. *NeuroImage*, *84*, 320–341.

Power, J. D., Plitt, M., Laumann, T. O., & Martin, A. (2017). Sources and implications of whole-brain fMRI signals in humans. *NeuroImage*, *146*, 609–625.

Pruim, R. H. R., Mennes, M., van Rooij, D., Llera, A., Buitelaar, J. K., & Beckmann, C. F. (2015). ICA-AROMA: A robust ICA-based strategy for removing motion artifacts from fMRI data. *NeuroImage*, *112*, 267–277.

Schölvinck, M. L., Maier, A., Ye, F. Q., Duyn, J. H., & Leopold, D. A. (2010). Neural basis of global resting-state fMRI activity. *Proceedings of the National Academy of Sciences of the United States of America*, *107*(22), 10238–10243.

Sled, J. G., Zijdenbos, A. P., & Evans, A. C. (1998). A nonparametric method for automatic correction of intensity nonuniformity in MRI data. *IEEE Transactions on Medical Imaging*, *17*(1), 87–97.

Wang, S., Peterson, D. J., Gatenby, J. C., Li, W., Grabowski, T. J., & Madhyastha, T. M. (2017). Evaluation of Field Map and Nonlinear Registration Methods for Correction of Susceptibility Artifacts in Diffusion MRI. *Frontiers in Neuroinformatics*, *11*, 17.
